# Opioid- and NMDA-receptor-dependent neural plasticity mediates long-term analgesia from motor cortical stimulation

**DOI:** 10.64898/2026.07.01.735554

**Authors:** Nicole Mercer Lindsay, Simon Haziza, Sean Mackey, Thomas M. Baer, Grégory Scherrer, Mark J. Schnitzer

**Affiliations:** CNC Program, Stanford University, Stanford, CA 94305; Department of Biology, Stanford University, Stanford, CA 94305; Department of Cell Biology & Physiology, The University of North Carolina at Chapel Hill, Chapel Hill, NC 27599; UNC Neuroscience Center, The University of North Carolina at Chapel Hill, Chapel Hill, NC 27599; Department of Pharmacology, The University of North Carolina at Chapel Hill, Chapel Hill, NC 27599; Department of Anesthesiology, Perioperative, and Pain Medicine, Stanford University, Stanford, CA 94304; Stanford Photonics Research Center, Stanford University, Stanford, CA 94305; Department of Applied Physics, Stanford University, Stanford, CA 94305; Department of Neurosurgery, Stanford University, Stanford, CA 94305; Howard Hughes Medical Institute, Stanford University, Stanford, CA 94305

## Abstract

Exogenous opioids that activate μ-opioid receptors (MORs) in nociceptive circuits mediate transient pain relief lasting minutes to hours but have more limited utility for treating chronic pain. By comparison, electrical or magnetic stimulation of the motor cortex can induce pain relief lasting weeks, for which the underlying mechanisms have remained unclear. Here we report an unconventional role for endogenous opioidergic signaling in the rapid induction of long-lasting analgesia from motor cortical stimulation, which triggers opioid-peptide-dependent neural plasticity in the rostral ventromedial medulla (RVM), a key node in the brain’s descending pain control pathways. To dissect the circuit and cellular bases for these effects, we created a miniaturized, millimeter-sized device allowing focal, non-invasive transcranial magnetic stimulation (TMS) of the mouse motor cortex. In mice with chronic neuropathic pain, reflexive and affective pain behaviors diminished for 1–2 weeks after one session of TMS treatment. Chemogenetic and optogenetic manipulations showed that motor cortical layer 5 pyramidal neurons with axonal projections to the RVM mediated TMS-induced pain relief. High-density electrophysiological recordings revealed that TMS treatment shifted the balance of RVM activity between pain-ON and pain-OFF neurons to a state promoting greater suppression of pain. Genetic and neuropharmacological manipulations revealed that NMDA-receptor-dependent signaling and MOR activation by endogenous opioid peptides in the RVM jointly mediate the long-lasting analgesia induced by a transient bout of TMS. Strikingly, enkephalinase inhibition in the RVM during TMS treatment enhanced the amplitude and duration of analgesia, showing that transiently boosting endogenous opioidergic signaling during TMS increases analgesia-conferring plasticity. In accord, re-analyses of data from human subjects with chronic pain support the idea that opioid administration amplifies analgesia from motor cortical TMS. Overall, our results showcase miniaturized TMS devices as versatile tools for basic and translational neuroscience and detail a hybrid, long-range neural network and NMDA- and opioid-receptor-dependent plasticity mechanism for durable pain relief. These findings point the way to mechanistically grounded, synergistic neurostimulation and drug therapies for brain diseases and disorders that jointly target neural circuit and molecular signaling pathways.

## Introduction

Hundreds of millions of people worldwide report that chronic pain disrupts their daily lives^1–3^. Extant therapies can effectively relieve acute pain but are much less successful in treating chronic pain^4–6^. Opioid drugs that activate MORs typically blunt nociception for minutes to hours, so, unfortunately, opioid administration yields only temporary relief to patients with chronic pain^7–11^. Further, prolonged opioid drug treatment can lead to harmful side effects, including drug tolerance and potential addiction^10^. Hence, clinical management of chronic pain often combines drugs suited to treat acute pain with additional medications repurposed from the treatments of other neurological disorders^12^. Yet, for many patients with chronic pain, these approaches offer only modest, short-term relief ^7,12^.

Unlike pain medications, neurostimulation methods for treating pain can induce long-lasting analgesia^13–15^. An initial discovery revealed that electrical stimulation of the human motor cortex can reduce pain induced by thalamic stroke^16^. Subsequent research found that one session of motor cortical stimulation in humans^14,16–30^ or animals^31–38^ induced prolonged analgesia lasting up to multiple weeks^39^. This durability suggests neurostimulation engages distinct biological mechanisms from those providing temporary pain relief after opioid administration. However, the neural circuit and cellular bases for the induction of long-term analgesia via neurostimulation remain poorly understood.

To address this mystery, we created an experimental platform to dissect the mechanisms by which motor cortical stimulation confers long-term relief of chronic pain. For this dissection, we aimed to leverage the powerful suite of transgenic, optogenetic, chemogenetic, genetic trapping, viral labeling and tracing, and high-density electrophysiological recording methods available for studies in mice. To this end, we examined a mouse model of chronic trigeminal neuropathic pain. In human patients, trigeminal neuropathic pain is often profoundly debilitating and resistant to conventional treatments for chronic pain^40,41^. However, some patients with the condition may receive effective pain relief from motor cortical stimulation^17,42–44^. To identify a biological basis for this phenomenon in mice, we sought means of performing focal, transcranial magnetic stimulation (TMS) of the mouse motor cortex.

Since electrical approaches to neurostimulation are usually invasive and require intracranial electrodes^16,45^, TMS, which is non-invasive, has become the most commonly used form of neurostimulation in the clinic^46,47^. However, clinical TMS devices generate magnetic fields that span cubic-centimeter-scale volumes much larger than the entire mouse brain^48,49^; this makes it infeasible to perform targeted motor cortical stimulation in mice using such instruments, as they broadly excite many areas of the mouse brain concurrently^50^. Past devices for magnetic stimulation in animals or humans have taken a wide variety of forms, ranging from microscopic coils used to stimulate individual neurons^51,52^ to macroscopic coils (0.8–70 cm width) used for non-invasive neuromodulation^53–58^. To model in mice the motor cortical excitation by TMS that patients receive clinically for pain treatment, we designed and built a miniaturized, monopolar TMS device^59^ that can target mouse neocortex with millimeter-scale precision (**Fig. 1a**). With this device, we treated mice with trigeminal neuropathic pain and took a multi-disciplinary approach to dissect the network, cellular, and molecular mechanisms of the resulting long-term analgesia. The data revealed that motor cortical stimulation had effects elsewhere in the brain that were crucial for long-lasting pain relief. Notably, TMS induced an NMDA-receptor- and MOR-dependent long-term plasticity in the rostral ventromedial medulla (RVM)^60–64,65^, a brainstem region that exerts descending control over ascending pain signals in trigeminal and spinal areas. We refer to this effect as long-term analgesic plasticity.

**Figure 1.**
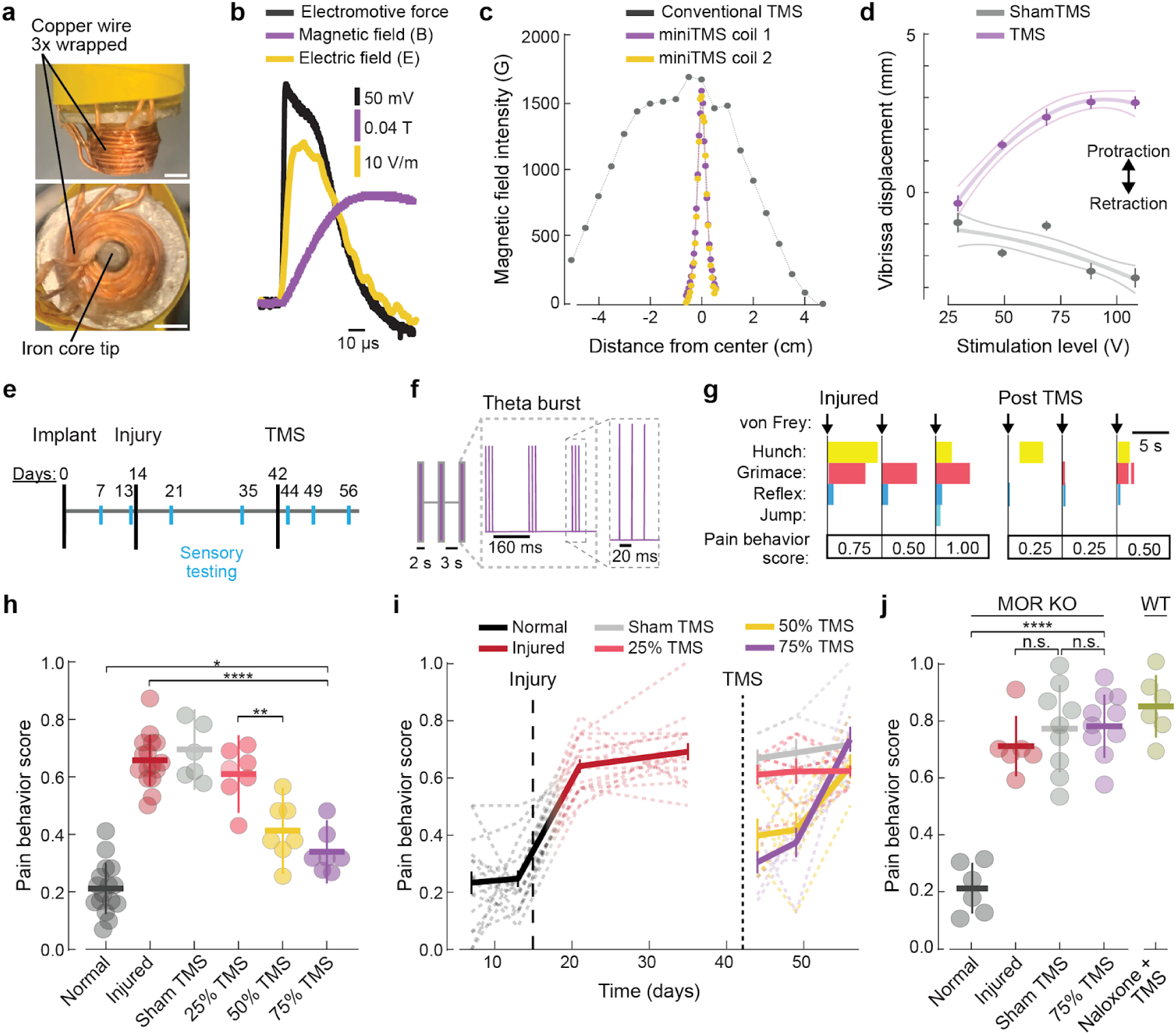
Stimulation of the motor cortex with a miniaturized TMS device yielded pain relief that lasted >1 week and required μ-opioid receptor signaling for induction. **(a)** Side- (*upper*) and bottom-view (*lower*) photographs of a miniTMS coil (3-mm-diameter iron core, sharpened to a 2-mm-diameter tip) with a 3-fold wrapping of copper wire. Scale bars: 2 mm. **(b)** We drove miniTMS devices with electromagnetic pulses, each tens of microseconds in duration. Plotted traces show for a single pulse the time-varying electromotive force (black trace; measured with a 2-mm-diameter current loop ∼1 mm from the core tip), magnetic field (purple; measured with a Hall probe axially displaced ∼1 mm from the tip of the TMS coil’s iron core), and electric field strength (yellow; calculated from the measured magnetic field). We used sequences of pulses with this waveform for all experiments in Figs. 1–5. **(c)** Cross-sectional profiles of the peak magnetic field for two different miniTMS coils (purple and yellow curves; pulses of amplitudes 15% of device maximum), showing the high similarity of the two miniaturized devices, and for a conventional figure-eight TMS coil used with human subjects (gray curve; 25% of maximum device power). Using a Hall probe, we measured the magnetic fields at axial distances of 1.5 mm and 1.5 cm, respectively, from the tip of the miniTMS and the plane of the figure-eight coils. **(d)** Vibrissae displacements evoked in an example mouse (mean ± s.e.m.; n=50 trials per datum) by a single TMS pulse applied to the motor cortex (purple) or with a sham TMS protocol in which the miniTMS coil was held 5 cm above the skull (gray) to convey to the mouse the sound but not the magnetic field from the TMS pulse, plotted as a function of the peak stimulation voltage. Thick lines: parametric fits to quadratic functions. Thin lines: ±95% CI. **Supplemental Fig. 2d** shows similar results for two additional example mice. **(e)** Timeline of experimental protocol. On Day 0, mice received a surgically implanted headplate. On Day 14, mice received a nerve injury (chronic constriction of the infraorbital branch of the trigeminal nerve). We delivered TMS treatment (see **f**) on Day 42. On intermediate days (marked in cyan), we tested the animals’ behavioral responses to tactile stimulation of the face with a von Frey filament (0.07 g, innocuous touch). After the initial bout of TMS, mice received 1 or 2 additional TMS bouts (spaced 21 days apart, not shown on the timeline) and additional testing of facial sensitivity at the same intervals as those following the first TMS bout. **(f)** Schematic of the TMS pulse protocol for intermittent theta burst stimulation (iTBS). At the start of each TMS session, we determined the minimum stimulation level needed to induce a motor response (*viz.,* the motor threshold). The amplitude of TMS treatment was then set to be a fraction (0.75, 0.5, or 0.25) of this threshold. iTBS comprised a series of 3 pulses delivered 20 ms apart, each with the waveform shown in **b**, followed by a 160-ms-interval until the start of the next set of 3 pulses. This pattern repeated for 2 s, was followed by a 3-s-interval with no stimulation, and then began again. The overall duration of stimulation was 5 min (1800 pulses in total). **(g)** Schematic showing how pain scores reflect a set of pain behaviors evoked in response to a touch to the face (black arrows) with a von Frey filament (0.07 g). Scores (ranging from 0 to 1) were based on reflexive and affective-motivational nocifensive behaviors, including hunched posture, grimacing, and jumping. To compute daily pain scores, we averaged scores evoked by 7–10 touches with the von Frey filament in each testing session. (See **Methods** and **Supplemental Fig. 3**). **(h)** TMS induced significant analgesia following a neuropathic nerve injury. Plotted are mean ± s.d. pain scores (n=6–16 mice per group) in response to a touch to the face with a 0.07 g von Frey filament. Each data point shows the mean result for an individual mouse, averaged across 2 consecutive testing sessions (see **e**), for the following conditions. Normal (dark gray, n=16 mice): pain scores measured ≥ 7 days after headplate implantation and ≤ 2 days before the nerve injury. Injured (red, n=16 mice): mean pain score, averaged over the testing sessions at 7 and 21 days after nerve injury. 25% TMS (orange, n=7 mice), 50% TMS (yellow, n=7 mice), 75% TMS (purple, n=7 mice), and Sham TMS (light gray, n=6 mice): mean pain scores, average results over testing sessions at 2 and 7 days after TMS treatment at the designated % amplitude of motor threshold or with the coil withdrawn from the head for sham treatment (ANOVA, p<0.0001; Fisher’s least significant difference (LSD) post hoc test with Holm-Bonferroni correction for multiple comparisons; *p<0.05, **p<0.01, ****p<0.0001). **(i)** Pain behavior scores of individual mice plotted as a function of time to show the changes after the nerve injury (vertical dashed line) and TMS treatment (vertical dotted line). Thin dashed lines: Data from individual mice. Thick lines: population averages. Error bars: s.e.m. (n=6–16 mice as in **h**). Analgesia lasted >7 days for stimulation at 50% or 75% of motor threshold. (Repeated measures two-way ANOVA for post-TMS results: Time, p<0.001; TMS dose, p<0.001; Time × TMS dose interaction, p=0.0015). **(j)** Induction of TMS analgesia requires endogenous opioid signaling. Plotted are mean ± s.d. pain scores, with each datum showing results from individual μ-opioid receptor knockout (MOR KO) or wildtype (WT) mice. Pain scores were measured under normal conditions (dark gray) or after nerve injury (red, averages over results at 14 and 21 days after injury), or 2 days after treatment with either sham (light gray) or real TMS (purple, 75% of motor threshold), for 6 KO mice that collectively had 18 treatment sessions (9 sham and 9 real TMS sessions; each mouse had 3 sessions spaced 4 weeks apart). WT mice received a nerve injury and then treatment combining naloxone (an opioid receptor antagonist) and TMS (olive, n=6 WT mice). ANOVA for the MOR KO groups; p<0.0001; Fisher’s LSD post hoc test with Holm-Bonferroni correction for multiple comparisons; ****p<0.0001.

First, through genetic trapping studies using activity-dependent immediate early gene labeling, we mapped at single neuron resolution the brain-wide patterns of activity evoked by TMS of the motor cortex. This revealed widespread excitation of neurons across neocortex and thalamus ipsilateral to the cortical stimulation, including prominent activation of neurons in motor cortical layers 5 and 6, which send descending axonal projections to subcortical areas. Subcortical regions implicated in pain regulation, including the RVM, also exhibited TMS-activated neurons. Today, nearly all TMS devices used clinically are bipolar, with a pair of magnetic coils that generally target human brain areas centimeters beneath the skull. Our genetic trapping results show the efficacy of an alternative approach, in which a monopolar TMS device is used to target dendrites of layer 5 cortical neurons just below the skull, activating subcortical regions receiving axons from the excited cortical cells.

Second, to map the circuitry by which motor cortical TMS induces durable analgesia, we applied viral tracing, chemogenetic, and optogenetic tools in genetically defined neuron-types. Dual genetic trapping and viral tracing studies showed that motor cortical TMS activates deep layer 5 motor cortical pyramidal cells with axonal projections in the RVM. Chemogenetic silencing of layer 5 motor cortical pyramidal cells during the administration of TMS abolished the subsequent long-term analgesia, showing that activity in this neuron class is necessary for the induction of the analgesic plasticity. Conversely, one session of selective optogenetic activation of layer 5 motor cortical cells with axonal projections in the RVM induced long-term analgesia in mice with chronic pain, showing that a suitable activation of these neurons suffices to induce durable pain relief.

Third, through high-density (Neuropixels) electrical recordings of RVM neural activity, we found that motor cortical TMS modifies the activity of one of the two key neuron classes of the descending pain control system, termed ON cells and OFF cells^61,66^. These two neuron classes were originally defined by their opponent activity patterns during nocifensive withdrawal behavior and have since been shown to promote and inhibit pain, respectively^4,61,64–66^. Opioid medications used for pain management produce analgesia in part by suppressing the spiking of ON cells and prolonging that of OFF cells^64^, but these effects are short-lasting and linked to drug presence. By contrast, our Neuropixels recordings revealed that OFF cells exhibited increases in spiking activity that outlasted the administration of TMS. This fits with the idea that motor cortical TMS induces a long-term plasticity with effects evident distal to the motor cortex, in the RVM, where the balance of RVM OFF and ON neural activity is shifted toward a more pain-suppressive state.

Fourth, to test whether these TMS-evoked changes in RVM activity reflect plasticity induced elsewhere in the brain or whether the RVM is a key induction site of long-term analgesic plasticity, we manipulated RVM activity pharmacologically or chemogenetically during TMS administration. These studies revealed that both NMDA-receptor and endogenous opioid peptide signaling in the RVM, as well as the activity of OFF cells, are indispensable for the induction of the analgesic plasticity. Conversely, we found that boosting endogenous opioid signaling via local administration of an enkephalinase inhibitor in the RVM during TMS treatment enhanced the magnitude and longevity of TMS-induced analgesia. These results show the key role of endogenous opioids in the RVM for neurostimulation-induced pain relief and provide strong evidence that engagement of opioid receptors in the descending pain control system, when paired with suitable neural excitation, can yield long-lasting analgesia—not just short-term relief. More broadly, these findings reveal a multi-modal analgesic mechanism, combining neural network activation and molecular signaling, for the induction of analgesic plasticity.

With these mechanistic discoveries in hand, we sought to gauge the potential clinical relevance of our findings. To this end, we examined whether existing clinical datasets might contain evidence for the opioidergic mediation of analgesic plasticity induction via motor cortical TMS. A re-analysis of data from patients with complex regional pain syndrome^39^ revealed that whether patients were receiving opioid medications at the time of TMS treatment associated more strongly with the improvements in pain scores a week after TMS treatment than either patient age, the duration of chronic pain prior to TMS treatment, or baseline pain levels. This finding is consistent with the hybrid network-molecular mechanism we identified in mice for analgesic plasticity and points to the potential clinical promise of pain therapies that induce analgesic plasticity by jointly engaging opioid receptors while also exciting neurons within the brain’s own pain control pathways.

Altogether, by combining insights obtained with multiple techniques, our data reveal an NMDA- and opioid-receptor-dependent plasticity in the RVM as a necessary and sufficient mechanism for the rapid induction of a long-lasting analgesia from motor cortical stimulation. Notably, TMS and simultaneous inhibition of RVM enkephalinase activity had a greater therapeutic effect than TMS alone. The hybrid mechanism that our work reveals, combining long-range network activation and neural plasticity, opens the door to a new generation of treatments for brain diseases and disorders that integrate neurostimulation with drug delivery to jointly target specific circuits as well as molecular signaling. Today, therapeutic discovery in academic and industrial contexts is usually segregated into efforts to find chemical or biologic treatments and those to develop medical devices. Our work highlights the potential of a third way, adaptive re-tuning of selected brain circuits via a synergistic combination of neurostimulation and drug-enhanced plasticity.

## Results

### Miniaturized coils for TMS of the mouse cortex

Clinical TMS apparatus typically activate cubic centimeters of human brain tissue, *i.e.*, volumes comparable to or larger than the entire mouse brain^49^. To excite specific areas of the mouse cortex at the cubic millimeter-scale, we built miniature TMS coils and accompanying power and control electronics (**Fig. 1a**; **Supplemental Fig. 1a,b**). Each coil comprises a ∼2-mm-diameter solenoid with a ferromagnetic iron core encased in wound copper wire. To generate magnetic field pulse trains of precise amplitudes and timing patterns, the driver electronics include a capacitor-discharged insulated gate bipolar transistor (IGBT) with ∼1-μs-timing accuracy, enabling brief, monophasic ∼1.5 T magnetic pulses while limiting average power consumption and heat production (**Fig. 1b**).

To characterize the operation of these ‘miniTMS’ devices, we measured the magnetic fields and electromotive forces produced by individual pulses. This confirmed our ability to shape brief pulses of electromotive force (∼50 μs duration) that were well predicted from the measured magnetic fields, as needed for temporally precise stimulation of the brain (**Fig. 1b**). We also compared the spatial extent of the magnetic fields created by the miniTMS coil to those from a conventional figure-eight coil used for clinical treatments. Magnetic fields from the latter coil extended over a ∼5 cm (FWHM) width, whereas those from the miniTMS devices had a ∼1–2 mm width, providing the spatial confinement needed for targeted stimulation of the mouse motor cortex (**Fig. 1c**; **Supplemental Fig. 1c**).

### Acute physiological responses induced by mini-TMS

To target the motor cortex for medical applications of TMS, clinicians rely on the capability of TMS pulses that are well-targeted to this brain area to evoke time-locked muscle activity, as identified via electromyographic (EMG) recordings or visual observations of a muscle twitch^67,68–70^. To apply this same targeting criterion in our studies of mice, we first sought to identify a miniTMS coil that elicited readily recordable EMG signals.

For this purpose, we empirically evaluated the magnetic pulses produced by our apparatus when used with miniTMS coils of inductance values between 7–62 µH. As expected, given fixed values of the driver current, coils of lower inductance yielded greater electromotive forces and magnetic field pulses of briefer duration (**Supplemental Fig. 1d–f**). Given this, we compared the capabilities of two different low-inductance coils (7 and 17 µH) to elicit EMG signals in the mouse vibrissa protractor muscle when used to stimulate the contralateral motor cortex. At equivalent input power levels, when centered over the orofacial portion of secondary motor cortex (area M2), the 17-µH-coil evoked substantially larger EMG responses than the 7-µH-coil, suggesting that the longer pulse duration of the former coil more effectively excited neural tissue (**Supplemental Fig. 1g, h**). Videography and machine vision analyses^71^ of the whisker movements evoked with the 17-µH-coil confirmed that individual TMS pulses elicited robust facial responses. The amplitude of whisker movements rose monotonically with the pulse amplitude, verifying the activation of physiological motor circuits (**Fig. 1d, Supplemental Fig. 2**). Based on these results, we used miniTMS coils of 17 µH for all subsequent experiments and, for each treatment session, determined the motor threshold as the lowest stimulation intensity that reliably evoked visible vibrissa responses.

### TMS of the motor cortex induces long-lasting analgesia

To examine how TMS of the motor cortex impacted chronic pain, we studied a mouse model of neuropathic orofacial pain involving the trigeminal nerve. In clinical settings, orofacial pain can be especially painful and challenging to treat, but there is evidence that it can be ameliorated by motor cortical TMS^14,17^. To model and investigate these phenomena in mice, we studied mice with a chronic constriction injury (CCI) of the infraorbital branch of the trigeminal nerve^72–75^. The injury induces a neuropathic state and hypersensitivity of the innervated orofacial area, as evidenced by nocifensive pain behaviors evoked in response to mechanical stimuli normally considered benign, a phenomenon termed tactile allodynia^76,77^.

To evaluate pain levels and tactile allodynia in mice that received the trigeminal nerve injury, we used von Frey filaments^78,79^ to stimulate the affected portion of the face (*viz.*, the nose and vibrissa pad) before and after the trigeminal nerve injury and then after TMS treatment (**Fig. 1e,f**). We then scored the mouse’s resulting nocifensive behavioral responses with a scale that evaluates both protective reflexes (*e.g.,* rapid head withdrawals) and sustained aversive responses (*e.g.,* grimacing, hunching) (**Fig. 1g, Supplemental Fig. 3; Methods**). Before the nerve injury, stimulating the face with an innocuous von Frey filament (0.07 g) rarely evoked nocifensive behaviors, but, after the injury, the same stimulus commonly triggered reflexive head withdrawal, followed by prolonged aversive responses such as grimacing (**Fig. 1g, Supplemental Fig. 3**). Given these observations, we examined whether motor cortical stimulation with a miniTMS coil would alleviate such behaviors.

Clinical TMS regimens, such as those for treating depression or orofacial pain^80,81^, often use intermittent ‘theta burst stimulation’ (iTBS), with three pulses applied 20 ms apart and a repetition of this three-pulse set at a rate of 5 Hz. An entire theta burst comprises a 2-s-interval of this stimulation pattern and is typically repeated intermittently, *e.g.*, every 10 s for a depression treatment^82^. When applied for tens of minutes, iTBS can induce prolonged changes in cortical excitability^83^.

Here, to mimic clinical protocols that induce long-lasting analgesia through brief TMS treatments, our protocol involved a 5-min duration of iTBS, organized into 2-s-bouts of stimulation separated by 3-s-intervals (**Fig. 1f**). After positioning a miniTMS coil over the portion of the right motor cortex controlling facial movement, we determined the motor threshold as the minimum stimulation level needed to evoke a facial twitch when the mouse was still. The mouse then received a 5-min bout of iTBS at an intensity of either 25%, 50%, or 75% of the motor threshold. Alternatively, mice underwent a sham stimulation protocol with the coil positioned ∼5 cm above the head, allowing the mouse to hear the sound generated during the TMS pulses without receiving the magnetic stimulation.

Notably, TMS treatment at either 50% or 75% of the motor threshold significantly reduced nocifensive behaviors in response to mechanical stimulation of the mouse’s face with an innocuous 0.07 von Frey filament (**Fig. 1h,i**). The analgesic effects of TMS were pronounced at 2 days after the treatment session and persisted for at least 7 days, resembling the duration of analgesia of a week or more in chronic pain patients treated with TMS^39^ (**Fig. 1i**, **Supplemental Fig. 4a**). The analgesia was specific to the mouse’s face, as behavioral responses to von Frey stimuli applied to the body were unaltered by TMS treatment (**Supplemental Fig. 4b**). In accord, nocifensive responses were driven by the direct contact of the von Frey filament to the face, rather than by the sight of or movement of the filament toward the mouse (**Supplemental Fig. 4c**). Female and male mice exhibited comparable levels of analgesia, with no sex-related differences detected (**Supplemental Fig. 4d**).

To assess the behavioral effects of TMS in more detail, we tracked head and limb motion using machine vision methods^71^ (**Fig. 1e, Supplemental Fig. 5a**). Net levels of spontaneous movement were comparable across cohorts of normal and injured mice as well as those that had received real or sham TMS treatment (**Supplemental Fig. 5b**). Analyses of how TMS affected individual mice revealed a modest decline in spontaneous head movements after real but not sham TMS, suggesting TMS might reduce spontaneous as well as evoked pain behaviors (**Supplemental Fig. 5c, d**).

To test more directly whether treatment with a miniTMS device can dampen acute nociceptive responses^84^, we administered real or sham TMS treatments to a separate group of uninjured mice and then 30 min later assessed hindpaw sensitivity to painful mechanical (1.0 and 2.0 g von Frey filaments) or cold (dry ice) stimuli. (For these and all subsequent experiments with mice in the paper, we delivered real TMS at 75% of motor threshold, with one exception as noted below). Mice that had received real but not sham TMS exhibited reduced levels of paw withdrawal to these painful stimuli, verifying that TMS can provide analgesia against noxious stimulation (**Supplemental Fig. 5e**).

### Induction of analgesia via TMS requires endogenous opioid signaling

Next, we examined the role of endogenous opioid signaling in the induction of analgesia from motor cortical TMS in mice with chronic pain. In healthy humans, after systemic naloxone administration (a non-selective antagonist of μ-, δ-, and κ-opioid receptors), motor cortical stimulation fails to elicit acute analgesia in response to acute painful stimuli^31^. To test whether TMS analgesia similarly requires opioidergic signaling in our mouse model of neuropathic pain, we delivered naloxone systemically prior to motor cortical TMS, which abolished TMS-induced antinociception (**Fig. 1j**).

Although it is well known that MOR mediates the analgesic properties of clinical opioid medications^5^, it has not been previously determined whether MOR mediates TMS-induced antinociception. To address this, we delivered motor cortical TMS to MOR knockout (*Oprm1^−/−^*) mice with CCI-induced neuropathic pain. After CCI, the mice showed tactile allodynia that was unaltered by TMS treatment (**Fig. 1j**), showing that MOR signaling is required for TMS-induced analgesia.

Clinically, MOR agonists are widely used as analgesics but can have harmful side effects, including analgesic tolerance and respiratory depression^85–87^. Given our finding that TMS requires MOR signaling, we examined whether tolerance develops to TMS by subjecting mice to a trio of TMS sessions spaced four weeks apart. After each TMS session, the mice had reduced pain behavior scores on days 2 and 7 post-treatment, but by day 14 their pain scores were statistically indistinguishable from those of mice that went untreated after CCI (**Supplemental Fig. 5f**). The persistence of analgesia across multiple rounds of TMS treatment suggests that the opioid receptors engaged by TMS do not desensitize, unlike the desensitization that occurs after treatment with opioid medications^88^.

We also examined whether TMS triggers respiratory depression, the main cause of fatal opioid overdoses^89^. Using whole-body plethysmography, we tracked respiratory rates in mice at baseline, after sham or TMS, or after a systemic administration of morphine that served as a positive control (**Supplemental Fig. 5g**). Although morphine strongly depressed respiratory rate, respiratory rates were indistinguishable between mice given real versus sham TMS (**Supplemental Fig. 5h**).

Overall, our data show motor cortical TMS generates MOR-mediated antinociception without respiratory depression or tolerance over repeated sessions, suggesting a low potential for abuse.

### Surveying brain-wide patterns of neural activation by motor cortical TMS

Toward identifying the brain areas mediating the long-term pain relief from motor cortical TMS, we first sought to map at cellular resolution the resulting anatomic patterns of neural activation. To this end, we performed genetic trapping studies with a Fos-TRAP2 × Ai14 mouse line^90^, which allows drug-inducible fluorescent labeling of neurons with immediate early gene (*Fos*) expression in response to electrical activation. Neurons with activity-dependent *Fos*-expression were labeled with a red marker, tdTomato, as induced via administration of the drug inducer, 4-hydroxytamoxifen (4-OHT), to mice that either received TMS or, as a control, were placed in the TMS apparatus but not treated. We then identified brain areas with greater densities of tdTomato-labeled neurons in treated versus control mice as candidate regions potentially involved in the induction or expression of long-term analgesia.

Motor cortical TMS produced substantial tdTomato-labeling of neurons in the stimulated cortical hemisphere as compared to the contralateral hemisphere (**Fig. 2a**). To quantify densities of labeled cells across the brain, we used the QUINT pipeline^91^, which allowed us to align postmortem brain sections to the Allen Mouse Brain Common Coordinate Framework (CCFv3)^92^ and to generate unbiased, atlas-wide cell density measurements^91–94^ (**Fig. 2b**; **Methods**). This revealed a characteristic, asymmetric pattern of neural activation. Namely, TMS strongly activated cortical and amygdalar regions ipsilateral to the stimulation site, plus thalamic and brainstem regions bilaterally (**Fig. 2c**). Mice that received no magnetic stimulation but heard the sounds produced by the TMS pulses lacked this pattern of neural activation, showing that it arose specifically from electromagnetic excitation of brain tissue (**Supplemental Fig. 6a**).

**Figure 2.**
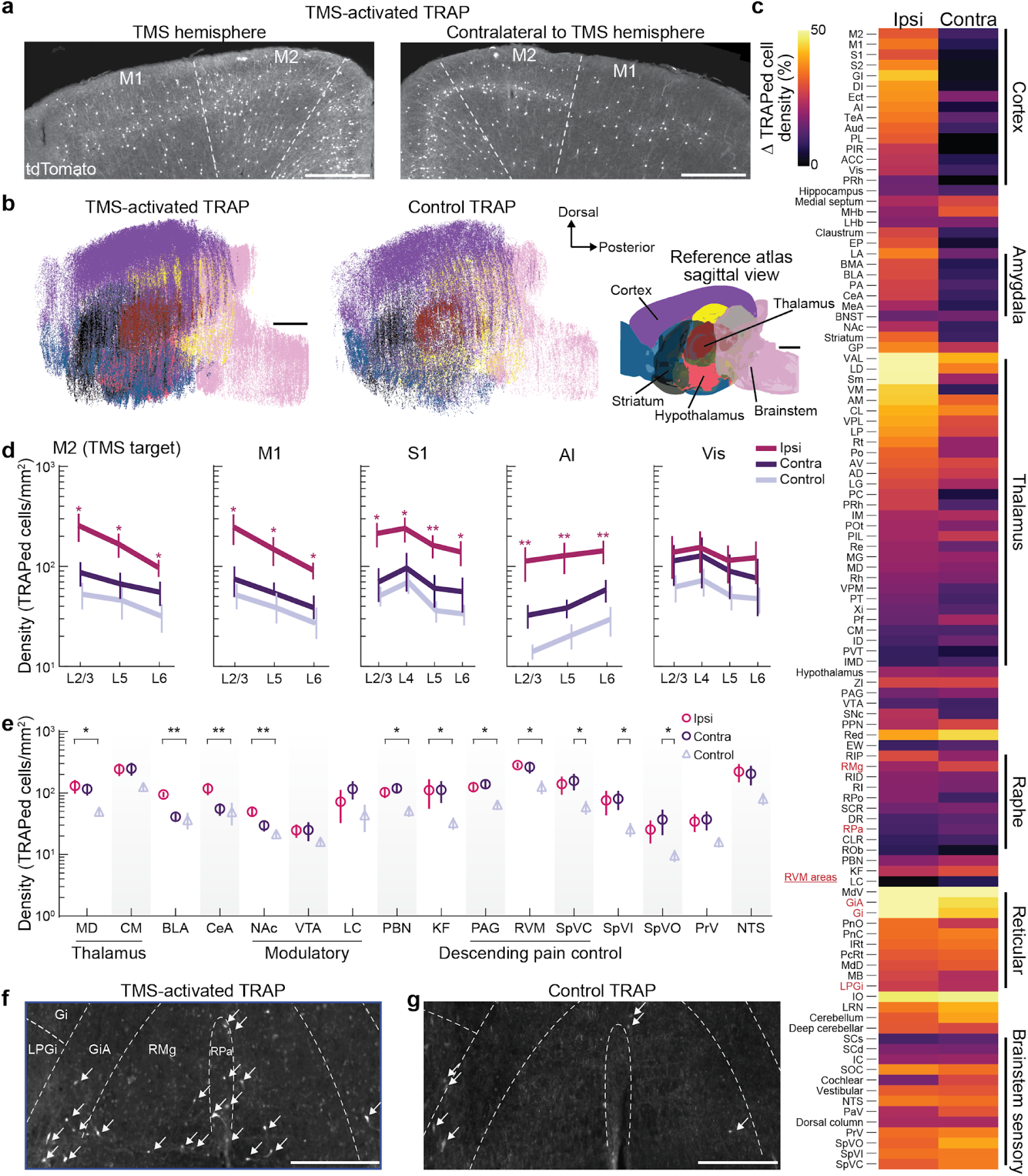
Motor cortical miniTMS activates neurons in the brain’s pain-related regions. To identify neurons activated by using a miniTMS coil to stimulate motor cortex at 75% of motor threshold, we performed genetic trapping experiments using a Fos-TRAP2 ✕ Ai14 mouse line. Neurons with TMS-evoked, activity-dependent Fos-expression were labeled with the red fluorescent marker tdTomato by delivering 4-OHT right after TMS (TMS-activated TRAP) or placement in the apparatus but no TMS delivery (Control TRAP, see **Methods**). **(a)** Example tissue sections showing TMS-activated neurons (tdTomato+) in secondary and primary motor cortices (M2 and M1, respectively) in the hemisphere ipsilateral (*left*) or contralateral (*right*) to the stimulation site in a TMS-activated TRAP mouse. Scale bars: 500 µm. All data for **a–g** were acquired using tissue perfused 7 days after the TMS session. **(b)** Two-dimensional maps showing a sagittal reference projection (*right*) and the anterior-posterior and dorsal-ventral coordinates of all tdTomato+ cell bodies in TMS-activated (*left,* n=3 mice) and control TRAP-mice (*middle*, n=3 mice). Each dot denotes an individual cell, with color indicating the brain region (Purple: Cortex, Dark blue: Olfactory areas, Yellow: Hippocampal formation, Black: Cerebral nuclei, Red: Thalamus, Green: Hypothalamus, Pink: Brainstem). Scale bars: 1 mm. **(c)** TMS activated neurons in cortical and amygdalar areas ipsilateral to the stimulation and bilaterally in many subcortical areas. The color plot shows the increase in the density of trapped neurons for areas ipsilateral (Ipsi) or contralateral (Contra) to the site of TMS, quantified as the percentage increase in the mean density of trapped cells in each of these two experimental groups relative to the mean density in the control group (n=5 hemispheres each for the Ipsi and Contra groups; n=10 control hemispheres; Kruskal-Wallis ANOVA; p<0.0001). Red font labels denote subregions of the rostral ventromedial medulla (RVM). Brain area abbreviations defined below. **Supplemental Fig. 6b** shows mean ± s.e.m. values from the same dataset. **(d)** TMS broadly activates cortical neurons ipsilateral to the stimulation. Plotted are the densities of trapped neurons, summed across individual cortical layers (L2/3, L4, L5, or L6), in secondary motor (M2, the stimulation target), primary motor (M1), primary somatosensory (S1), agranular insular (AI), and visual (Vis) cortices in ipsilateral (Ipsi), contralateral (Contra), and control hemispheres (n=5 hemispheres for Ipsi and Contra and n=10 control hemispheres; repeated measures two-way ANOVA; p-values for the effect of the stimulation condition are p=0.017 (M2), p=0.014 (M1), p=0.009 (S1), p<0.0001 (AI), p=0.35 (VIS); *p<0.05, **p<0.01 for differences between ipsilateral *vs* control hemispheres, Fisher’s LSD post hoc test with Holm-Bonferroni correction for multiple comparisons). **(e)** TMS treatment strongly activated neurons in subcortical pain centers. Plotted are mean ± s.e.m. densities of trapped neurons in ipsilateral (*Ipsi*, red), contralateral (*Contra,* purple), and control (gray) hemispheres for pain-related brain areas (Repeated measures two-way ANOVA; brain area: p<0.0001; TRAP condition: p=0.016; Fisher’s LSD post hoc test comparing Ipsi or Contra *vs.* control hemispheres with a Holm-Bonferroni correction for multiple comparisons; *p<0.05, **p<0.01, ***p<0.001). **(f, g)** Example tissue sections showing tdTomato+ cells (arrows) in subregions of the RVM in TMS-activated **(f)** and control TRAP-mice **(g)**. Scale bars: 500 µm. **Brain area abbreviations for c–g:** AD: Anterodorsal thalamus, AI: Agranular insular cortex, AM: Anteromedial thalamus, ACC: Anterior cingulate cortex, Aud: Auditory cortex, AV: Anteroventral thalamus, BLA: Basolateral amygdala, BMA: Basomedial amygdala, BNST: Bed nucleus of the stria terminalis, CeA: Central amygdala, CL: Centrolateral thalamus, CLR: Central linear raphe, CM: Centromedian thalamus, DI: Dysgranular insular cortex, DR: Dorsal raphe, Ect: Ectorhinal cortex, EP: Endopiriform, EW: Edinger-Westphal, GI: Granular insular cortex, Gi: Gigantocellular reticular formation, GiA: Gigantocellular reticular formation, alpha part, GP: Globus pallidus, IC: Inferior colliculus, ID: Interanterodorsal thalamus, IM: Interanteromedial thalamus, IMD: Intermediodorsal thalamus, IO: Inferior olive, IRt: Intermediate reticular formation, KF: Kölliker-Fuse, LA: Lateral amygdala, LC: Locus coeruleus, LD: Lateral dorsal thalamus, LHb: Lateral habenula, LG: Lateral geniculate thalamus, LP: Lateral posterior thalamus, LRN: Lateral reticular formation, LPGi: Lateral paragigantocellular reticular formation, M1: Primary motor cortex, M2: Secondary motor cortex, MB: Midbrain reticular formation, MD: Medial dorsal thalamus, MeA: Medial amygdala, MG: Medial geniculate thalamus, MHb: Medial habenula, MdD: Medullary reticular formation, dorsal, MdV: Medullary reticular formation, ventral, NAc: Nucleus accumbens, NTS: Nucleus of the solitary tract, PAG: Periaqueductal gray, PA: Posterior amygdala, PaV: Paratrigeminal, PC: Paracentral thalamus, PBN: Parabrachial, PcRt: Parvicellular reticular formation, Pf: Parafascicular thalamus, PIL: Posterior intralaminar thalamus, PIR: Piriform cortex, PL: Prelimbic cortex, PnC: Pontine reticular formation, caudal, PnO: Pontine reticular formation, oral, Po: Posterior thalamus, POt: Posterior triangular thalamus, PPN: Pedunculopontine, PRh: Perirhinal cortex, PT: Paratenial thalamus, PVT: Paraventricular thalamus, Re: Reuniens nucleus, RID: Interpeduncular raphe, RI: Interfascicular raphe, RIP: Interpositus raphe, RMg: Raphe magnus, ROb: Raphe obscurus, RPa: Raphe pallidus, RPo: Raphe pontis, Rh: Rhomboid thalamus, Rt: Reticular thalamus, S1: Primary somatosensory cortex, S2: Secondary somatosensory cortex, SCd: Superior colliculus, dorsal, SCs: Superior colliculus, superficial, SCR: Superior central raphe, Sm: Submedial thalamus, SNc: Substantia nigra compacta, SOC: Superior olivary complex, SpVC: Spinal trigeminal nucleus caudalis, SpVI: Spinal trigeminal nucleus interpolaris, SpVO: Spinal trigeminal nucleus oralis, TeA: Temporal association cortex, VAL: Ventral anterior-lateral thalamus, Vis: Visual cortex, VM: Ventral medial thalamus, VPL: Ventral posterolateral thalamus, VPM: Ventral posteromedial thalamus, VTA: Ventral tegmental area, Xi: Xiphoid thalamus, ZI: Zona incerta

Within cortex, TMS significantly activated neurons in somatomotor areas, including the primary somatosensory cortex (S1), primary and secondary motor cortices (M1 and M2), and the agranular insular (AI) and anterior cingulate (ACC) cortices (**Fig. 2c**; **Supplemental Fig. 6b**). Even though magnetic field amplitudes decline with distance from the miniTMS coil (**Supplemental Fig. 1a**)^95^, neurons in neocortical layers 5 and 6 (L5, L6) were significantly activated in multiple cortical areas, in some cases with comparable densities of trapped cells to those in layer 2/3 (L2/3) (**Fig. 2d**). This recruitment of neurons across the full depth of the neocortex suggests TMS may directly excite the apical dendrites of deep-layer pyramidal neurons. Alternatively, TMS might excite these cells indirectly via their synaptic inputs. Regardless, the densities of trapped neurons were not uniform across cortical areas; for example, in the visual cortex, the density of genetically trapped neurons was not significantly greater in mice that received TMS than in control animals. This indicates TMS preferentially engaged specific networks associated with the motor cortex, rather than indiscriminately activating areas across the cortical mantle (**Fig. 2d**).

Beyond the cortex, we examined whether TMS excites subcortical areas involved in pain (**Fig. 2e**). In the thalamus and amygdala, we found elevated densities of trapped cells in nuclei implicated in the affective and motivational components of pain, including the mediodorsal thalamus and the basolateral and central amygdala. However, in other nearby areas implicated in pain control, such as the centromedian thalamus, we did not detect elevated densities of trapped cells in mice administered TMS (**Fig. 2e**). Among neuromodulatory structures, TMS led to substantial densities of trapped cells in the nucleus accumbens but not the ventral tegmental area or locus coeruleus (**Fig. 2e**), arguing against a nonspecific arousal or global activation of neuromodulatory signals.

Most notably, TMS produced substantial densities of trapped cells across multiple areas of the descending pain control system, including the periaqueductal gray (PAG), rostral ventromedial medulla (RVM), and spinal trigeminal nucleus caudalis (SpVC) (**Fig. 2e–g**). These structures form an integrated network that gates nociceptive transmission^62,96,97^. The finding that TMS collectively excites these key areas for descending pain control gave us a likely anatomical substrate for the sustained analgesia resulting from motor cortical TMS and implicated the RVM (the central node of this pathway) in the down-regulation of pain. More broadly, the genetic trapping data showed that motor cortical TMS excites a specific brain-wide network that spans the site of cortical stimulation, deep-layer cortical neurons, and descending pain control pathways in the brainstem.

### TMS activates layer 5 pyramidal neurons that project to the RVM

To identify circuits mediating the induction of long-lasting analgesia from TMS, we examined whether this induction requires the activation of particular genetic classes of motor cortical neurons. For these studies, we selectively inhibited distinct cortical neuron populations during TMS treatment and then assessed mouse pain behaviors afterward.

To silence specific classes of motor cortical neurons, we virally expressed an inhibitory chemogenetic actuator, the DREADD hM4Di^98^, in Cre-driver mouse lines allowing targeted gene expression in cortical L2/3 or L5 pyramidal neurons or somatostatin- (SST) or parvalbumin-expressing (PV) interneurons (**Fig. 3a**). At three distinct sites spanning the orofacial representation in area M2, we injected a virus expressing hM4Di in a Cre-dependent manner (**Fig. 3b,c**). After viral expression and the trigeminal nerve injury, we treated the mice with TMS along with either clozapine-N-oxide (CNO) or saline administration. We assessed mechanical allodynia two days after TMS treatment (**Fig. 3a**).

**Figure 3.**
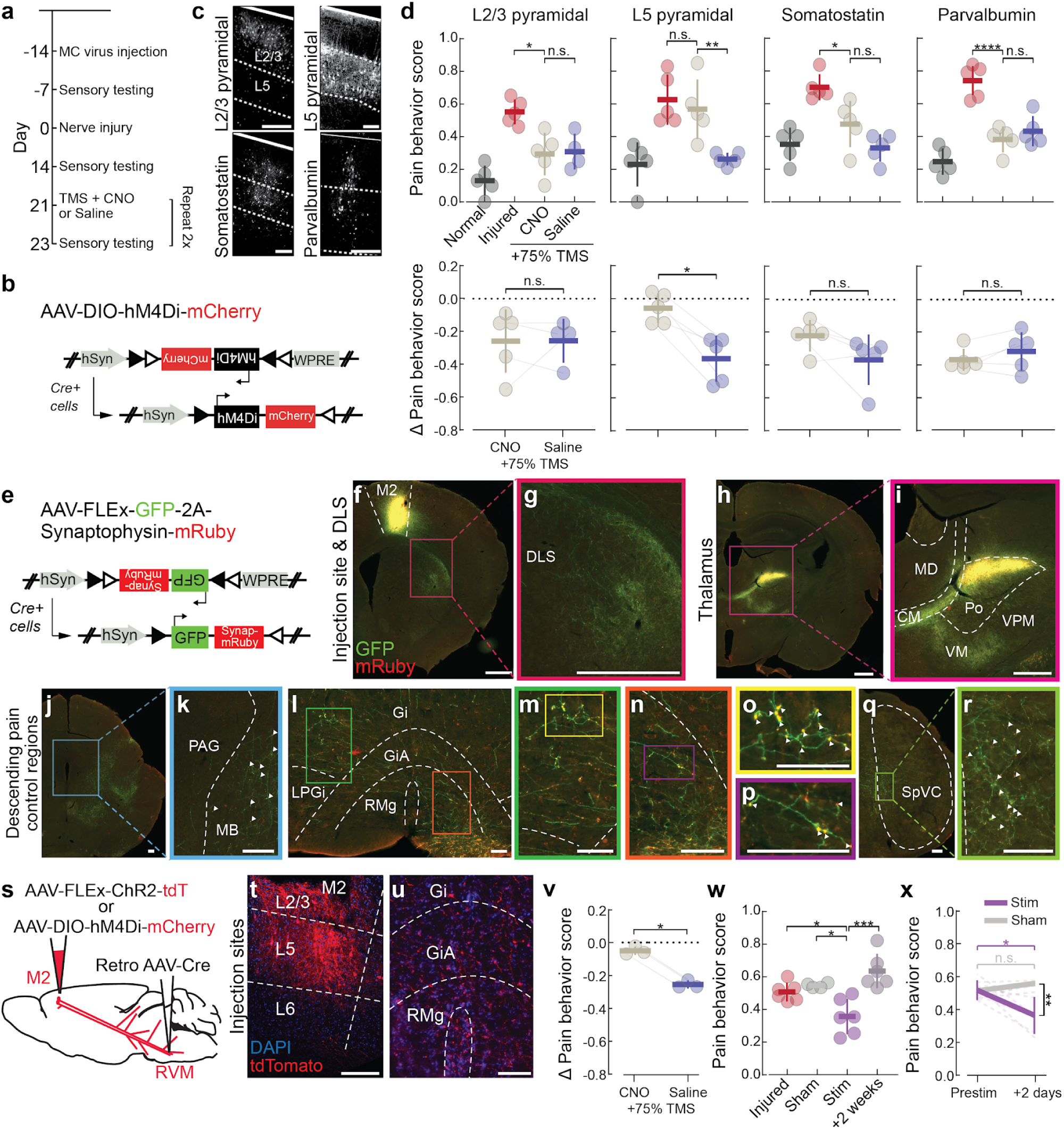
Activation of layer 5 motor cortical pyramidal cells with axons in the rostral ventromedial medulla (RVM) is necessary and sufficient to induce long-lasting analgesia via motor cortical stimulation. (**a–d**) Chemogenetic silencing of layer 5 motor cortical neurons abolished the long-lasting analgesia resulting from TMS of the motor cortex. **(a)** Timeline of experimental protocol for cell-type-specific chemogenetic silencing of cortical cell-types during TMS. We used Cre-driver mouse lines to selectively express Cre recombinase in the following cell-types: L2/3 pyramidal neurons (*Cux2^CreER^* mice), L5 pyramidal neurons (*Rbp4^Cre^*), and somatostatin and parvalbumin inhibitory interneurons (*SST^Cre^* and *PV^Cre^*, respectively). 14 days before inducing the nerve injury, we injected a virus expressing a Cre-dependent chemogenetic inhibitor hM4Di into motor cortex. 21 days after the injury, we injected mice with either clozapine N-oxide (CNO, the hM4Di activator) or a saline vehicle, followed by TMS treatment one hour later. We assessed pain behavior scores 7 days before injury, 14 days after injury, and 2 days after TMS. **(b)** Schematic of the Cre-dependent AAV-DIO*-hM4Di-mCherry* viral construct, which induced expression of the inhibitory DREADD hM4Di and mCherry in Cre-expressing neurons. **(c)** Epifluorescence images of coronal brain sections from example mice of the 4 Cre driver lines showing neural cell bodies in layers 2/3 and 5 (L2/3 and L5) that expressed mCherry. Solid white lines indicate the pial surface. Scale bars: 200 µm. **(d)** The activity of layer 5 pyramidal cells during TMS administration is required for long-lasting analgesia. *Upper*: Mean ± s.d. pain scores, with individual data points showing behavioral results from individual mice in which we chemogenetically silenced L2/3 pyramidal, L5 pyramidal, somatostatin, or parvalbumin neurons, determined before (*Normal*, dark gray) or after nerve injury (*Injured*, red) or 2 days after TMS treatment combined with administration of either CNO (light gray) or Saline (blue). ANOVA results: L2/3, L5, SST, and PV: p=0.004, 0.003, 0.005, and 0.002, respectively; *p<0.05, **p<0.01, ****p<0.0001, and n.s. mark cell-types with and without significant differences, respectively, between the CNO and either the saline or injured groups using Fisher’s LSD post hoc test with Holm-Bonferroni correction for multiple comparisons. *Lower*: Mean ± s.d. change in pain behavior scores for individual mice from prestimulus baseline to 2 days after CNO or saline injection (n=5 mice per Cre line and treatment condition; paired t-tests comparing results across individual mice; L2/3, L5, SST, and PV: p=0.7, 0.014, 0.14, and 0.25, respectively). *p<0.05; n.s.: not significant. **(e–r**) Viral tracing studies revealed motor cortical layer 5 pyramidal tract neurons that are activated by TMS have prominent axonal projections to the RVM, with additional axons observed in the dorsolateral striatum (DLS), thalamus, and spinal trigeminal nucleus caudalis (SpVC). **(e)** To perform anterograde viral tracing of neurons excited by TMS, we injected AAV-DJ-*hSyn*-FLEx-*GFP-2A-Synaptophysin-mRuby* in the motor cortex of TRAP2 mice. The mice subsequently received TMS and a 4-OHT injection to induce Cre recombinase activity in motor cortical neurons that were activated during the TMS session. In these cells, the virus expressed GFP in the cytoplasm, whereas the synaptophysin-mRuby fusion protein was predominantly localized to axon presynaptic terminals. Thus, the GFP label highlights the major targets and axon trajectories, whereas the yellow overlap indicates individual presynaptic terminals. **(f–r)** Epifluorescence images of coronal brain sections showing injection sites and downstream projections (green) and individual presynaptic terminals (yellow) using the virus of **e**. TMS-activated motor cortical neurons (injection site shown in **f**) had prominent projections to the dorsolateral striatum (DLS, **g**), thalamus (**h, i**), and the descending pain control regions RVM and spinal trigeminal nucleus caudalis (SpVC) (**j–r**) but a striking lack of axons targeting the PAG (**k**). **g**, **i**, **k**, **m–n**, **o**, **p**, and **r** respectively show magnified views of the areas within the color-corresponding rectangles in **f**, **h**, **j**, **l**, **m**, **n**, and **q**. White arrowheads in **k**, **o**, and **r** mark examples of virus-labeled presynaptic terminals. Mice were perfused ≥ 21 days after TMS + 4-OHT administration. Abbreviations: DLS: dorsolateral striatum; MD: mediodorsal thalamus; CM: centromedian thalamus; Po: posterior thalamus; VPM: ventral posteromedial thalamus; VM: ventromedial thalamus; PAG: periaqueductal gray; MB: midbrain reticular formation; Gi: gigantocellular reticular formation; GiA: alpha part of Gi; RMg: raphe magnus; RPa: raphe pallidus; LPGi: lateral paragigantocellular reticular formation; SpVC: spinal trigeminal nucleus, caudalis. Scale bars: 500 µm in **f–i** and 100 µm in **j–r**. **(s–x**) Activation of layer 5 motor cortical pyramidal cells with axons in the RVM is necessary and sufficient to induce long-lasting analgesia via motor cortical stimulation. **(s)** To selectively express channelrhodopsin (ChR2) or the inhibitory chemogenetic actuator, hM4Di, in M2 neurons that project to RVM, we used an intersectional strategy combining injections of a virus enabling Cre-dependent expression of either ChR2 **(t,u,w,x)** or hM4Di **(v)** in the motor cortex and a retrograde virus expressing Cre in RVM. **(t, u)** Epifluorescence images of coronal brain sections showing the injection sites described in **s** in motor cortex (**t**, cell bodies expressing tdT, injection site of AAV*-*FLEx*-ChR2-tdT*) and downstream axons in RVM (**u**, axons expressing tdT, injection site of Retro AAV*-Cre*). Scale bars: 200 µm. **(v)** Chemogenetic silencing of M2→RVM neurons during TMS administration showed this pathway is required for TMS-induced analgesia in mice with nerve injury. Mean ± s.d. changes in pain behavior scores for individual mice assessed at pre-stimulus baseline and 2 days after injection of either CNO or saline (n=3 mice injected as shown in **s**; paired t-test comparing results of individual mice; *p=0.013). **(w, x)** Optogenetic activation of M2→RVM neurons, targeted to express ChR2 as shown in **s**, reduced pain behaviors for ∼1 week. Mean ± s.d. pain scores, **(w**), with each data point showing results for an individual mouse after nerve injury (*Injury*, red), 2 days after sham (defocused illumination, light gray) or optogenetic (purple) stimulation, or 2 weeks after optogenetic stimulation (dark gray). (ANOVA; p<0.0001; Fisher’s LSD post hoc test with Holm-Bonferroni multiple comparisons correction, *p<0.05, ***p<0.001). Mean ± s.e.m. pain scores, (**x)**, with lines connecting the behavior scores for individual mice from pre-treatment to 2 days post-treatment (Repeated measures two-way ANOVA; Baseline *vs.* Post: p=0.159, Stim *vs.* Sham: p=0.004; Interaction: p=0.021; Fisher’s LSD post hoc test with a Holm-Bonferroni correction for multiple comparisons, *p<0.05, **p<0.01). Data are from n=6 mice that collectively underwent 10 sessions of optogenetic treatment.

After the nerve injury, mice from all four Cre-driver lines displayed robust nocifensive behaviors in response to innocuous mechanical stimulation. As expected, TMS paired with saline produced significant analgesia and reduced nocifensive behavior (**Fig. 3d**). Strikingly, chemogenetic silencing of L5 pyramidal cells during TMS abolished the analgesia, whereas silencing of either L2/3 pyramidal, SST, or PV neurons had no effect (**Fig. 3d**). These findings identify L5 pyramidal neurons as uniquely required among the cell-types we tested for TMS-induced analgesia, raising the question of how activity in these neurons is routed to downstream pain-modulatory circuits.

To address this, we next mapped the axonal projection targets of motor cortical neurons that were genetically trapped during TMS administration. We injected into the right motor cortex (M2) of Fos-TRAP2 mice an anterograde viral tracer^99,100^ expressing cytoplasmic GFP and synaptophysin-mRuby (labeling presynaptic terminals) in a Cre-recombinase-dependent manner (**Fig. 3e**). We then administered 4-OHT concurrently with a 5-min TMS treatment session.

The M2 neurons trapped during the TMS session exhibited a broad set of axonal outputs to cortical and subcortical targets (**Fig. 3f–r**), including prominent projections to brainstem pain-modulatory regions (**Fig. 3j–r**). These projections innervated classic descending pain-control regions such as RVM (**Fig. 3l–p**) and SpVC (**Fig. 3q, r**) but were largely absent in PAG (**Fig. 3j, k**), indicating that motor cortical TMS mainly regulates pain transmission via a PAG-bypassing, RVM-focused pathway. Among higher-order thalamic nuclei known to receive inputs from the stimulated portion of motor cortex^101,102^, axonal labeling was concentrated in the posterior, centromedian, and ventromedial nuclei (Po, CM, and VM), but there was little labeling in MD (**Fig. 3h,i**).

Our findings show that TMS excites layer 5 motor cortical neurons that innervate both higher-order thalamic nuclei and brainstem pain-modulatory regions, including a prominent corticobulbar pathway to the RVM. Further, motor cortical layer 5 cells play a crucial role during TMS for the induction of long-lasting analgesia. These results point to the M2→RVM pathway and the RVM as key structures mediating TMS-evoked analgesia and long-term pain relief (**Fig. 3d**).

### Excitation of the M2→RVM pathway is necessary and sufficient to stimulate long-term analgesia

We next performed a more specific examination of whether stimulation of the M2→RVM pathway is necessary and sufficient to induce analgesic plasticity. To selectively manipulate this projection, we used an intersectional viral strategy that enabled pathway-specific expression of either an inhibitory chemogenetic actuator or an excitatory optogenetic actuator. We injected a retrograde AAV expressing Cre-recombinase into the RVM of mice together with a Cre-dependent virus expressing either hM4Di or channelrhodopsin-2 (ChR2) in the motor cortex (**Fig. 3s–u**). This approach restricted expression of the hM4Di or ChR2 actuator to motor cortical neurons with axonal projections to the RVM.

To test whether the M2→RVM pathway is necessary for TMS-induced analgesia, we chemogenetically silenced these projection-specific neurons during TMS treatment of mice with trigeminal nerve injury and then assessed pain behaviors two days after stimulation, using an approach similar to that in our silencing studies of genetically defined neuron classes (**Fig. 3a**). Chemogenetic inhibition of M2→RVM neurons abolished the prolonged analgesia normally produced by TMS, indicating that this direct corticobulbar projection is required for TMS-induced pain relief (**Fig. 3v**).

We next asked whether selective activation of the M2→RVM projection sufficed to induce prolonged analgesia in the absence of TMS. To test this, we replaced TMS treatment with a 5-min bout of high-frequency 20 Hz optogenetic stimulation. Notably, optogenetic activation of this pathway significantly reduced mechanical allodynia at both 2 and 7 days post-stimulation (**Fig. 3w, x**), whereas sham stimulation with defocused light produced no change in pain scores from those measured before treatment. These findings show that activation of the M2→RVM pathway is sufficient to recapitulate the long-lasting analgesia induced by TMS.

Together, these experiments define a cortico–brainstem circuit through which TMS exerts its analgesic effects. Motor cortical stimulation recruits deep-layer 5 pyramidal neurons that directly engage the RVM, a central hub of descending pain control, and the activation of this pathway is both necessary and sufficient to produce sustained, stimulation-induced analgesia.

### TMS increases the spike rates of RVM OFF neurons during and after treatment

Having established the RVM as a potential downstream site where motor cortical TMS might exert its analgesic effects, we examined how TMS influences RVM neural activity in mice with the trigeminal nerve injury and whether TMS induces observable plasticity. Prevailing conceptions of descending pain modulation focus on the PAG→RVM circuit and RVM ON and OFF cells that bidirectionally regulate nociceptive transmission via projections to the SpVC and spinal cord dorsal horn^60,65,103^. We examined how these functionally defined RVM cell-types respond during TMS and in the tens of minutes right after stimulation.

To track population-level neural dynamics at single-cell resolution, we recorded RVM activity with high-density Neuropixels probes^104^, which allowed us to monitor hundreds of neurons during and after TMS treatment (**Fig. 4a–c, Supplemental Fig. 7**). After implanting two electromyography (EMG) wires in the intrinsic vibrissa muscle, we inserted a Neuropixels probe into the RVM. We applied a series of noxious stimuli (0.07-g and 2.0-g von Frey filaments, as well as pin prick) to the injured side of the face to evoke withdrawal responses, as monitored via the EMG recordings. This allowed us to functionally classify neurons as ON, OFF, or neutral, based on their spike rates during nocifensive withdrawal. After recording neural activity across a 5-min-baseline period, we applied a 5-min-protocol with TMS centered on brain area M2. Post-TMS recordings started 15 min after completion of the TMS protocol and lasted 5 min, allowing us to track how each of the three functionally categorized neural populations responded after TMS.

**Figure 4.**
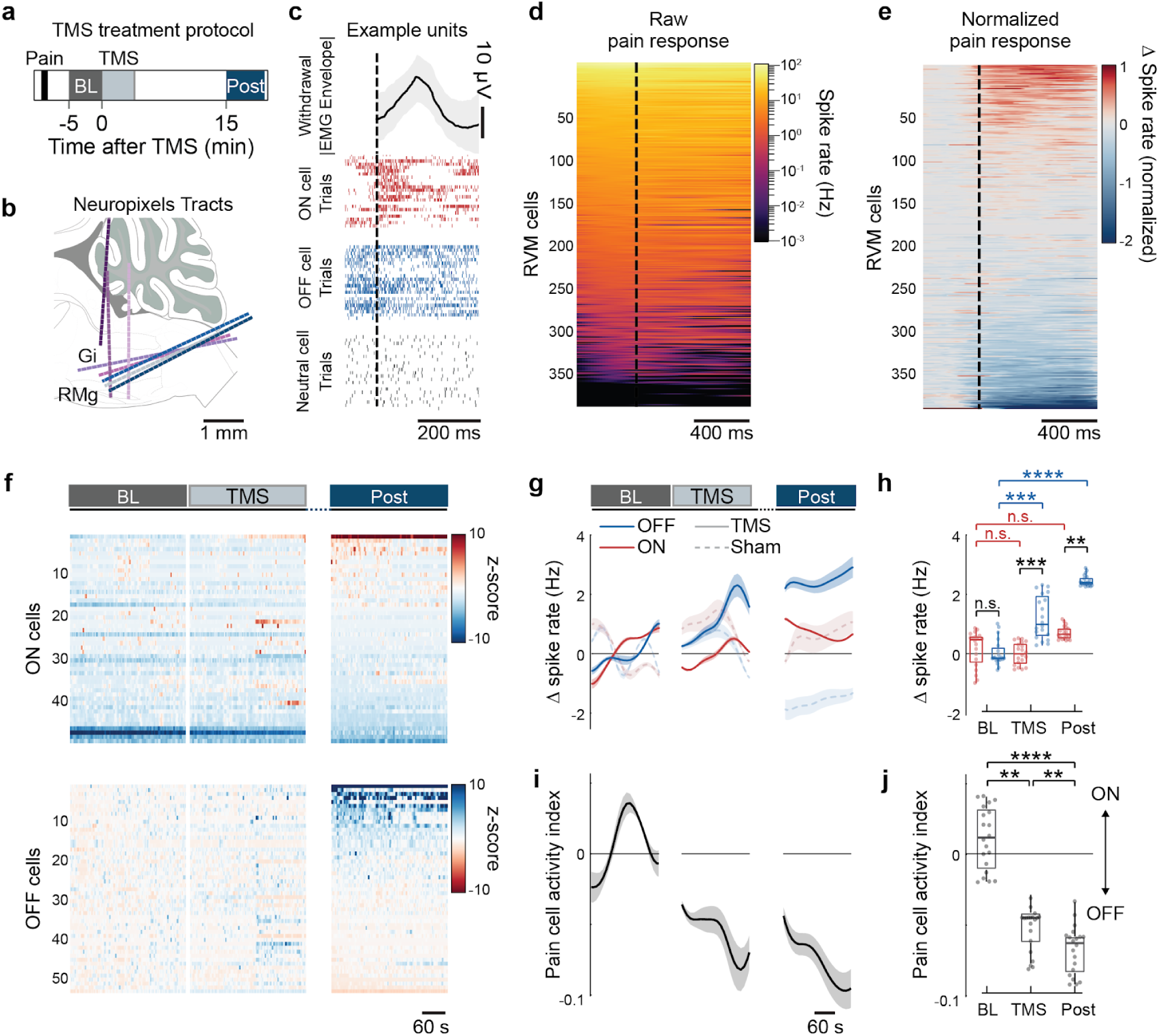
Spike rates of RVM OFF neurons increase after TMS of the motor cortex. **(a, b)** Neuropixels electrophysiological recording sessions in mice with an infraorbital nerve constriction followed the timeline of **(a)**. After implanting two electromyography (EMG) wires in the intrinsic vibrissa muscle, we inserted **(b)** a Neuropixels probe into the RVM from the foramen magnum at a 45° angle (n=5 recordings) or directly above the cerebellum (n=3 recordings). We applied a series of noxious stimuli (0.07-g and 2.0-g von Frey filaments, as well as pin prick) to the injured side of the face to evoke withdrawal responses, evaluated via the EMG recordings. After recording neural activity across a 5-min-baseline period, we applied a 5-min-protocol of TMS to the motor cortex. Measurements within a post-TMS period started 15 min after completion of TMS and lasted 5 min. See also **Methods** and **Supplemental Fig. 7.** A group of sham-treated mice received the nerve injury, had EMG wires implanted, sat in the TMS apparatus, and received noxious stimuli but not TMS. **(c)** Example responses of ON, OFF, and other (neutral) RVM neurons during pain withdrawal. *Upper*: Trial-averaged withdrawal responses (mean ± s.d. trace of the rectified EMG envelope across 20 trials). *Lower*: Spike raster plots for example ON, OFF, and neutral cells (red, blue, and gray, respectively), temporally aligned to the peak of pain withdrawal responses as determined from the EMG. Vertical dashed lines in **c–e** mark onset times of withdrawal responses. **(d, e)** We identified a total of 399 RVM neurons across 8 mice. Following the classic physiological classification of OFF and ON cells according to their activity patterns during pain withdrawal behaviors^62–64^, we categorized RVM neurons as ON, OFF, or neutral cells by comparing their dynamics during withdrawal responses to pain to those during randomly chosen intervals of equal duration in temporally shuffled datasets (p<0.05; Wilcoxon rank-sum test). Raster plots, **(d)**, show the time-varying spike rates of individual RVM neurons before, during, and after withdrawal, with the cells arranged according to their baseline spike rates. In **(e**), the rasters show the changes in spike rates relative to each cell’s baseline value, normalized to the cell’s peak increment in spiking rate. Cells in **e** are sorted by the amplitude of their spike rate changes during the 200-ms-interval immediately after withdrawal onset. **(f)** ON and OFF cell populations show divergent activity during and after TMS treatment. ON and OFF cell activity during baseline (*BL*), TMS administration, and post-TMS (*Post*) intervals. Raster plots show z-scored mean activity levels for ON (*upper*) and OFF (*lower*) neurons, arranged in the plot based on their spiking rates during the post-TMS period. Note that the color scale is reversed between the two cell-types, such that bluer tones imply less pain in both cases. **(g, h)** OFF cell spiking activity increased during and after TMS. Shown are the time-dependent, mean ± s.e.m. changes in ON and OFF neural spike rates relative to baseline levels, **(g)**, and box-and-whisker plots, **(h)**, of the distributions of spike rates during baseline (BL), TMS administration, and post-TMS intervals, in treated and sham-treated mice. Solid and dashed lines in **g** denote data from real and sham TMS sessions, respectively. Red and blue in **g** and **h** denote data from ON and OFF cells, respectively. **g–j** are based on 8 recordings in 7 mice that tracked a total of 49 ON and 53 OFF cells during TMS treatment and 73 ON and 80 OFF cells during sham TMS treatment. Repeated measures Friedman ANOVA, p<0.0001; Fisher’s LSD post hoc test with Holm-Bonferroni correction for multiple comparisons, **p<0.01, ***p<0.001, ****p<0.0001. In the box-and-whisker plots of **h, j**, horizontal lines show medians, boxes span 25th–75th percentiles, whiskers extend to 1.5 times the interquartile range, and data points show results from individual 15-s-time bins. **(i, j)** TMS of the motor cortex induced a pain-suppressing state by shifting the balance of RVM activity toward that of OFF cells. Panel **i** shows mean ± s.e.m. values of the pain cell activity index, 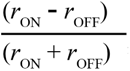, across the same time intervals used in **g**, where *r*_ON_ and *r*_OFF_ denote the mean spike rates of ON and OFF cells, respectively. Panel **j** shows box-and-whisker plots of the index values over the same three time intervals. Repeated measures Friedman ANOVA, p<0.0001; **p<0.01, ****p<0.0001, Fisher’s LSD post hoc test with Holm-Bonferroni correction for multiple comparisons.

Across 8 recording sessions, we monitored the electrical activity of 399 individual neurons across several cytoarchitectural regions of the RVM (Gi, GiA, LPGi, RMg; **Fig. 4b**). During pain-evoked withdrawal, ON cells increased their spiking activity, OFF cells suppressed spiking activity, and neutral cells showed no significant modulation (**Fig. 4c–e**), consistent with classical definitions of pain-modulatory RVM neurons. We next examined how ON and OFF neural activity evolved across baseline (BL), TMS, and post-stimulation (Post) periods (**Fig. 4f–h**). In the baseline period, ON and OFF neural spike rates were statistically indistinguishable between trials with real *vs.* sham TMS treatment (**Fig. 4g,h**). However, during TMS, OFF cells in the group of mice receiving real treatment showed a pronounced and selective increase in spiking relative to OFF cells in sham-treated mice or to ON cells (**Fig. 4g,h**). Notably, this shift in neural dynamics toward increased spiking by OFF cells persisted into the post-TMS period (**Fig. 4g,h**), showing TMS induces a sustained rise in OFF cell activity rather than a transient modulation.

To quantify how these changes affect RVM neural population dynamics, we calculated a pain cell activity index, defined as the difference between the mean spike rates of ON and OFF cells divided by the sum of the mean spike rates (**Methods**). Given this definition, negative values of the index indicate a network state favoring pain suppression. Hence, it is notable that TMS significantly reduced this index to more negative values (**Fig. 4i, j**). Moreover, this shift persisted into the post-TMS consolidation period (**Fig. 4i, j**), indicating a sustained reorganization of RVM cell population activity toward an analgesic state.

Altogether, our electrophysiological recording data show that TMS reconfigures RVM dynamics by persistently elevating the activity of OFF but not ON cells. By inducing long-lasting increases in OFF cell spiking and shifting the balance of RVM activity toward a more pain-suppressive state, motor cortical TMS produces a plastic modulation of descending pain control circuitry that favors sustained analgesia.

### Induction of analgesic plasticity requires NMDA-receptor and MOR signaling in the RVM

Having found that the activation of the M2→RVM pathway is both necessary and sufficient for the induction of stimulation-evoked analgesia (**Fig. 3s–x**), and that TMS persistently biases RVM activity toward spiking by OFF cells (**Fig. 4g–j**), we next asked which RVM mechanisms mediate the induction of long-term analgesic plasticity. Based on the broad role for NMDA receptors in neural plasticity^105^ and our specific results showing MOR signaling is required for TMS-induced analgesia (**Fig. 1j**), we examined whether the induction of long-lasting analgesia from motor cortical TMS involves MOR signaling and/or NMDA-receptor-dependent plasticity in the RVM.

First, to test whether various forms of RVM neural activity are required for TMS-induced analgesia, we performed targeted *in vivo* pharmacological manipulations of the RVM during TMS treatment (**Fig. 5a**). To examine broadly whether RVM neural activity is necessary for the induction of analgesic plasticity, we microinjected muscimol, a GABA_A_ receptor agonist, into the RVM of trigeminal nerve-injured mice immediately before TMS, to silence RVM neural activity throughout the stimulation and post-TMS consolidation periods. Although control mice that received vehicle injections exhibited the expected, major reduction in mechanically evoked nocifensive behaviors at two days after TMS, mice treated with muscimol showed no such analgesic effects (**Fig. 5b, c**). These results show that RVM neural activity is essential for the induction of long-lasting analgesic plasticity.

**Figure 5.**
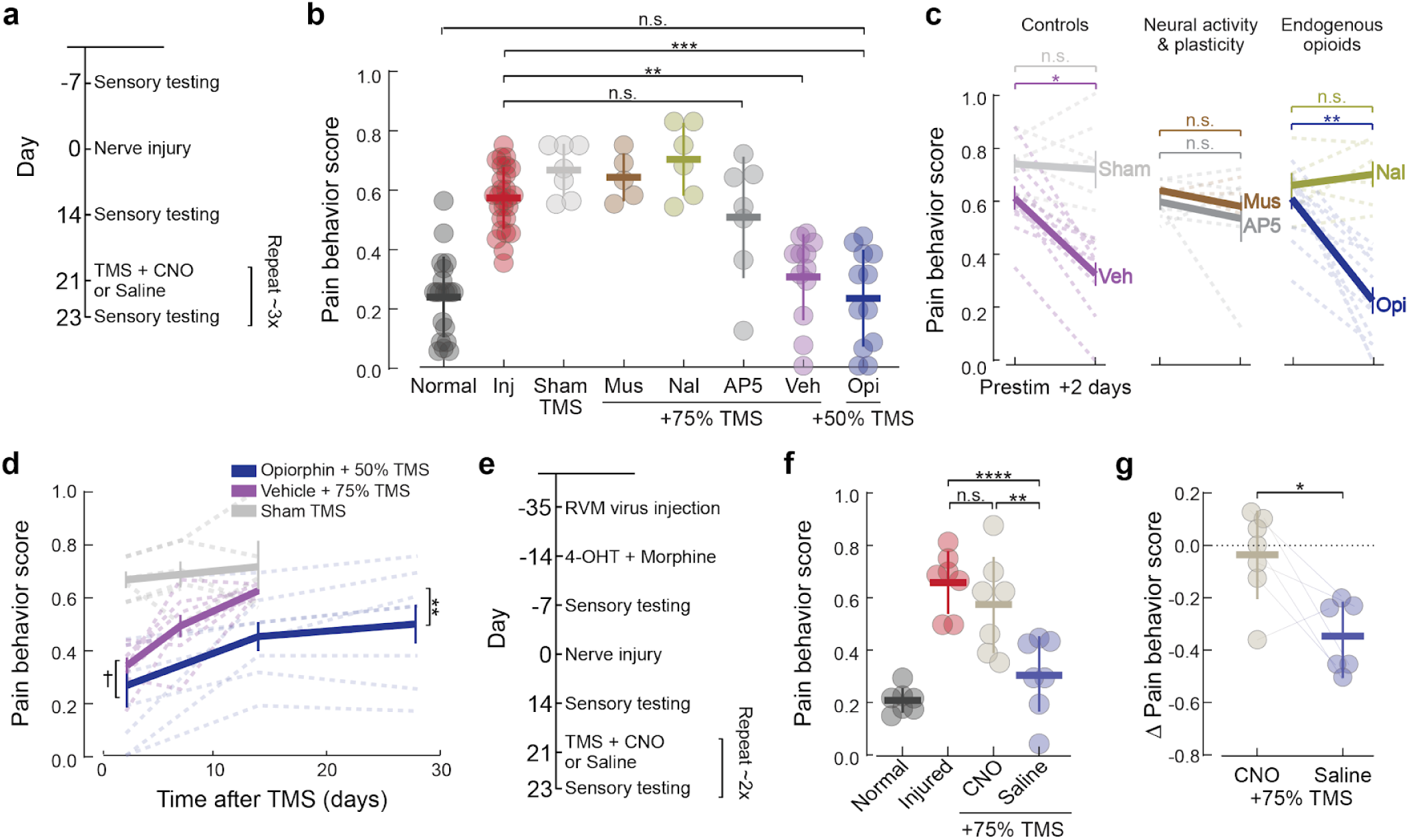
Endogenous opioid- and NMDA-receptor signaling and OFF cell activity in the RVM mediate the induction, amplitude, and duration of the long-term analgesia from motor cortical TMS. **(a–d)** Pharmacologic manipulations of RVM neural activity, opioidergic signaling, and NMDA receptor function identify the RVM as a key site of plasticity underlying TMS-induced analgesia. **(a)** Experimental timeline for pharmacologic manipulation of RVM neurons during TMS. We assessed mouse pain behavior scores 7 days before and 14 days after an orofacial nerve injury, which occurred on day 0. 21 days after the injury, we injected into the RVM either muscimol (a GABA_A_ receptor agonist that silences neural activity), naloxone (an opioid receptor antagonist), AP5 (an NMDA receptor antagonist), opiorphin^147^ (an enkephalinase inhibitor blocking the degradation of endogenous enkephalin peptides), or saline vehicle, followed by TMS treatment 5 min later. The opiorphin group received TMS at 50% of motor threshold; all others received TMS at 75% of motor threshold. Mice in the sham TMS group entered the TMS apparatus but did not receive magnetic field stimulation. 2 days after TMS treatment, we again assessed pain behavior scores. After the effects of TMS had worn off, we repeated two more rounds of TMS treatment and sensory testing. **(b)** Analgesia from TMS treatment was abolished by blocking either neural activity, opioidergic signaling, or NMDA receptors in the RVM. Plotted are mean ± s.d. pain scores (5–27 mice per group) in response to a touch to the face with a 0.07 g von Frey filament, with each datum showing results for an individual mouse for the following conditions noted in **a**. Normal (dark gray, 20 mice): pain scores at ≤ 2 days before nerve injury. Injured (red, 27 mice): pain scores at 21 days after nerve injury. Sham TMS (light gray, 6 mice), Muscimol + 75% TMS (brown, 5 mice), Naloxone + 75% TMS (olive, 6 mice), AP5 + 75% TMS (gray, 7 mice), Saline vehicle + 75% TMS (purple, 12 mice), and Opiorphin + 50% TMS (blue, 12 mice): pain scores at 2 days after TMS treatment with the designated % amplitude of motor threshold. (Kruskal-Wallis ANOVA, p<0.0001; Fisher’s least significant difference (LSD) post hoc test with Holm-Bonferroni correction for multiple comparisons; **p<0.01, ***p<0.001). **(c)** Mean ± s.e.m. pain scores measured before and 2 days after treatment, with dashed lines connecting the scores of individual nerve-injured mice that had one of the treatments in **b**, (Friedman ANOVA; p=0.029; Wilcoxon rank-sum tests with Holm-Bonferroni correction for multiple comparisons, *p<0.05, **p<0.01). Data are from n=27 mice that collectively underwent 51 treatment sessions. **(d)** Opiorphin boosted the magnitude and duration of TMS analgesia. Mean ± s.e.m. pain scores are plotted as a function of time after TMS treatment, with dashed lines showing data from individual mice. Analgesia lasted ≥ 28 days when TMS at 50% of motor threshold was combined with an RVM injection of opiorphin, versus ≈ 14 days after TMS at 75% motor threshold in control mice. (Repeated measures two-way ANOVA; time after TMS: p=0.001; drug/vehicle: p = 0.70; time ✕ drug/vehicle interaction: **p=0.002). The magnitude of TMS analgesia when opiorphin was co-administered was significantly greater than when saline vehicle was co-administered, notwithstanding that the amplitude of TMS was greater in the latter case (TMS at 75% of motor threshold for saline co-administration *vs.* 50% of motor threshold for opiorphin): †p<0.05 one-tailed paired t-test between vehicle and opiorphin treatments. Data from mice that received sham treatment are from Fig. 1i. **(e–g)** Chemogenetic silencing of RVM OFF cells diminished the long-lasting analgesia resulting from TMS of the motor cortex. **(e)** Timeline of protocol used in **f** and **g** for targeted silencing of RVM OFF cells during TMS treatment. 35 days before inducing the nerve injury, we injected into the RVM a virus expressing a Cre-dependent chemogenetic inhibitor, hM4Di (Fig. 3b). Next, at 14 days before the injury, to gain genetic access to OFF cells, we genetically trapped these neurons by systemically injecting 4-OHT and 20 mg/kg morphine; morphine disinhibits OFF cells, thus morphine-TRAP induces Cre expression selectively in OFF cells^107^. 21 days after the injury, we systemically injected mice with either saline or clozapine N-oxide (CNO, an hM4Di agonist) followed by TMS treatment at 1 hr later. We assessed pain behavior scores 7 days before the injury, 14 days after injury, and 2 days after TMS. We performed two rounds of TMS and sensory testing, with sufficient time between rounds for the analgesia to wear off. **(f)** TMS analgesia was blocked by silencing RVM OFF cells during TMS administration. Mean ± s.d. pain scores from the silencing experiment, **e**, with data points showing results from individual mice before (*Normal*) or after nerve injury (*Injury*) or 2 days after TMS administered at 75% of the motor threshold together with either CNO or saline (ANOVA; p<0.0001; Fisher’s LSD post hoc test with Holm-Bonferroni correction for multiple comparisons; **p<0.01, ****p<0.0001). Data are from 7 mice that collectively had 13 sessions of TMS. **(g)** Mean ± s.d. changes in pain behavior scores for the individual mice of **f**, evaluated at the pre-stimulus baseline and 2 days after TMS treatment co-administered with injection of either CNO or saline (n=7 mice; paired t-test comparing results across individual mice; *p=0.033).

We next asked whether the induction of TMS-evoked analgesic plasticity involves NMDA-receptor–dependent mechanisms in the RVM. Across many brain areas, NMDA receptors mediate the induction of long-term potentiation (LTP) and depression (LTD), which support sustained changes in synaptic strength^105^. To test whether such plasticity contributes to the long-lasting analgesia from motor cortical TMS, we infused the NMDA receptor antagonist AP5 into the RVM prior to TMS treatment. This blockade of NMDA receptors in the RVM fully abolished the long-term analgesia from TMS; namely, mice that received AP5 displayed comparable pain behaviors before and two days after motor cortical TMS treatment (**Fig. 5b,c**). Thus, NMDA-receptor–dependent plasticity in the RVM is required for inducing the sustained analgesia from motor cortical TMS.

Given the central role of opioidergic signaling in RVM-mediated pain modulation and our prior finding that a systemic administration of naloxone blocks TMS-induced analgesia (**Fig. 1j**), we next examined whether manipulations of opioid signaling in the RVM might alter the analgesic response to motor cortical stimulation. To test whether local opioid signaling is required for the induction of analgesic plasticity, we infused naloxone into the RVM to block μ-, δ-, and κ-opioid receptors during TMS. This manipulation abolished the analgesia from motor cortical TMS and yielded behavioral outcomes similar to those observed after TMS and either muscimol or AP5 injections into the RVM (**Fig. 5b, c**). Conversely, when we enhanced endogenous opioid signaling by infusing the dual enkephalinase inhibitor opiorphin into the RVM, mice receiving this drug treatment with TMS exhibited significantly greater pain relief than mice that received a vehicle infusion, even when the opiorphin-treated mice received a lower intensity of TMS (**Fig. 5b–d**). Remarkably, the opiorphin-treated mice displayed reduced nocifensive behaviors for up to 28 days after TMS, and the duration of analgesia approximately doubled relative to that in mice infused with vehicle (**Fig. 5d**).

Overall, our pharmacologic results show motor cortical TMS induces a previously unreported form of NMDA receptor–mediated plasticity in the RVM that converts transient brain stimulation into long-lasting analgesia. Endogenous opioid peptides gate the induction of this long-lasting plasticity, rather than simply mediating acute pain suppression. Given our results showing the sustained elevation by TMS of OFF cell activity (**Fig. 4g–j**), we next tested whether OFF cell activity in the RVM is causally required for the induction of the long-lasting analgesia.

We selectively silenced OFF cells during TMS treatment using a strategy that combined genetic trapping and chemogenetics (**Fig. 5e**). OFF cells are disinhibited following MOR activation^64,106^ and selectively express Fos after morphine administration^107^. Leveraging these past results, we injected into the RVM of Fos-TRAP2 mice a virus expressing the chemogenetic inhibitor hM4Di in a Cre-dependent manner (**Fig. 3a, b**). We then systemically administered morphine and 4-OHT to induce Cre recombination preferentially in RVM OFF cells^107^. This gave us the genetic access needed to silence the OFF cells chemogenetically during a subsequent TMS session.

Strikingly, chemogenetic inhibition of OFF cells with CNO during TMS treatment completely abolished the analgesic effects of motor cortical stimulation. CNO-treated mice with neuropathic pain showed no improvements in mechanical hypersensitivity following TMS, whereas saline-treated control mice showed greatly improved pain scores two days after TMS (**Fig. 5f, g**). These results show that OFF cell activity at or near the time of TMS treatment is required for the induction of long-term analgesia.

### Patient use of opioid medication associated with enhanced analgesia from motor cortical TMS

To explore the translational relevance of our preclinical findings, we re-analyzed published clinical data from patients with complex regional pain syndrome (CRPS) who received motor cortical TMS^39^. In the original study, TMS produced a sustained reduction in pain ratings that persisted for at least two weeks after treatment; some individuals experienced analgesia for as long as 10–14 weeks (**Fig. 6a**).

**Figure 6.**
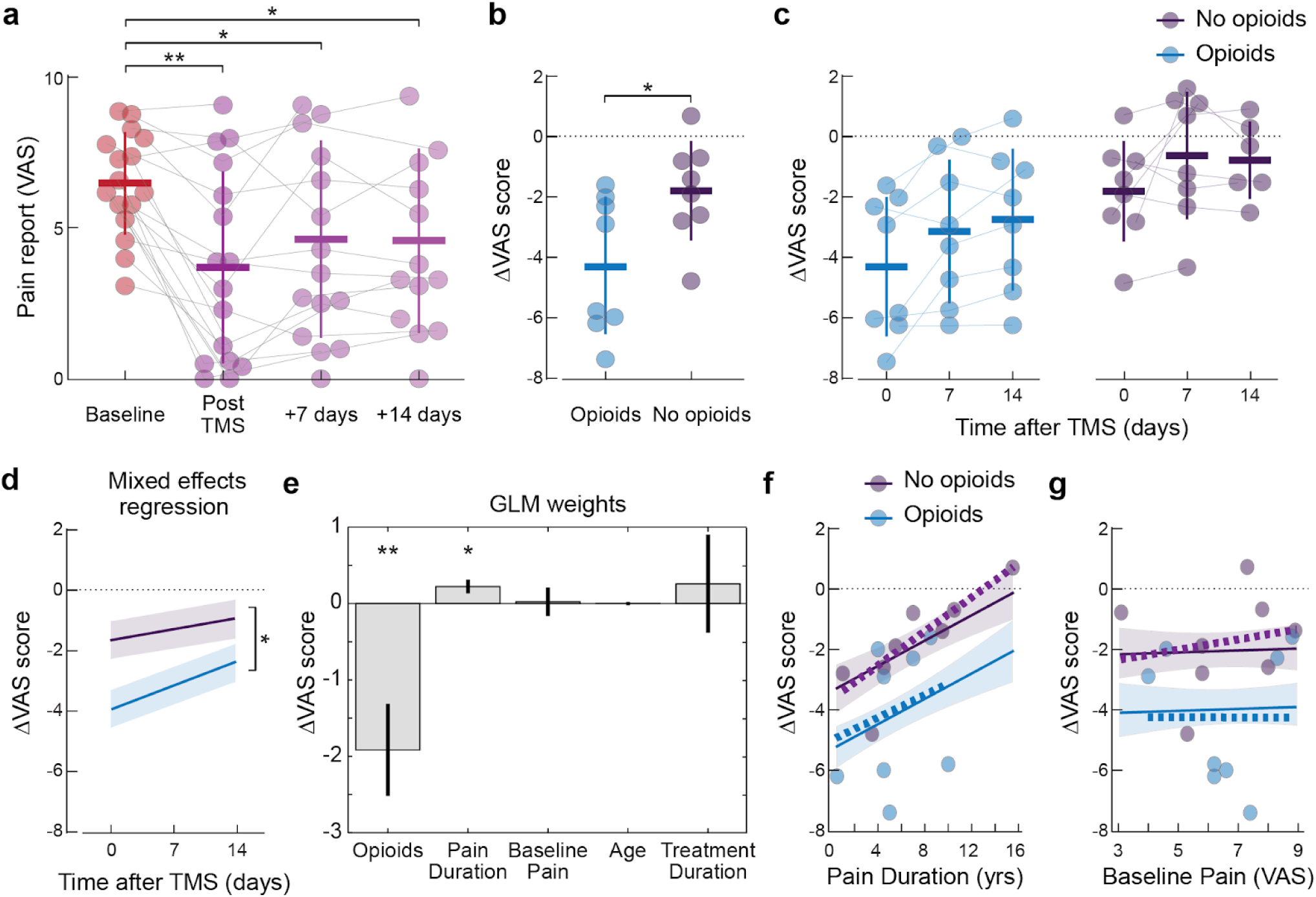
Chronic pain patients who were taking opioid medications at the time of TMS administration had greater long-term analgesic responses to TMS treatment. We re-analyzed data^39^ from patients with complex regional pain syndrome who underwent TMS treatment. Consistent with our findings in mice that MOR signaling has a key role in and can bidirectionally modulate the long-term analgesia induced by TMS (Fig. 5), the patient data revealed greater analgesic efficacy in those taking opioids at the time of TMS administration. In **a–c, f,** and **g**, each datum shows results from one of the n=16 individual patients. In **a** and **c**, thin gray lines connect results for individual patients across time points; horizontal and vertical lines denote mean ± s.d. **(a)** Mean pain intensities for each patient scored on a 0–10 visual analog scale^39^ (VAS, a measure of human pain) plotted at 4 time points (*Baseline*, over 3 days before TMS; *Post TMS*, right after the last TMS session; or *+7 days* and *+14 days*, 7 and 14 days after TMS). Colors denote periods before (red) or after (purple) treatment. Repeated measures one-way ANOVA (p<0.0001) shows significant effects of treatment (*p<0.05, **p<0.01 using Fisher’s LSD post hoc test with Holm-Bonferroni correction for multiple comparisons). **(b)** Patients taking opioid medications received greater analgesia from TMS. Plotted are patients’ decreases in pain scores (ΔVAS values) right after TMS treatment, for patients that either were (cyan) or were not (violet) taking opioid agonists at the time of TMS treatment (unpaired t-test; p=0.026). **(c, d)** Patients taking opioid medications had greater long-term analgesic responses to TMS. Plotted are, **c**, patients’ decreases in pain scores (ΔVAS values) as a function of elapsed time after TMS treatment (Repeated measures two-way ANOVA; Opioids *vs.* no opioids p=0.03; Time: p=0.003). A linear mixed effects model, **d**, yielded significantly different regression fits (shading: 66% C.I.) to the data of patients who were (cyan) or were not (violet) taking opioid agonists. (*p<0.05 for opioid *vs.* non-opioid groups), indicating that opioid use at the time of TMS treatment was associated with an additional pain score reduction of ∼1.5 points on the VAS scale over the 14-day post-treatment period. **(e)** A generalized linear model (GLM) analysis revealed predictors of TMS-induced analgesic plasticity. Bar plot shows estimated GLM coefficients ± 66% C.I. for candidate predictors (opioid use, duration of chronic pain, baseline pain, patient age, and treatment duration) of post-TMS analgesia (ΔVAS). (*p<0.05, **p<0.01 denote GLM weights significantly different from zero). GLM fits to the data are in **f, g**. **(f, g)** Opioid usage predicted increased TMS analgesia and decreased pain scores, irrespective of other factors in the GLM model. Plotted are ΔVAS scores and GLM fits (Shading: ± 66% C.I.), for pain assessments performed right after TMS treatment. Each point shows results from an individual patient, plotted as a function of, **f**, the duration of chronic pain prior to TMS treatment (p=0.02), or **g**, each patient’s baseline pain before treatment (p=0.900), for those who were (cyan) or were not (violet) taking opioids at the time of TMS treatment. For comparison to the GLM results, the thick dashed lines in **f** and **g** show the results of linear regressions that were fit independently to the data of each group, without accounting for other variables. In both graphs, the y-intercept values of the simple regressions confirm the finding in **d** that opioid use at the time of TMS treatment was associated with an additional pain score reduction of ∼1.5 points on the VAS scale over the 14-day post-treatment period.

Motivated by our findings from mice that endogenous opioid signaling regulates TMS-induced analgesic plasticity, we asked whether patients’ use of opioid medications modulated their therapeutic responses to TMS. We stratified patients based on whether they were taking opioid agonists at the time of TMS administration and found that such individuals had greater and more persistent reductions in pain scores after TMS treatment than those not taking opioids (**Fig. 6b,c**). Notably, a majority of the strongest responders to TMS were in the opioid-treated group (**Fig. 6c**), suggesting that opioid receptor signaling may indeed influence the capacity of TMS to induce durable analgesic effects.

To supplement our initial simple comparisons and regression results (**Fig. 6b,d**), we examined the relative contributions of opioid administration and other recorded clinical variables to the improvement in patients’ pain scores after TMS treatment. To do this, we created a generalized linear model (GLM) that incorporated opioid use, duration of chronic pain, baseline pain severity, patient age, and treatment duration (1 *vs.* 5 days of TMS treatment^39^) as concurrent predictors of post-TMS pain relief scores (**Fig. 6e**). This multivariate approach allowed us to assess how strongly each factor associated with improved pain scores while mitigating confounds arising from correlations across factors. Within this framework, opioid use emerged as the factor associated most strongly with enhanced TMS-induced analgesia, whereas baseline pain severity, patient age, and treatment duration contributed minimally to outcome variability (**Fig. 6e**).

Predicted reductions in patients’ pain scores from the GLM revealed that opioid-associated enhancement of TMS efficacy lasted across a range of pain durations and was most evident in patients with shorter disease histories (**Fig. 6f**). In contrast, neither baseline pain ratings nor age associated with the magnitude of analgesia once opioid use and pain duration were taken into account (**Fig. 6e, g**).

Altogether, our analysis suggests that opioid receptor signaling at the time of stimulation may facilitate the induction of long-lasting analgesia by motor cortical TMS in humans. Notably, opioid use is typically associated with negative long-term outcomes in chronic pain. However, our results suggest that endogenous or pharmacologically enhanced levels of opioidergic signaling may influence the success of neurostimulation therapies in promoting durable therapeutic plasticity, rather than a transient suppression of pain symptoms.

## Discussion

Opioid medications generally produce analgesia over the hours timescale; here we found that endogenous opioid signaling mediates the induction of an analgesia lasting weeks. How does this durable analgesia arise? The findings here establish a mechanistic framework for how motor cortical stimulation combined with endogenous opioid signaling yields durable relief of chronic pain.

By engineering a miniTMS device and applying it to mice with neuropathic pain (**Fig. 1a–i**), we identified circuit, cellular, and molecular mechanisms of TMS-induced analgesia. With this approach, we found that motor cortical TMS excites deep cortical layer pyramidal cells with descending axons in brainstem pain-control centers, most notably the RVM (**Figs. 2e, 3d–r**). This induces a plastic shift in the RVM activity state, with a sustained enhancement of OFF cell spiking, which is known to suppress ascending pain signals^108^.

Critically, rapid induction of this analgesic plasticity requires both NMDA and endogenous opioid receptor signaling in the RVM, identifying this brainstem region as a key site where a brief bout of neural excitation during TMS is transduced into persistent pain relief (**Fig. 5d**). Consistent with this hybrid network-molecular mechanism, re-analysis of data from chronic pain patients showed that current use of opioid medications associated more strongly with pain reduction from TMS than patient age, baseline pain severity, or the duration of chronic pain prior to TMS (**Fig. 6d**). This fits with the idea that opioidergic signaling modulates the analgesia from TMS in humans as in mice. However, these clinical findings are observational, retrospective, and based on a modest set of patients, and thus should not be interpreted as evidence supporting a clinical initiation or escalation of opioid therapy prior to TMS treatment. Notwithstanding, these observations from chronic pain patients do fit with our biological results and motivate larger, prospective clinical trials expressly designed to test the idea that a transient, mechanism-targeted enhancement of endogenous opioid peptide signaling could augment the amplitude and duration of analgesia from TMS.

### Layer 5 pyramidal and downstream RVM neurons mediate the durable analgesia from TMS

The specific neuron-types underlying the long-term analgesia from motor cortical stimulation were previously unidentified^28,37,58,109^. Here, we used mouse transgenic, genetic trapping, and chemogenetic techniques to identify area M2 layer 5 pyramidal cells with axonal projections to the RVM as the key cortical neuron-type. Selective excitation of motor cortical neurons projecting to the RVM sufficed to induce long-term analgesia (**Fig. 3w,x**), whereas chemogenetic silencing of M2→RVM neurons or RVM OFF cells or pharmacologic blockade of RVM neural signaling during TMS abolished the long-lasting analgesia that normally follows (**Figs. 3v, 5b,c,f,g**). Because layer 5 pyramidal cells typically send collateral axons to multiple cortical and subcortical targets^100,110^, TMS might in principle recruit additional brain areas besides the RVM that contribute to the magnitude or persistence of analgesia. The same caveat applies to our finding that optogenetic stimulation of M2 neurons with axonal projections in the RVM suffices to induce analgesia. Collectively, our findings reveal a hybrid mechanism for the rapid induction of a sustained analgesia via a combination of M2→RVM excitation and endogenous opioid signaling in the RVM.

### Endogenous MOR-mediated plasticity sustains persistent analgesia

Past work showed that levels of endogenous opioid release correlated with the extent of analgesia immediately after direct current stimulation of the motor cortex^111^ and that systemic injection of a pan-opioid-receptor blocker at the time of motor cortical TMS abolished the normally occurring analgesia^31^. However, these studies neither demonstrated a long-term plasticity involving opioid signaling nor identified a cellular and circuit basis for the analgesia.

In mice with neuropathic pain, motor cortical TMS produces an analgesia lasting about two weeks; local injection into the RVM of a pan-opioid-receptor blocker at the time of stimulation blocks this analgesia, whereas local injection of an enkephalinase inhibitor extends it to about four weeks (**Fig. 1i, 5d**). This striking, bidirectional role for opioid signaling in the RVM indicates endogenous opioid peptides both enable the long-lasting analgesia from TMS and regulate its duration (**Fig. 5b–d**). The RVM controls the strength of ascending pain signals^106,108^, consistent with the idea that a long-term reshaping of RVM activity could produce sustained pain relief. TMS-induced analgesia was absent in transgenic mice lacking MOR expression, implicating MOR-dependent induction mechanisms (**Fig. 1j**). However, we cannot exclude contributions from δ- or κ-opioid receptor signaling.

### Short- versus long-term analgesia from modulation of descending pain control circuits

Opioidergic signals in descending pain control pathways, especially the PAG→RVM→dorsal horn circuit, are known to be key mediators of the effects of centrally-acting analgesic drugs or threat- or stress-induced analgesia^112^. These effects are relatively short-lived and distinct from the long-term analgesic plasticity we found to be induced in the RVM by a bout of motor cortical stimulation.

Historically, the PAG→RVM→dorsal horn circuit was initially found to modulate pain in a series of studies in which PAG stimulation yielded up to ∼60 min of pain relief in animal models^113^. In humans, a subset of chronic pain patients that received ongoing stimulation of the PAG experienced reduced pain^114^. Opioidergic signaling in both the PAG and RVM is essential for these stimulation-induced analgesic effects^97,112^. Further, in the presence of opioid-receptor agonists, OFF cells in the RVM are disinhibited for about an hour or less, either through MOR-mediated inhibition of presynaptic inputs from the PAG or hyperpolarization of RVM ON cells^97,108^. Finally, it is interesting to note that the elicitation of such short-term analgesic effects can itself be conditioned, such as through associative threat conditioning^115,116^. However, a long-term associative memory of a conditioned stimulus that triggers a short-term analgesic response is very different from the continuous, long-term analgesic response we report here.

### Long-term modulation of descending pain control circuits in chronic pain

To date, work on long-term adaptations of the brainstem’s descending pain control circuitry has mainly focused on changes that underlie chronic pain. These changes generally develop gradually and seem to underlie the hyperalgesia and allodynia that characterize a chronic pain state^112,117^. By comparison, we found a rapidly induced change in these circuits that yields persistent analgesia, not pain sensitization.

During the development of chronic pain, both the PAG and RVM can exhibit maladaptive plasticity. For example, NMDA-receptor activation in the RVM can promote the initial facilitation of nociception in response to persistent inflammation^118^. Once established, chronic pain states appear to involve maladaptive plasticity in descending pain control areas including the RVM, where an imbalance of ON and OFF neural activity can lead to a chronic hypersensitivity^119^. Thus, past work has implicated RVM plasticity in promoting and sustaining pain, whereas, in contrast, the mechanisms we report here appear capable of muting pain for weeks.

### Past work on neural plasticity from opioid-receptor drug agonists

A core finding from our work is that a single bout of opioid receptor activation can have a long-term analgesic effect lasting weeks. This rapid induction of analgesia differs from past findings about opioidergic signaling and neural plasticity, much of which has come from research on either drug addiction or the biological processes underlying the development and maintenance of chronic pain.

Studies of brain tissue slices from the ventral tegmental area, PAG, parabrachial nucleus, or dorsal horn of the spinal cord showed that perfusion of a MOR agonist combined with *in vitro* electrical stimulation induced various forms of neural potentiation or depression lasting minutes to hours^120–124^. Since systemic opioid administration affects multiple brain areas, it might promote distinct (and in some cases opposing) forms of plasticity across the brain’s reward- and pain-related circuits^122^. However, the above studies in tissue slices did not examine behavioral correlates of neural plasticity.

Overall, while opioidergic pathways are known to have key roles in pain in the peripheral and central nervous systems and in chronic pain states, past work neither showed that acute opioid signaling can induce long-term analgesic plasticity nor defined the brain circuits and areas mediating any such effects. Thus, past results are quite distinct from our finding that, under suitable patterns of neural circuit activation, a brief bout of opioid signaling can confer a long-lasting analgesia.

### NMDA-receptor–dependent plasticity in chronic pain

In contrast to our finding of an NMDA-receptor-dependent form of long-term analgesia, past work implicates NMDA-receptor-dependent plasticity in the development and maintenance of chronic pain^125–127^. NMDA-receptor signaling in the spinal dorsal horn and supraspinal pain circuits is a key mechanism for an LTP-like sensitization promoting increased nociceptive transmission and pain^126^. Further, NMDA-receptor antagonists have been studied as treatments for neuropathic pain and complex regional pain syndrome^128^. These antagonists can rapidly reduce pain intensity, consistent with the idea that ongoing NMDA-receptor-mediated plasticity amplifies pain signaling, although their analgesic effects diminish as tolerance develops^129,130^, limiting their benefits over placebo effects^12^.

Here, we found that NMDA-receptor signaling in the RVM is required for the induction of a weeks-long analgesia. This involves the excitation of layer 5 corticobulbar neurons, their inputs to the RVM, and RVM OFF cells (**Fig. 5b,c**), which induces an NMDA-receptor–dependent plasticity that appears to re-balance the RVM activity state to one with enhanced pain–OFF cell activity (**Fig. 4**). In this view, NMDA receptors act as a molecular gate that converts a transient cortical drive into a durable reconfiguration of RVM brainstem circuits that suppresses ascending pain signals.

Our results also point to an interaction between NMDA-receptor–dependent plasticity in the RVM and endogenous opioid signaling in the induction of long-lasting analgesia (**Fig. 5a–d**). In other brain areas, NMDA-receptor activation can cooperate with neuromodulator signaling to couple Ca^2+^ entry through NMDA-receptors to biochemical cascades that promote lasting synaptic and circuit plasticity^131^. In the RVM, endogenous opioid release during motor cortical stimulation might analogously boost NMDA-dependent plasticity toward the promotion of analgesia, rather than chronic pain, through a persistent disinhibition of OFF cells.

Thus, NMDA-receptor signaling may provide a core mechanism for the plasticity induction, whereas MOR signaling may tilt the plasticity effects toward those that yield sustained analgesia. This type of three-way interaction between excitatory inputs, neuromodulatory (opioidergic) signaling, and NMDA-receptor activation offers a mechanistic explanation for how a brief bout of stimulation can produce durable analgesia. Such three-factor plasticity has been proposed to exist in other neural systems that show rapid induction of long-lasting plasticity^132–134^, albeit not with opioid signaling. Future molecular and electrophysiological studies will be needed to determine the exact mechanisms by which NMDA-receptor and MOR signaling jointly contribute to analgesic plasticity in the RVM.

### Activating a subcortical brain area via TMS with a miniature monopolar coil

Conventional TMS coils can stimulate human brain regions ∼1–2 cm below the skull surface^95^. To produce the magnetic field intensities needed to activate neurons at these depths, the coils have much larger diameters than those suitable for focal stimulation of the mouse brain^49^. To activate millimeter-wide areas in the mouse brain, we created miniTMS coils with diameters at this scale and control electronics enabling temporally precise magnetic field pulses (∼1 μs rise time) of comparable intensities (∼1.5 T) to those applied in humans (**Fig. 1a–c**; **Supplemental Fig. 1**). The coils’ small size allows them to be combined with electrical (**Fig. 4**) or fiber-optic recordings (data not shown) in the mouse brain, facilitating research on the biological mechanisms for a range of TMS therapies^135,136^.

Beyond miniaturization of the coil, our apparatus differs from typical clinical TMS devices in its use of a monopolar, rather than a bipolar, coil geometry. Neural excitation from TMS varies with the spatial profile and orientation of the electromotive forces that are induced relative to a neuron’s morphology, in particular the axon and dendritic tree^137^. The identity of the neuron-types activated by TMS has been widely debated^138–143^, and due to the geometric differences in the magnetic field patterns produced by bipolar versus monopolar coils^95^, the activated neuron classes seem likely to differ between the two magnet designs^140,143,144^. The magnetic field from a miniature monopolar coil has its highest intensity at the coil tip and declines with depth in tissue^95^, implying that superficial cortical layers, where pyramidal cells extend their apical dendritic tufts, may undergo maximal excitation.

Thus, despite the superficial penetration of their magnetic field pattern, small monopolar TMS coils may provide a convenient way to activate a subcortical target by exciting apical dendrites of layer 5 cortical pyramidal cells with axons in the chosen subcortical area. In our experiments, the key subcortical area for pain relief proved to be the RVM, but the same strategy may be widely applicable for exciting a range of clinically important subcortical targets. Although the miniTMS coils activated neurons with cell bodies in superficial and deep cortical layers (**Fig. 2d**), chemogenetic studies showed that layer 5, but not layer 2/3, pyramidal cell activity was crucial for analgesia (**Fig. 3d**). This fits with the anatomical fact that layer 2/3 pyramids commonly project axons horizontally to other cortical areas, whereas layer 5 pyramids are noted for their subcortical projections^102^.

Hence, our results motivate a re-thinking of TMS targeting strategies. Instead of using large bipolar coils for direct stimulation of deep brain areas, the identification and focal stimulation of neocortical areas with axonal projections to desired subcortical targets may allow an entirely new set of clinical applications that use one or more small monopolar coils to excite superficial cortical dendrites. Miniature monopolar coils, such as future versions of those built here, might even be permanently implanted in the cranium for chronic stimulation of apical dendrites without breaching the dura.

### Potential for new therapies that combine neurostimulation and drug adjuvants

The joint network-molecular mechanism that we uncovered for analgesia points to the potential efficacy of hybrid therapies that target a specific neural circuit together with a molecular signaling pathway to retune activity in the circuit. Here, infusion of an enkephalinase inhibitor doubled the duration of analgesia in mice to ∼4 weeks, versus ∼2 weeks when TMS was used on its own (**Fig. 5d**). Further, patients taking opioid medicines at the time of TMS treatment had enhanced analgesia that was statistically unaccounted for by other parameters (**Fig. 6b–g**). These results may pave the way to novel clinical treatments that combine neurostimulation and drug delivery.

Presently, efforts to develop new treatments are nearly cleanly divided between medical device engineering and molecular medicine. The gains we achieved by combining drug and neurostimulation therapies suggest clinical neuroscientists should explore a third treatment paradigm, in which one adjusts the dynamics of a chosen brain circuit by eliciting a drug-enhanced plasticity as the circuit is transiently activated through neurostimulation.

Treatments from this third paradigm would follow an intersectional logic. The network to be re-tuned would be briefly excited using a stimulation protocol sufficient to induce long-term plasticity. The magnitude, duration, and possibly the sign of this plasticity would be enhanced by a drug adjuvant to increase plasticity at the synapses of interest. A few past studies explored the capability of a partial NMDA-receptor agonist to enhance effects of transcranial direct current^145,146^ or neuromodulation treatment for major depression^135^. However, in many cases, the drug adjuvant might boost neuromodulator signaling, based on not just our work but also growing evidence for neuromodulator enhancements of long-term neural plasticity^132–134^. Note that the use of neurostimulation is critical to the treatment strategy, as it not only specifies the circuit but also creates the activity patterns needed to gate the plasticity induction. Overall, the development of hybrid drug and device treatments might unlock unique therapeutic benefits that neither individual treatment category on its own could confer.

## Acknowledgements

We thank William McCallum, Nicole Ochandarena, Fatih Dinç, Michelle Redinbaugh, Adeeti Aggarwal, Daniel Berg, Chong Cheng, Adrien Tassou, and Joshua Blair for helpful discussions. This work was supported by NIH grants F32DE03003 (N.M.L.), K99DE031802 (N.M.L.), R01NS106301 (G.S.), R01DA054583 (G.S. and M.J.S.), and R01NS124590 (M.J.S.), a Department of Defense Chronic Pain Management Research Program award HT9425-23-1-0634 (G.S. and M.J.S.), a Department of Defense Vannevar Bush Faculty Fellow award (M.J.S.), the New York Stem Cell Foundation (G.S.), and a Stanford Wu Tsai NeuroTranslate award (S.M., T.M.B., M.J.S., G.S., N.M.L.). This project was also supported in part by award 1S10OD010580-01A1 from the National Center for Research Resources (NCRR). The Stanford Cell Sciences Imaging Facility (RRID:SCR_017787) provided training and user support.

## Author Contributions

N.M.L., S.H., T.M.B., G.S., and M.J.S. designed experiments. N.M.L. and M.J.S. designed data analyses. M.J.S., S.M., and N.M.L. designed the *post hoc* analyses of patient data. T.M.B. designed and built the miniaturized TMS devices, and N.M.L., S.H., and T.M.B. empirically characterized them. N.M.L. and S.H. built the Neuropixels electrophysiology rig. N.M.L. performed all biological experiments and analyzed the mouse behavioral, histological, and electrophysiological data. N.M.L. and M.J.S. wrote the manuscript. All authors edited the manuscript.

---

**Supplemental Figure 1.**
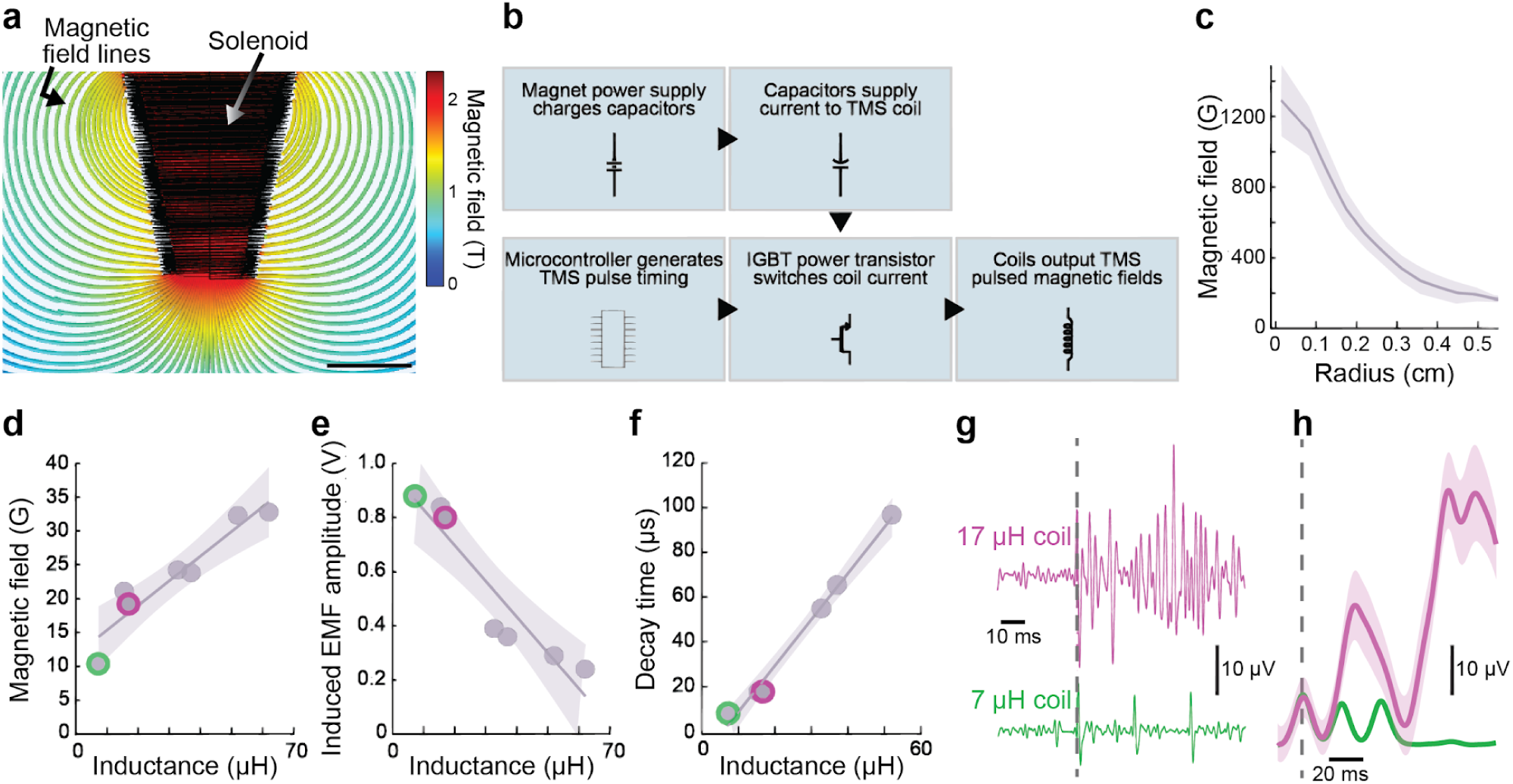
Design and testing of miniaturized TMS coils. **(a)** Map of the magnetic field lines (colored lines) generated by a miniTMS coil with three layers of 0.4-mm-diameter Cu wire wound around a powder-iron core (relative permeability of 75, driven at 100 A), as computed with the COMSOL finite-element physics simulation package. Colors indicate magnetic field amplitudes. Scale bar: 2 mm. **(b)** Basic schematic of the driver electronics for the miniTMS apparatus. A power supply (100 V; 2 A) charges a capacitor bank while a microcontroller generates precisely timed trigger pulses. The supply current from the capacitor bank and the trigger pulses converge on an insulated-gate bipolar transistor (IGBT) switch that delivers monophasic, square current pulses to the coil, creating the rapidly changing magnetic field values needed to depolarize neurons. Compared to traditional TMS coils that have an air-core, our coils substantially reduce the challenges of heat production, owing to the concentration of the magnetic field in the iron core and the use of brief (∼50 µs) pulses (Fig. 1b). **(c)** Mean amplitude of the magnetic field from a miniTMS coil, plotted as a function of the radial distance from the coil axis, as measured with a Hall probe positioned axially ∼1.0 mm beneath the core tip. Shading: s.e.m. across sampled grid points within each radial bin (n=4–21 points per bin). **(d–f)** Electromagnetic properties of miniTMS coils with inductances between 7–62 µH. Plotted are the **(d)** magnetic field strength (measured with a Hall probe axially displaced ∼1.0 mm from the tip of each coil), **(e)** induced electromotive force (EMF, measured with a 2-mm-diameter current loop axially displaced ∼1.0 mm from the coil tip), and **(f)** EMF decay time (measured from coil current decay, recorded with a 0.1 Ω current-sensing resistor in series with the coil), shown as functions of coil inductance (measured with an electronic inductance meter). To vary the coil inductance value, we changed the number of wire wraps (2 *vs.* 3) and the winding geometry around the iron core. Data points encircled in green or pink indicate the two coils for which EMG traces are shown in **g**. Solid lines: Linear regression fits (Shading: ± 95% C.I.). **(g, h)** Example raw EMG traces (bandpass-filtered between 10 Hz–20 kHz by the EMG amplifier), **g**, and trial-averaged traces of the EMG envelope (n=45 trials; bandpass filtered (100–500 Hz), rectified, and then low-pass filtered (50 Hz)), **h**, as recorded in the vibrissa protractor muscle. Traces are plotted as functions of time relative to delivery of the TMS pulse (dashed vertical line) to the contralateral motor cortex (**Methods**). TMS pulses delivered in triplets (20-ms-interpulse interval) were delivered at pseudorandomly chosen times with inter-pulse-intervals of 30–50 ms. We used a miniTMS coil with either a 17 µH (magenta traces) or a 7 µH (green traces) inductance. Based on the greater EMG signals elicited by the 17-µH-coil, we used coils of this design and inductance value for all subsequent experiments described in this paper. Shading in **h**: s.e.m. over 45 trials. (Note that s.e.m. values are too small to see on the lower trace).

**Supplemental Figure 2.**
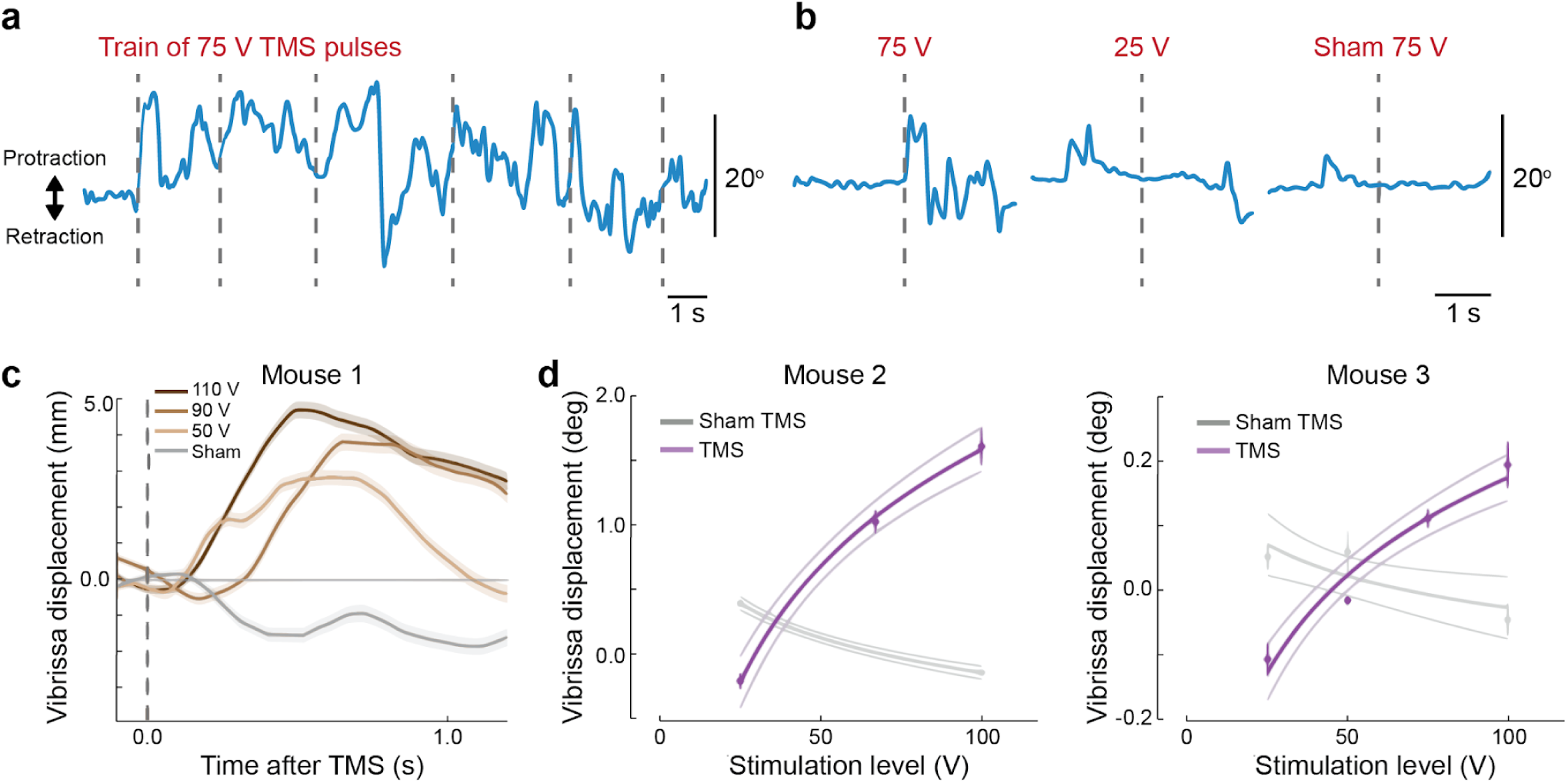
Vibrissa movement increases with the intensity of TMS. **(a, b)** Traces of vibrissa angle relative to the mouse’s head, as determined by machine vision analyses of behavioral videos taken during stimulation of the contralateral motor cortex with a 17-µH miniTMS coil. **(a)** Example trace of vibrissa angle during a series of TMS pulses delivered with a drive voltage of 75 V. **(b)** Traces showing vibrissa angle in response to TMS pulses with drive voltages of either 75 V or 25 V. In the sham (75 V) stimulation condition, the miniTMS coil was held ∼5 cm above the mouse’s head. Dashed vertical lines mark times at which TMS pulses were applied to the motor cortex. **(c)** Displacements of the vibrissa base for an example mouse (mean ± s.e.m.; n=50 trials per stimulation intensity) evoked by single TMS pulses delivered with drive voltages of 110 V (dark brown), 90 V (light brown), or 50 V (peach) TMS, or by sham stimulation with a 50 V drive voltage (gray, TMS coil positioned ∼5 cm above the head to maintain the auditory stimulus associated with a TMS pulse while precluding effective magnetic stimulation of the brain). **(d)** Vibrissa displacements evoked by a single pulse of real or sham TMS, plotted as a function of peak stimulation voltage (mean ± s.e.m.; n=50 trials per datum). The graph format, parametric fits, and experimental details are the same as those in Fig. 1d.

**Supplemental Figure 3.**
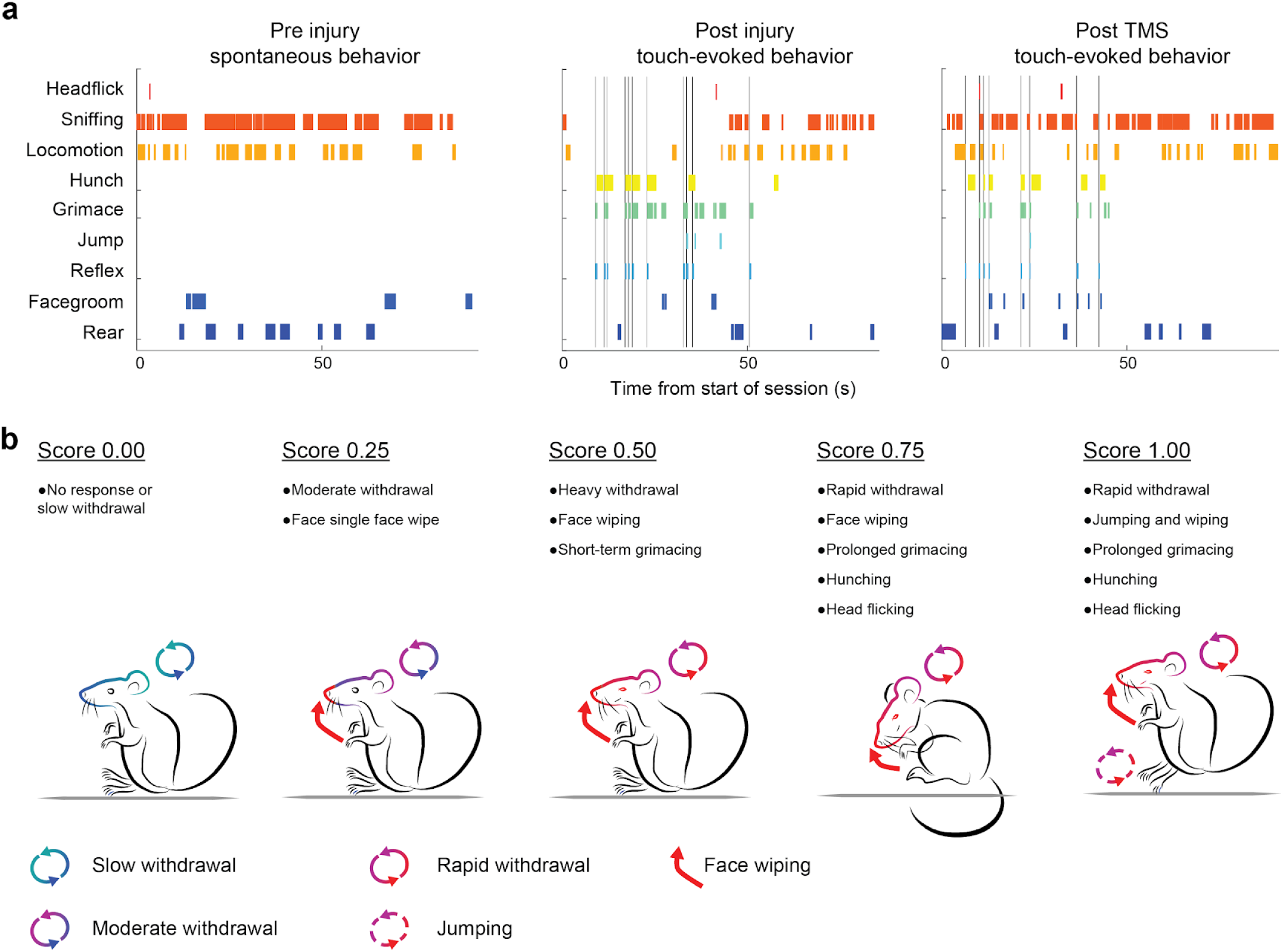
Numeric grading of mouse nocifensive behaviors in response to mechanical stimuli applied to the face. **(a)** Example raster plots showing results obtained through visual scoring of mouse behavior, revealing both pain-related (head flick, hunching, grimace, jump, reflex, and facial grooming) and healthy behaviors (sniffing, locomotion, and rearing). Each plot shows data from the same individual mouse engaged in spontaneous behavior during an initial, pre-injury testing session (*left*), during sensory testing after the facial nerve injury (*middle*), and 2 days after TMS treatment applied at 75% of the motor threshold (*right*). Vertical lines in the middle and right panels mark times at which the von Frey filament was applied to the mouse’s face. **(b)** Schematic illustration of the scoring method used to grade nocifensive responses and orofacial pain. Each touch-evoked response was assigned a score between 0 and 1 based on the number, duration, and type of nocifensive behaviors. Colored ellipses indicate the relative velocity of withdrawal or jumping. Red arrows and facial expressions depict face wiping and grimacing, respectively.

**Supplemental Figure 4.**
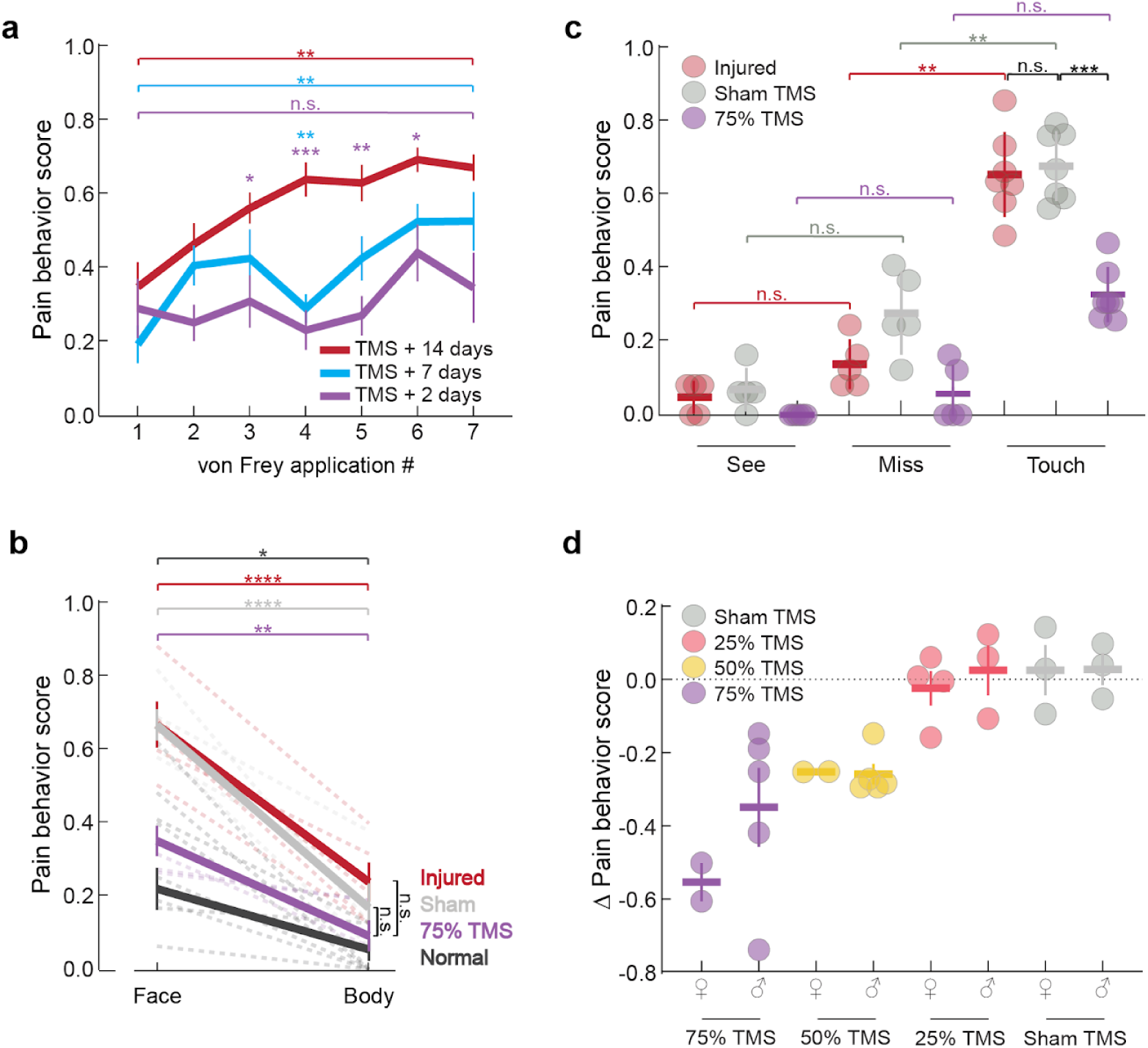
Pain scores were comparable across female and male mice after TMS treatment and were driven by the touches to the face with the von Frey filament. **(a)** Within individual testing sessions, pain sensitivity rose in response to multiple applications of a von Frey filament at 7 and 14 days but not 2 days after TMS treatment. Mean ± s.e.m. pain behavior scores in response to successive touches to the face with a von Frey filament during a single testing session, evaluated at 2 d (red), 7d (blue), or 14 d (purple) after TMS treatment at 50% or 75% of motor threshold (Repeated measures two-way ANOVA: von Frey applications #, p<0.0001; days after treatment, p<0.0001; application # × days post treatment interaction, p=0.084; n=15 mice). **(b)** TMS treatment led to reduced nocifensive behaviors in response to touches with a von Frey filament to the face but not to the body of the mouse. Plotted are mean ± s.e.m. pain behavior scores, evaluated after a touch to the injured face or body, at ∼7 days before injury (*Normal*), 21 days after injury (*Injured*), or 2 days after real (*75% TMS*) or sham (*Sham*) TMS treatment (n=5 mice per condition. Colors denote different treatment conditions. Black: Normal. Red: Injured. Gray: Sham TMS. Purple: 75% motor threshold TMS). (Repeated Measures two-way ANOVA; Condition or treatment (Normal, Injured, Sham TMS, 75% TMS): p<0.0001; Face *vs.* body: p<0.0001; Condition × face/body interaction: p=0.009; *p<0.05, **p<0.01, ****p<0.0001, Fisher’s least significant difference (LSD) test with a Holm-Bonferroni correction for multiple comparisons). **(c)** Nociceptive stimuli are the primary drivers of mouse nocifensive responses, rather than the sight of nearby or approaching von Frey filaments. Plotted are mean ± s.d. behavior scores of injured mice before (*Injured*) or after sham (*Sham TMS*) or real TMS treatment at 75% of motor threshold (*75% TMS*), evaluated at the time at which the von Frey filament first appeared in the mouse’s field of view (*See*), in cases when the experimenter missed the target region (*Miss*), or in response to a successful touch of the von Frey filament (*Touch*) (n=5-7 mice per group; repeated measures two-way ANOVA; p<0.0001; Fisher’s least significant difference (LSD) post hoc test with Holm-Bonferroni correction; **p<0.01, ***p<0.001 after adjustment for multiple comparisons). **(d)** Male and female mice had comparable treatment responses to TMS. Plotted are mean ± s.e.m. changes in the pain behavior scores of individual mice, evaluated over the period extending from ∼7 days before TMS treatment to 2 days afterward, as scored following a touch to the face with the von Frey filament. Data points show results from individual mice, following TMS treatment at the following percentages of motor threshold: *75% TMS* (purple, n=2 female and 5 male mice); *50% TMS* (yellow, n=2 female and 5 male mice); *25% TMS* (orange, n=4 female and 3 male mice); *Sham TMS* (light gray, n=3 female and 3 male mice). (Repeated measures two-way ANOVA; sex: p=0.27; TMS dose: p=0.001; sex × TMS dose interaction: p=0.46).

**Supplemental Figure 5.**
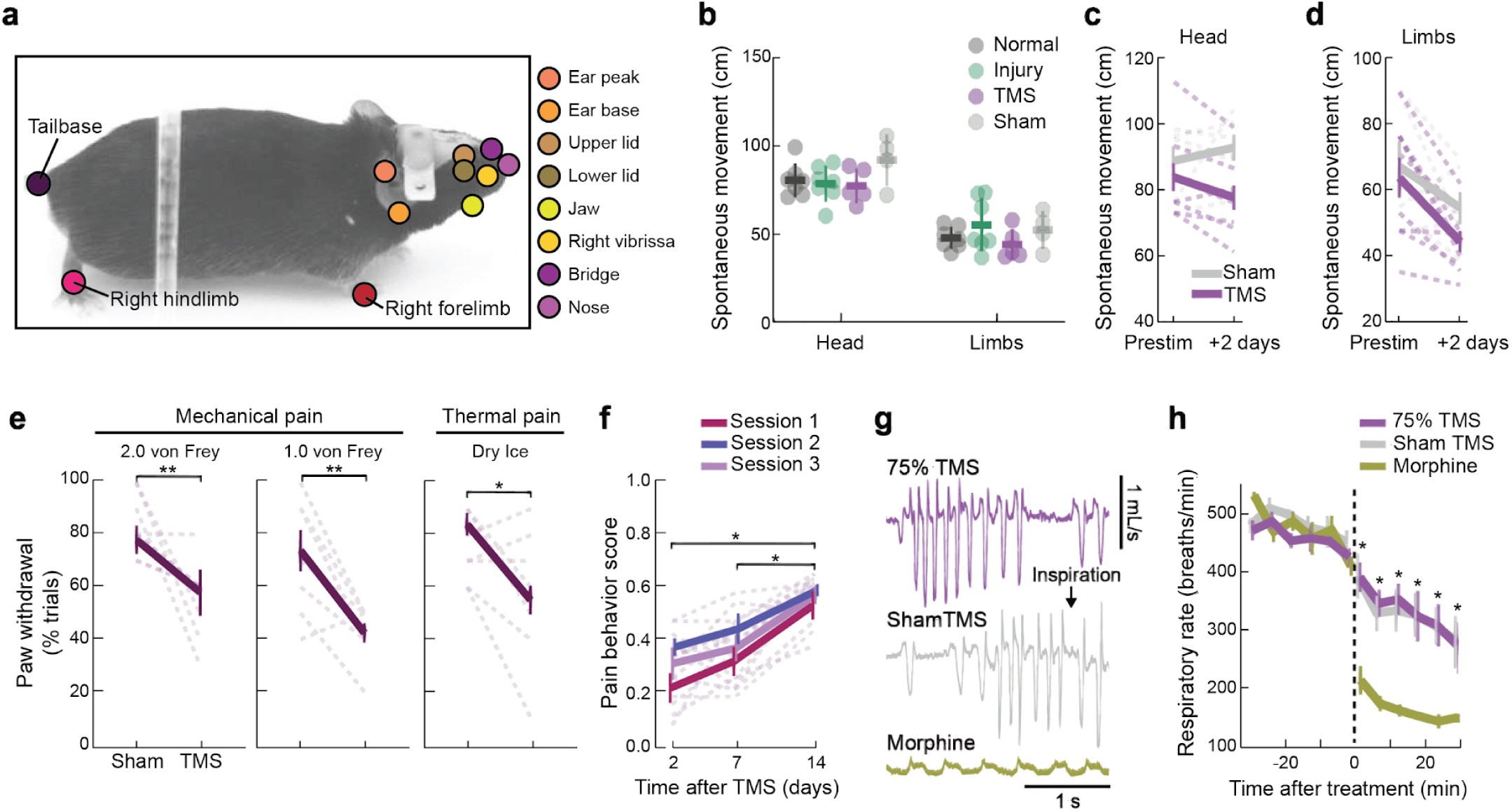
TMS reduced pain-associated behaviors without impairing spontaneous movement or respiratory rates. **(a)** Example video frame from a freely moving mouse, overlaid with DeepLabCut markers used to track normal or pain-related head movements (*e.g.,* reflexive withdrawal or grooming), as well as limb movements during locomotion. We tracked keypoints on all four limbs, the base and tip of the tail, and 11 points on the head (nose, jaw, bridge of the nose, left and right vibrissa pads, right and left ear peaks, and the top and bottom points of both the right and left eyelids). We interpolated these positional measurements when body parts were transiently out of view of the camera as the mouse explored its environment. Before each sensory testing session (Fig. 1e), we recorded the mouse’s behavior for 3 min. To quantify the total amount of head movement during this period, we computed the total mean distance traveled by the keypoints on the nose, nose bridge, vibrissa pads, ear peaks, and jaw. We estimated the mouse’s net locomotor distance traveled based on its limb displacements. **(b)** Levels of spontaneous head and limb movements were unaffected by TMS treatment at 75% of motor threshold. Plotted are mean ± s.d. distances traveled by the heads and limbs of the mice (n=4–7 mice per group), with each data point showing the mean result for a single mouse, averaged across 2 consecutive testing sessions. (*Normal*: average results for each mouse tested twice at least 7 days after headbar implantation but no more than two days prior to the nerve injury; *Injury*: average results for each mouse at 14 and 21 days after nerve injury; *real and sham TMS groups*: average results for each mouse at 2 and 7 days after treatment; ANOVA across conditions for head movements; p=0.23; ANOVA across conditions for limb movements; p=0.57). **(c, d)** TMS produced a subtle reduction in spontaneous head movements but not locomotor distance, suggesting TMS might selectively reduce spontaneous pain in addition to evoked pain. TMS-treated mice exhibited reduced spontaneous head movements, **(c)**, at 2 days post-treatment relative to pre-treatment baseline levels. Spontaneous locomotion, **(d)**, decreased similarly after treatment in mice that received real or sham TMS. Plotted are mean ± s.e.m. distances traveled; thin lines connect results for individual mice from pre-treatment to 2 days post-treatment. **(c)**, Repeated measures two-way ANOVA for head movements, Prestim *vs.* 2 days post-treatment: p=0.92; TMS *vs.* Sham: p=0.02; Interaction: p=0.31; Fisher’s LSD post hoc test with Holm-Bonferroni correction for multiple comparisons, *p<0.05. **(d)**, Repeated measures two-way ANOVA for limb movements, Prestim *vs.* 2 days post-treatment: p=0.001; TMS *vs.* Sham: p=0.33; Interaction: p=0.41). (n=4–7 mice per group). **(e)** TMS reduced acute pain sensitivity. We applied mechanical and cold nociceptive stimuli to an uninjured hindlimb 30 min after real or sham TMS treatment at 75% of motor threshold. Plotted are the mean ± s.e.m. (n=8 mice) percentage of trials in which mice withdrew from the painful stimulus (n=10 Trials per stimulus). Dashed lines: results from individual mice. (Wilcoxon signed-rank test for each stimulus; 2.0 g von Frey filament: p=0.008; 1.0 g filament: p=0.008; dry ice: p=0.04). **(f)** Mice did not desensitize to TMS over repeated administrations. Plotted are mean ± s.e.m. pain scores at days 2, 7, and 14 following each of three sessions of TMS treatment applied at 75% of motor threshold. Successive TMS sessions were 4 weeks apart. Dashed lines: results for each of n=5 individual mice. Repeated measures two-way ANOVA showed that pain scores varied with the time elapsed since TMS treatment (p<0.0001) but were statistically indistinguishable across the 3 sessions (p=0.2). (*p<0.05; Fisher’s LSD post hoc test with Holm-Bonferroni correction for multiple comparisons). **(g, h)** TMS, despite requiring MOR signaling for analgesia induction, did not cause respiratory depression. As deaths from opioid overdose result mainly from respiratory depression, we used whole-body plethysmography to check for possible respiratory side effects of TMS. Example airflow traces, (**g**), and mean ± s.d. baseline and post-treatment respiratory rates, (**h**), are shown for mice that received either sham or real (75% of motor threshold) TMS or 20 mg/kg morphine (a MOR agonist). The morphine group served as a positive control and received a dosage known to cause respiratory depression in mice. Vertical dashed line: time of treatment delivery. Repeated measures two-way ANOVA (n=6 mice per group; time: p<0.0001, treatment: p<0.001; Time × treatment interaction: p<0.0001) revealed significant effects of morphine compared to either TMS protocol. (*p<0.05 denotes time points at which the morphine group differed from the TMS groups using Fisher’s LSD post hoc test with a Holm-Bonferroni correction for multiple comparisons).

**Supplemental Figure 6.**
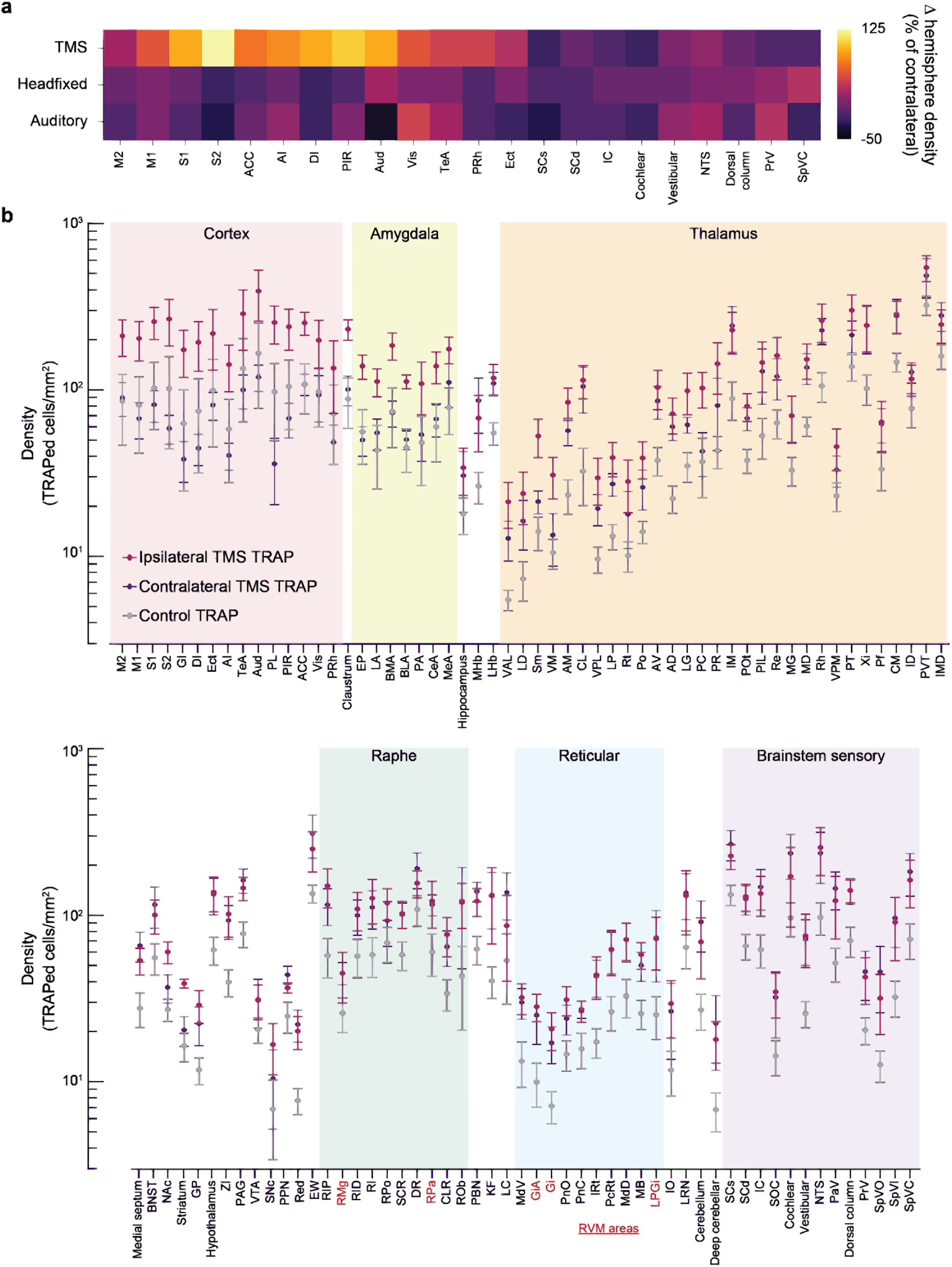
Genetic trapping studies show motor cortical TMS activates neurons ipsilaterally in the neocortex and amygdala and bilaterally in the brainstem. **(a)** Motor cortical TMS activates neurons in regions of the ipsilateral neocortex (*TMS*), whereas mere placement of the mouse into the TMS apparatus (*Headfixed*) or exposure of the animal to the sounds produced by the TMS pulses (*Auditory*) do not. The color plot shows, for individual brain areas in each of these three groups of mice, the percentage difference in the mean density of trapped neurons in the hemisphere ipsilateral to the site of TMS, relative to the mean value in the contralateral hemisphere. For the two control groups of mice that received no genuine TMS pulses, we assigned the right hemisphere to the ipsilateral group and the left hemisphere to the contralateral group, just as for the experimental group of mice. (Repeated measures two-way ANOVA; Group (TMS *vs.* Headfixed *vs.* Auditory): p=0.001; Brain areas: p=0.15; Group × Brain area Interaction: p=0.4; n=3 mice per group). **(b)** Expanded set of results supporting Fig. 2c, showing that neurons were trapped ipsilaterally to the site of stimulation in the neocortex and amygdala and bilaterally in thalamic and brainstem regions. Plotted are mean ± s.e.m. densities of genetically trapped neurons across 109 brain regions in n=5 ipsilateral and contralateral brain hemispheres of mice that received TMS (75% of motor threshold) and in n=10 hemispheres from control mice that were headfixed but did not receive TMS. **Abbreviations:** AD: Anterodorsal thalamus, AI: Agranular insular cortex, AM: Anteromedial thalamus, ACC: Anterior cingulate cortex, Aud: Auditory cortex, AV: Anteroventral thalamus, BLA: Basolateral amygdala, BMA: Basomedial amygdala, BNST: Bed nucleus of the stria terminalis, CeA: Central amygdala, CL: Centrolateral thalamus, CLR: Central linear raphe, CM: Centromedian thalamus, DI: Dysgranular insular cortex, DR: Dorsal raphe, Ect: Ectorhinal cortex, EP: Endopiriform, EW: Edinger-Westphal, GI: Granular insular cortex, Gi: Gigantocellular reticular formation, GiA: Gigantocellular reticular formation, alpha part, GP: Globus pallidus, IC: Inferior colliculus, ID: Interanterodorsal thalamus, IM: Interanteromedial thalamus, IMD: Intermediodorsal thalamus, IO: Inferior olive, IRt: Intermediate reticular formation, KF: Kölliker-Fuse, LA: Lateral amygdala, LC: Locus coeruleus, LD: Lateral dorsal thalamus, LHb: Lateral habenula, LG: Lateral geniculate thalamus, LP: Lateral posterior thalamus, LRN: Lateral reticular formation, LPGi: Lateral paragigantocellular reticular formation, M1: Primary motor cortex, M2: Secondary motor cortex, MB: Midbrain reticular formation, MD: Medial dorsal thalamus, MeA: Medial amygdala, MG: Medial geniculate thalamus, MHb: Medial habenula, MdD: Medullary reticular formation, dorsal, MdV: Medullary reticular formation, ventral, NAc: Nucleus accumbens, NTS: Nucleus of the solitary tract, PAG: Periaqueductal gray, PA: Posterior amygdala, PaV: Paratrigeminal, PC: Paracentral thalamus, PBN: Parabrachial, PcRt: Parvicellular reticular formation, Pf: Parafascicular thalamus, PIL: Posterior intralaminar thalamus, PIR: Piriform cortex, PL: Prelimbic cortex, PnC: Pontine reticular formation, caudal, PnO: Pontine reticular formation, oral, Po: Posterior thalamus, POt: Posterior triangular thalamus, PPN: Pedunculopontine, PRh: Perirhinal cortex, PT: Paratenial thalamus, PVT: Paraventricular thalamus, Re: Reuniens nucleus, RID: Interpeduncular raphe, RI: Interfascicular raphe, RIP: Interpositus raphe, RMg: Raphe magnus, ROb: Raphe obscurus, RPa: Raphe pallidus, RPo: Raphe pontis, Rh: Rhomboid thalamus, Rt: Reticular thalamus, S1: Primary somatosensory cortex, S2: Secondary somatosensory cortex, SCd: Superior colliculus, dorsal, SCs: Superior colliculus, superficial, SCR: Superior central raphe, Sm: Submedial thalamus, SNc: Substantia nigra compacta, SOC: Superior olivary complex, SpVC: Spinal trigeminal nucleus caudalis, SpVI: Spinal trigeminal nucleus interpolaris, SpVO: Spinal trigeminal nucleus oralis, TeA: Temporal association cortex, VAL: Ventral anterior-lateral thalamus, Vis: Visual cortex, VM: Ventral medial thalamus, VPL: Ventral posterolateral thalamus, VPM: Ventral posteromedial thalamus, VTA: Ventral tegmental area, Xi: Xiphoid thalamus, ZI: Zona incerta

**Supplemental Figure 7.**
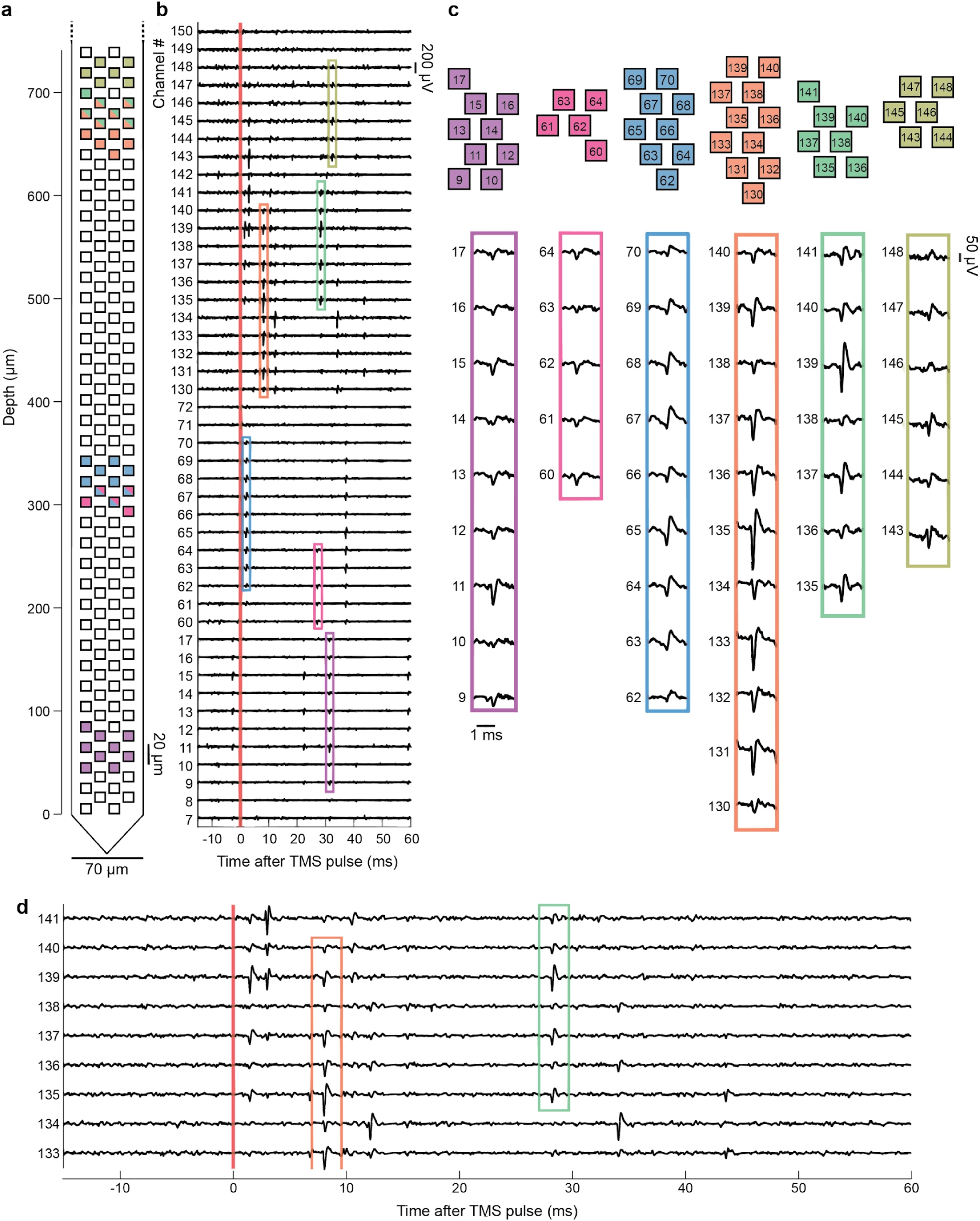
Spike waveforms of RVM neurons recorded with Neuropixels probes during TMS pulses. **(a, b)** Electrical activity observed on multiple Neuropixels recording channels after a single pulse of TMS (red vertical line). (**a**) Schematic of a Neuropixel probe. Recording sites on which the electrical dynamics of 6 different example RVM neurons (from the light blue-labeled probe in Fig. 4b) were observed are marked in 6 corresponding colors. (**b**) Band-pass filtered (300 Hz–10 kHz) electrical traces from 150 different recording channels reveal the responses of individual RVM neurons to a single TMS pulse. Auditory responses to the clicking sound of a TMS pulse appeared right after stimulation (2–7 ms after the pulse, as illustrated by the blue-labeled cell, likely relayed monosynaptically from the cochlear nuclei^148^), followed by activity driven by Layer 5 pyramidal tract neurons that project monosynaptically to the RVM (5–15 ms after the pulse, within the range of delays reported in past studies^149,150^, as illustrated by the orange-labeled cell). Subsequent waves of incoming excitation were likely polysynaptic and probably included a mix of auditory and TMS-driven signals (15 ms to >50 ms after the pulse, as illustrated by the pink-, purple-, green-, and olive-labeled cells). **(c)** Example traces of spike waveforms (*lower*) recorded across multiple channels of the Neuropixels probe (*upper*) for the same 6 color-corresponding neurons shown in **a** and **b**. The color associated with each set of spike waveforms conveys the location of the corresponding recording sites, as shown in **a**. **(d)** Illustrative subset of electrical traces from recording channels 133 to 141, shown on an expanded scale. Even though spike waveforms from different cells can be detected on overlapping sets of recording sites, spikes from the different neurons can still be differentiated based on their distinct footprints across the Neuropixels probe, as illustrated in panel **c**.

## Methods

### Mice

All animal procedures were approved by the Stanford University Administrative Panel on Laboratory Animal Care (APLAC) and the University of North Carolina Institutional Animal Care and Use Committee (IACUC) and abided by the recommendations of the International Association for the Study of Pain (IASP). We used 67 wildtype (C57BL/6) mice (Jax #000664; 30 females, 37 males), 6 *Oprm1^-/-^* mice (MOR KO, Jax#007559; 3 females, 3 males), 5 *Cux2^CreERT^*^2^ mice (L2/3 Pyramidal Cre, Allen Institute, 5 females), 5 *Rbp4^Cre^* mice (L5 Pyramidal Cre, Allen Institute, 3 females, 2 males), 5 *Sst^Cre^* (Somatostatin Cre, Jax #013044, 5 males), 5 *PV ^Cre^* (Parvalbumin Cre, Jax #008069, 5 males), 10 *Fos^2A-iCreER^*mice (TRAP2, Jax #030323, 7 females, 3 males), and 10 *Fos^2A-iCreER^/*Ai14 mice (TRAP2/Ai14, Jax #030323 × Jax #007914, 5 females, 5 males) of ages 6–100 weeks for this study. Mice resided in a temperature-controlled environment with a 12 h light/dark cycle and *ad libitum* access to food and water.

### Reagents

#### Drugs

We prepared clozapine-N-oxide (CNO, Tocris, #4936), naloxone hydrochloride (Sigma, #N7758), opiorphin (Santa Cruz, sc-253217), muscimol (Tocris, #2763-96-4), and D-AP5 (Tocris, #79055-68-8) in 0.9% sodium chloride (Hospira NDC 0409-4888-10). We prepared vehicle solutions and dilutions of the above compounds in 0.9% sodium chloride. For TRAPing procedures, we prepared 4-hydroxytamoxifen (4-OHT; Sigma, #H6278) in Kolliphor EL (Sigma, #27963).

#### Viruses

To express a chemogenetic actuator hM4Di in a Cre-dependent manner (**Figs. 3a–d,x,** and **5f,g**), we injected AAV2/8*-hSyn-*DIO*-hM4Di-mCherry* (University of North Carolina, titre: 6.4E12). To express ChR2 in M2 to RVM projection neurons (**Fig. 3s–u,w,x**), we injected AAV2/9*-CAG-*FLEx*-ChR2(H134R)-tdTomato* (Addgene, titre: 2.1E12) in cortical area M2 and AAV2-Retro*-Ef1a-Cre-WPRE* in RVM (Addgene, titre: 6.7E12). To trace axonal outputs of Cre+ neurons via expression of GFP and synaptophysin-mRuby (**Fig. 3e–r**), we injected AAVDJ*-hSyn-*DIO*-GFP-2A-synaptophysin-mRuby* (Stanford Viral Core, titre: 2.6E13) in M2.

#### Antibodies

To visualize the mCherry, GFP, and mRuby signal in the above studies (**Fig. 3c,f–r,t,u**), we performed immunohistochemical staining with the following primary antibodies: rabbit anti-GFP [Novus, NB600 (rabbit, 1:1000)] or chicken anti-RFP [Rockland, 600-901-379S (chicken, 1:1000)]. We then applied Alexa Fluor-conjugated secondary antibodies: donkey anti-rabbit 488 [Invitrogen, A21206 (1:1000)] or donkey anti-Chicken 594 [Jackson Immuno, 703-586-155 (1:1000)].

### Surgical procedures

We performed all surgeries under aseptic conditions using a digital small-animal stereotaxic instrument (David Kopf Instruments, Model 963). We initially anesthetized animals with 3-5% (v/v) isoflurane and maintained them at a surgical plane of anesthesia with 1-2% (v/v) isoflurane in oxygen. A heating pad (FHC, 40-90-8D) maintained body temperature at 37°C. We positioned mice in the stereotax, removed the hair above the skull, and opened the skin. Below are the detailed steps for each procedure.

#### Virus injection surgery

To stereotaxically inject viruses into the brain for chemogenetic modulation (**Figs. 3a–d,x, and 5f,g**), optogenetic manipulation (**Fig. 3s–u,w,x**), or viral tract tracing (**Fig. 3e–r**), we first leveled the skull. Then, we used a 0.5-mm-diameter drill bit to drill a burr hole(s) into the skull above the cortex or a 1.0-mm-diameter drill bit over the RVM (drill: Osada, EXL-M40; drill bits: Fine Science Tools, #19007). We performed all virus injections using a Nanoject III Nanoliter injector (Drummond, 3-000-207) and pulled glass capillaries (Drummond, 3-000-203-G/X). Injections into the RVM (see below) required long pulled tips (5–10 mm in length) to penetrate sufficiently deep into the brain, while cortical injections required shorter pulled tips (2–5 mm in length). We slowly lowered the nanoject pipette to the target coordinates (see next section). After injection, we slowly retracted the pipette and immediately performed a headbar surgery.

#### Viral injections for chemogenetic or optogenetic studies

To express hM4Di in M2 (**Fig. 3a–d**), we injected 100 nL of AAV2/8*-hSyn-*DIO*-hM4Di-mCherry* virus intracranially at three rostrocaudal coordinates (RC: +2.0, +1.5, +1.0 mm) along a single mediolateral axis (ML: 1.5 mm). At each RC, ML position, we injected at 5 distinct depths (DV: –0.9, –0.7, –0.5, –0.3, –0.1 mm from the brain surface). The total injected volume was 1.5 µL (0.5 µL/cortical location), distributed across ∼0.75 mm^3^ of tissue at a rate of 20 nL/s. To express channelrhodopsin in M2 neurons projecting to the RVM (**Fig. 3s–u,w,x**), we used the same injection anatomical coordinates in M2 as above, delivering AAV2/9-*CAG*-Flex-*ChR2(H134R)-tdTomato*. To ensure projection-specificity, we infused 200 nL of AAV2-Retro-*Ef1a-Cre-WPRE* in the RVM (RC: –5.2 mm, ML: 0 mm, DV: -5.0, -4.9, -4.8 from the surface of the brain or DV: +0.1, +0.2, +0.3 from the skull below the brain at 5 nL/s). To express hM4Di in the RVM (**Fig. 5f, g**), we intracranially injected 50 nL AAV2/8-*hSyn*-DIO-*hM4Di-mCherry* (RC: –5.2 mm, ML: 0 mm, DV: –5.0, –4.9, –4.8 mm from the surface of the brain or DV: +0.1, +0.2, +0.3 mm from the ventral surface of the skull with a total of 150 nL injected at 5 nL/s). For all RVM injections, we maintained the pipette at the final DV coordinate for 10-20 min after injection.

#### Viral injections for anterograde neural tracing

To trace neural outputs of TMS-activated neurons (**Fig. 3e–r**), we injected 100 nL AAVDJ*-hSyn-*DIO*-GFP-2A-synaptophysin-mRuby* in M2 at RC: +1.5 mm, ML: +1.5 mm, at 5 depths (DV: -0.9, -0.7, -0.5, -0.3, -0.1 mm from the brain surface) for a total of 500 nL injected at 50 nL/s.

#### Trigeminal neuropathic pain model

To induce a chronic constriction injury (CCI) of the left infraorbital branch of the trigeminal nerve, we placed mice in the stereotaxic apparatus with only the right ear bar stabilizing the head to ensure access to the left side of the face. We made a small skin incision (∼3 mm) between the vibrissa pad and the eye. We constricted the nerve with non-dissolvable silk sutures (Oasis, 5-0 MV-682-V) to ∼80% of its original size and confirmed by visual observation with the stereomicroscope that blood continued to flow normally. We then sutured the skin closed.

#### Headbar implantation for mice undergoing TMS

All mice that underwent TMS had a headbar implanted and fixed to their skull to stabilize the head during stimulation. After making an incision in the skin above the skull, we retracted the skin and secured it with ultraviolet (UV) curable glue (Loctite 4305). We delineated the skull sutures adjacent to the motor cortex with a marker for later TMS coil targeting. We then used a scalpel blade to score the visible skull plates before applying a thin layer of UV-curable glue over the open skull. We secured a headbar (0.8 cm x 0.5 cm, LaserAlliance, 18-24G thickness, stainless steel) over the posterior skull with an additional layer of UV-curable glue. Finally, we layered dental cement (#10000787, Fisher Scientific) over the UV glue.

Mice that received naloxone, muscimol, D-AP5, opiorphin, or vehicle intracranially underwent surgery to create a burr hole above the RVM (RC: –5.1 mm and ML: 0.0 mm) and secure a cannula. We used a 1.1-mm-diameter drill bit to create a burr hole over the RVM after which we gently positioned a stainless steel cannula (1.06 mm inner diameter, custom cut 18G McMaster’s 89935K66 to 3.7 mm length pieces at Stanford Varian Physics Machine Shop or ordered custom cut 304S/S Hypodermic Tubing 18G to 3.7 mm length pieces from Ziggy’s Tubes and Wires^77^) above the cerebellum. We then secured the cannula with UV curable glue and continued the surgery as described in the preceding paragraph.

#### Cranial window installation

To enable optogenetic stimulation of M2 neurons (**Fig. 3s–u, w, x**), we created a cranial window above the motor cortex. We removed the skin atop the cranium and mechanically removed all soft tissues from the skull surface with a scalpel. We then used a 0.7-mm-diameter micro drill burr (#19007-07, Fine Science Tools) to perform a craniotomy approximately 5-mm in diameter. After completing the drilling, we disconnected the circular-shaped bone piece from the surrounding skull, replaced it with a 5-mm diameter glass coverslip (Deckgläser Cover Glasses, 5mm ø), and pressed the window ∼100 µm downward to flatten the cortical tissue beneath it. We glued the window to the skull with UV-light-cured adhesive (#4305, Loctite). Finally, we installed a stainless steel headbar (described above) and filled the gap between the headplate and the skull with dental cement (#10000787, Fisher Scientific).

#### Craniotomy surgery for Neuropixels recordings

To enable Neuropixels electrophysiological recordings in head-fixed mice (**Fig. 4**), we used mice that had previously undergone chronic constriction of the infraorbital nerve (described above). In a second surgery, we secured a long headbar^100^ across the skull surface and performed a craniotomy over the intended recording target. We then used the same headbar surgery protocol described above, except here we positioned the headbar between the motor cortex and the RVM (*e.g.,* RC: -3.0 mm).

We acutely implanted the Neuropixels probe ∼20 minutes prior to initiating the recording, using a Sutter micromanipulator (MPC200 and ROE-200). For three of the eight Neuropixels recordings, we positioned the Neuropixels electrode above the dorsal surface of the cerebellum. For these mice, during surgery we made a rectangular craniotomy that was 2 mm ✕ 1 mm (RC ✕ ML) centered above RC: –5.2 and ML: 0.0. We next removed the dura by gently piercing the dura with an angled micro hook (Fine Science Tools, 10032-13) and then peeled off the dura with forceps or, if necessary, cut the dura with Vannas spring scissors (Fine Science Tools, 15001-08). Finally, we covered the brain with silicone (World Precision Instruments, Kwik-Cast).

For five of the eight recordings, the electrode approached the RVM from the dorsal surface of the caudal medulla, where it exits the skull^151^ (insertion site is ∼RC: –8.5, ML: 0.0, and DV: –4.25). For these surgeries, we made an incision along the back of the neck, gently separated the neck muscles along the midline, and removed the connective tissue at the foramen magnum (*i.e.,* between the skull and the dorsal surface of the medulla). Finally, we covered the exposed brain and spinal cord with silicone (World Precision Instruments, Kwik-Cast).

After surgery, we transferred the mouse to a recovery cage and placed it on a heating pad until it awoke. We then returned the mouse to its home cage and provided food and water on the cage floor without using a food hopper. Mice recovered for 2–7 d before the first recording session.

### Assaying nocifensive behaviors

We assayed mouse pain behaviors in response to innocuous touch across the normal baseline period, after chronic constriction injury, and after TMS. During testing, mice sat in a small, translucent chamber (5 cm ✕ 10 cm ✕ 20 cm). Each mouse had an initial habituation session in the chamber lasting 3 min; at three or four times in this interval, we slowly moved the 0.07 g von Frey filament toward their face. In subsequent sessions on later days, we used a FLIR camera (FL3-U3-13E4M-C; 60 fps; 672 × 656 pixels) to record each animal’s spontaneous behavior for the first 3 min they were in the chamber; we then captured a second video (1–2 min long) of their responses to innocuous touch. In all behavioral sessions (per timelines shown in **Figs. 1e, 3a, 5a)**, we applied the 0.07 g von Frey filament 4–7 times to the left vibrissa pad and peri-nasal region of the face. In some sessions, we alternatively touched the face and body (**Supplemental Fig. 4b**). To assay how the visual sight of the von Frey filament affected the mouse (**Supplemental Fig. 4c**), we evaluated the behavioral response the mouse had when the filament first entered the testing chamber. To assay miss trials (**Supplemental Fig. 4c**), we measured the pain behavior score in trials where the filament missed the face (typically ∼4-7 times per testing session).

We recorded videos of behavior and, after the experiment was completed, randomly watched (*i.e.,* across mice and sessions) and scored them. Playback occurred at ∼10% of the original frame acquisition rate; scorers exercised additional directional control of video frame advancement using the arrow keys to manually determine whether the withdrawal of the mouse from the filament was slow, moderate, heavy, or rapid (see scoring details below, **Supplemental Fig. 3b**). We recorded the frame of touch, the type of touch, the touch location on the body, and an observed response score for each face touch and, as necessary, for body touches, when the filament first entered the testing chamber, and on miss trials (**Supplemental Fig. 4b,c**). We averaged across all touches scored that day to create a composite score for that given day.

### Orofacial pain scoring system

We assigned the observed response score based on the scale detailed in **Supplemental Fig. 3b**, in which we used the velocity of withdrawal (slow or no response: <3 mm/s; moderate withdrawal: between 3 and 10 mm/s; heavy withdrawal: between 10 and 30 mm/s; rapid withdrawal: >30 mm/s) and additional affective-motivational behaviors assayed to determine the behavior score between 0 and 1 (with 1 being the most nocifensive). Besides withdrawal, mice would exhibit face wiping (single *vs.* multiple), grimacing (short-term: <2 s; prolonged: >5 s), hunching (noticeable back arch), and jumping (details for each numerical score between 0 and 1.00 are detailed in **Supplemental Fig. 3b**).

### DeepLabCut measurement of spontaneous behavior

To measure the spontaneous movement and pain-related movements of the head *vs.* the limbs of freely moving mice in video captured from the side (**Supplemental Fig. 5a–d**), we tracked 11 points on the head (nose, bridge of the nose, left and right vibrissa pads, right and left ear peaks, the top and bottom right and left eyelids, and the jaw), all four limbs, the tail base, and the tail tip, interpolating each point’s position when the body part was out of view (*e.g.,* when the mouse turned away from the camera). We recorded spontaneous behavior for three minutes before the start of each sensory testing session (aligned with the timeline in **Fig. 1e**). To determine total head movement during the spontaneous testing session, we calculated the mean displacement of the nose, bridge, vibrissa pads, ear peaks, and jaw (**Supplemental Fig. 5a–c**). To determine locomotor movement from the side-view recordings, we calculated the displacement of a body centroid estimated from the four limbs (**Supplemental Fig. 5a,b,d**). To identify changes before and after TMS, we compared each animal’s prestimulus spontaneous movement to its post-TMS or post-sham movement within the same animal (**Supplemental Fig. 5c,d**).

### Acute mechanical and thermal pain assays

To assess whether TMS or sham TMS altered acute nociceptive responses, we applied von Frey filaments and dry ice to the mice hindpaw 30 min post-stimulation (**Supplemental Fig. 5e**). We habituated mice to the testing chamber for 30 min on each of the two days prior to TMS. On the third day, we administered TMS or sham TMS, relocated them to the testing chamber, and assayed their sensory responses to mechanical (2.0 g and 1.0 g von Frey filaments) and thermal (packed dry ice in a 5 mL syringe with the tip cut off) stimuli 30 min later. We administered each stimulus-type ten times to the left hindpaw, with ∼15 s between each stimulus delivery. We applied the 2.0 g von Frey first, followed by the 1.0 g von Frey, and finally the dry ice. Paw withdrawal was measured as a brief lifting of the hindpaw in direct response to the filament or dry ice application and binary scored as # hindpaw withdrawals out of 10 applications. Two weeks later, they received the opposite treatment (TMS *vs.* sham) and were tested again.

### Plethysmography

To measure respiratory rate (**Supplemental Fig. 5g,h**), we used a Buxco Small Animal Whole Body Plethysmography (DSI). We placed adult wild-type mice in plethysmography chambers for 30 min to allow acclimation and obtain baseline respiratory rates. Afterward, we removed the mice and applied one of three treatments: 75% TMS, sham TMS, or an intraperitoneal injection of 20 mg/kg morphine. We then returned them to the chambers for another 30 minutes. Airflow voltage traces were exported and analyzed in MATLAB.

### Electromyography (EMG)

To perform acute EMG recordings (**Fig. 4c, Supplemental Fig. 1g, h**), we made electrodes from 50-µm-diameter insulated tungsten wire (AM Systems 795500). We stripped ∼1 mm from the recording tip, threaded it through a 30-gauge needle, and hooked the tip with forceps. On the recording day, we acutely implanted two wires in the vibrissa intrinsic muscle. We acquired EMG recordings using an A-M Systems amplifier (Model 1800, amplifier bandpass filtered 10–20,000 Hz). Signals were digitized using either a PowerLab 8/35 DAQ (AD Instruments, 40 kHz sampling rate) or a NI PXIe-6341, X Series DAQ (National Instruments, 30 kHz sampling rate).

**Supplemental Fig. 1g** shows raw EMG signals, whereas **Supplemental Fig. 1h** shows trial-averages of the rectified EMG envelope. In this case, we processed the raw signals to extract the envelope as follows: 1) fourth-order Butterworth bandpass filtering (100 Hz–500 Hz) using MATLAB *designfilt()* and *filtfilt()* functions, 2) rectification by taking the absolute value, and 3) second-order Butterworth low-pass filtering at 50 Hz using MATLAB *butter()* and *filtfilt()* functions.

For **Fig. 4c**, EMG acquisition was synchronized with the Neuropixels chassis (National Instruments, 30 kHz sampling rate). In this case, we processed the raw signals to extract the rectified EMG envelope as follows: 1) eighth-order Butterworth bandpass filtering (250 Hz–2.5 kHz) using MATLAB *designfilt()* and *filtfilt()* functions, 2) rectification by taking the absolute value, 3) second-order Butterworth low-pass filtering at 50 Hz using MATLAB *butter()* and *filtfilt()* functions, and 4) median filtering over a 2.5 ms window using the MATLAB *medfilt1()* function.

### Activity-dependent genetic TRAPing procedures

#### Genetic labeling of neurons activated by TMS

We habituated mice to head fixation for 15 min the day before TMS genetic TRAPing. On the day of TRAPing, we brought the experimental mice to the testing room, head-fixed them, and then gave them a 5 min TMS iTBS session at 75% motor threshold, followed by an injection of 50 mg/kg 4-OHT IP (**Figs. 2, 3e–r, Supplemental Fig. 6**). For head-fixed control mice (**Fig. 2, Supplemental Fig. 6**), we brought them to the testing room, head-fixed them, administered 50 mg/kg 4-OHT IP, and returned them to their home cage. Auditory control mice (**Supplemental Fig. 6a**) received the same procedures as the experimental mice, except the miniTMS coil was not activated. Instead, we activated a second miniTMS coil placed to the side of the animal so they would hear the same pattern but not receive the TMS treatment. We transcardially perfused TRAP2/Ai14 mice 7 days later and TRAP2 mice + virus 3 weeks later.

#### Genetic labeling of neurons disinhibited by morphine

To induce Cre in RVM OFF cells (**Fig. 5f, g**), TRAP2 mice with a prior injection of AAV2/8-*hSyn*-DIO-*hM4Di-mCherry* received an IP injection of 20 mg/kg morphine (West-Ward NDC 0641-6127-01, 10 mg/mL stock) followed immediately by an IP injection of 50 mg/kg 4-OHT.

### TMS device and coil construction

For TMS experiments, we used a solenoid comprising a 3-mm-diameter powder conical iron core sharpened to 2-mm at the tip (Micrometals material –52), surrounded by a triple layer of 0.4-mm-diameter copper wire (**Fig. 1a, Supplemental Fig. 1a**). The TMS electronic driver was designed to supply high current (up to ∼120 A) pulses to the electromagnets (**Supplemental Fig. 1b**). The current through the magnet was controlled by a high-power IGBT (30N135m, DIGIKEY) driven by a high-current IGBT/FET gate driver (TC4422, Microchip Technology). The TMS pulse timing was controlled by a high-speed, 32-bit microprocessor (ESP32, Espressif Systems) programmed to provide TMS intermittent theta bursts or single pulses. The microprocessor received timing instructions from a personal computer linked by Bluetooth. We measured the magnetic field intensity (**Fig. 1b,c** and **Supplemental Fig. 1c,d**) using a stepper motor-controlled micropositioning stage, which translated a small Hall probe (Bestol A1302) across the active regions of both the human and mouse TMS coils. We measured the induced EMF magnitude with a 2-mm-diameter current loop, similarly translated across the TMS coil’s active region. We measured the inductance of each miniTMS coil with a standard electronic inductance meter (**Supplemental Fig. 1d–f**).

### TMS treatment procedures

We first head-fixed the mice, positioned the coil approximately centered over the M2 region of motor cortex at ∼1.5 mm rostral and 1.5 mm lateral from bregma, and delivered ∼90 V stimulation while slightly adjusting coil position until a vibrissa twitch was observed. We then determined the motor threshold by gradually reducing the stimulation power in 5–10 V increments until TMS no longer evoked a twitch. We performed all TMS protocols subthreshold at 75%, 50%, or 25% of the motor threshold, as specified in the text and figures. We delivered TMS treatment using an iTBS pattern (**Fig. 1f**) for 5 min: a series of 3 pulses delivered 20 ms apart, each with the waveform shown in **Fig. 1b**, followed by a 160-ms-interval until the start of the next set of 3 pulses. This pattern repeated for 2 s, was followed by a 3-s-interval with no stimulation, and then began again. We used MATLAB to pseudorandomly assign mice to treatment groups and refined assignments to ensure balanced group composition. For sham TMS, we similarly head-fixed the mice but did not assess the motor threshold. Instead, we positioned the coil ∼5 cm above the skull and applied the 5-min iTBS protocol with the stimulation level set at 67 V, a value that we found to be the mean 75% motor threshold from our first batch of mice in **Fig. 1h,i**.

For IP naloxone + TMS (**Fig. 1j**), we administered 3 mg/kg naloxone (100 µL) 30 min before TMS. For intracranial drug administration + TMS (**Fig. 5a–d**), we first head-fixed the mouse, then lowered a syringe (Hamilton, 80430) into the RVM (DV: -4.5 to -5.3) through a cannula, and finally injected one of the following: naloxone, opiorphin, muscimol, and D-AP5 at 1, 5, 1, or 10 µg, respectively, in 200 nL of saline at a rate of 200 nL/min. After the infusion, we waited 5 min before removing the syringe and proceeding with our protocol for TMS administration.

### Measurement of TMS-induced muscle activity

To measure TMS-induced muscle activity, we recorded intrinsic vibrissa muscle EMG signals (see above) during 80 TMS pulses delivered via one of two miniTMS coils (**Supplemental Fig. 1g**). We aligned EMG responses to the onset of TMS and calculated the mean EMG response across trials (**Supplemental Fig. 1h**).

### Measurement of TMS dose-response curves

We determined TMS-evoked vibrissa movements from video recordings (FLIR Flea3 camera, FL3-U3-13E4M-C; 100 fps; 672 × 656 pixels) (**Fig. 1d, Supplemental Fig. 2d**). Video acquisition was synchronized with TMS output using the Video Capture add-on in the PowerLab8 data acquisition system and LabChart8 software. We processed videos using DeepLabCut^71^ software and temporally aligned the mouse’s body movement to the timing of TMS pulses using MATLAB software (Mathworks).

The location of the vibrissa pad center served as a reference for general vibrissa pad movement (**Fig. 1d, Supplemental Fig. 2a–c**). We tracked vibrissa angles by trimming all but the C3 whisker and then recording the angle of the C3 whisker relative to the head-nose axis (**Supplemental Fig. 2d**). We averaged the TMS-evoked movements over a 500-ms window across 40–50 trials for each TMS intensity. For sham TMS trials, we positioned the coil ∼5 cm above the mouse’s head, such that the mouse continued to hear the clicking noise emitted during TMS pulses, even though the magnetic field pulses delivered to the brain were greatly reduced.

### Optogenetic stimulation treatment procedures

We first head-fixed the mice, positioned the 470 nm LED (THOR labs, M470F4) approximately over the motor cortex, and delivered ∼15 mW stimulation while slightly adjusting the LED position until a vibrissa twitch was observed. We then determined the motor threshold by gradually reducing the stimulation power in 2-5 mW increments until stimulation no longer evoked a twitch. We performed optogenetic stimulation treatment (**Fig. 3w,x**) at 90% motor threshold (20 Hz for 5 min, 2 ms pulse width).

### Chemogenetic inhibition procedures

On the day of TMS, mice with expression of hM4Di in Cre+ neurons (**Figs. 3a–d,v** and **5f,g**) randomly received either CNO or saline vehicle injected IP (100 µL) one hour prior to a 75% motor threshold TMS treatment. Two weeks later, they underwent their second round of TMS with the opposite treatment (*i.e.,* all mice had one round with CNO and one with saline vehicle). We assayed mouse behavior responses two days after TMS (**Fig. 3a**).

### Neuropixels recordings

#### Identification of Neuropixels probe trajectories

Neuropixels probes were coated with 1% DiI to facilitate later identification of the recording site. Post hoc alignment was performed with the Neuropixels Trajectory Explorer^152,153^ together with images of coronal or horizontal brain sections (**Fig. 4b**). We advanced the recording probe to the RVM from either a dorsal or a posterior vantage, using the following methods.

#### Dorsal approach

We head-fixed the mice and removed the silicone from above the cerebellum. We inserted the probe at RC: –5.1 mm; ML: ∼0.0 mm; DV: 5–7 mm (relative to Bregma), perpendicular to the cerebellar surface. We waited 20 min for the probe to settle before starting recording.

#### Posterior approach

We first anesthetized mice with a cocktail of 100 mg/kg ketamine and 10 mg/kg xylazine. We inserted Neuropixels probes into the dorsal medulla at the foramen magnum (*e.g*., where the medulla and spinal cord exit the skull) at a 45° angle, to a depth of 4 mm. We waited at least 20 minutes after the mouse recovered from anesthesia before initiating the recording session.

#### Recording protocol

After initiating the recording session, we applied a series of 20-40 touches to the injured side of the face with 0.07-g or 2.0-g von Frey filaments, or pin prick (details below, **Fig. 4c–e**). For experiments examining the effect of the full 5-min treatment protocol (**Fig. 4f–j**), we first recorded baseline activity, performed a 5-min TMS iTBS protocol, and continued recording for at least 20 min. Mice in a sham group entered the TMS apparatus and were subject to the same protocol, with the exception that they received no TMS. At the end of each experiment, the mouse was transcardially perfused, and the brain was processed to confirm the recording location.

#### Recording and analysis

We acquired Neuropixels 1.0 data in SpikeGLX^154^. After completion of experiments, we sorted neural spikes using Kilosort 4.0^155^ and manually curated good cells in Phy^156^. We created data visualizations and analyzed the data with custom MATLAB code.

#### Identification of RVM ON and OFF neurons

We identified pain withdrawal events from videos aligned to EMG traces, then refined withdrawal onset based on EMG reflexive responses, time-locked to the peak EMG during withdrawal (video recorded at 60 frames/s; EMG at 30 kHz). Each recording session included 20–40 sequential painful stimuli: 15–20 applications of the 0.07 g von Frey filament, ∼10 applications of the 2.0 g von Frey filament, and ∼3 pinpricks. Unit firing rates during pain withdrawal (200 ms pre- to 500 ms post-withdrawal onset) were compared to shuffled non-pain time periods using Wilcoxon rank-sum tests. Units showing significant differences from shuffle (p<0.05) were classified as pain-modulated (**Fig. 4d,e** show trial-averaged responses across all modulated units from 8 mice).

ON units exhibited significantly increased firing relative to the 200-ms pre-withdrawal baseline (p < 0.05, Wilcoxon rank-sum); OFF units showed significant decreases. Units that did not significantly change from pre-withdrawal baseline (but remained significantly different from shuffle periods, indicating deviation from pre-pain-testing baseline firing) were classified as neutral

#### Analyses of TMS-induced changes in neural activity

We analyzed the activity patterns of ON and OFF neurons from 3 recordings during which the mouse received a 5 min TMS treatment (102 cells) and 3 recordings from a sham session in which we positioned the TMS coil and determined the motor threshold but did not deliver TMS to the mouse during the first portion of the experiment (153 cells). Note that the 3 mice in this sham group received the TMS protocol shown in **Supplemental Fig. 6a** after this sham period. We quantified the mean firing rate of each cell during a 5 min baseline period, the 5-min-period during which the mouse received TMS, and a 15–20 min period after the completion of TMS (**Fig. 4f–j**).

#### Pain cell activity index

The pain cell activity index, 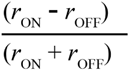, where *r*_ON_ and *r*_OFF_ denote the mean spike rates of ON and OFF cells, respectively. For each unit, we first calculated the mean spike rate during the baseline period (*r*_baseline_) and subtracted it from the spike rate during the TMS and post-TMS periods (*r*(t) - *r*_baseline_). We then calculated the average of the resulting baseline-corrected rates across ON and OFF cells to determine *r*_ON_ and *r*_OFF_ (**Fig. 4i,j**).

### Histological procedures

We deeply anesthetized mice and perfused them transcardially with 0.1 M phosphate-buffered saline (PBS, Gibco PBS, pH 7.4 10010023), followed by 4% formaldehyde (Thermo Fisher BP531-500). We dissected the brain, post-fixed it for 4–24 h at 4°C, and cryoprotected it overnight in a 30% sucrose solution in PBS (sucrose; Sigma, S0389). We cut 50-µm-thick sections from frozen brains using a Tanner Scientific TN50 Cryostat and stored them free-floating in PBS at 4°C.

#### Whole-brain analyses of genetically TRAPed cells

We mounted every fourth serial section from TRAP2;Ai14 brains on glass slides (Superfrost Plus Microscope slide: 12-550-15), then coverslipped them with Fluoromount-G (glass coverslips: Globe Scientific, 1419-20; mounting media: Southern Biotech, 0100-01). We acquired images of all sections on a Zeiss Axio Imager Z1 microscope (Texas red filter, Semrock, 2799) with a 10x objective (Zeiss, 420940-9901-000, numerical aperture 0.25). We exported images of individual brain sections as .png files (Zeiss, Zen).

We then aligned the images of brain sections to a 3-D volume with the QUINT pipeline in two parallel streams: one for histological alignment of the brain sections to the atlas and a second for identification of tdTomato+ cell bodies.

For CCF alignment, we converted images to .png format, scaled them to 25% resolution using the Nutil *Resize* function, and performed rigid registration in QuickNII^157,158^, followed by nonrigid alignment in VisuAlign^159^. For cell body identification, we separated the two hemispheres by duplicating each widefield image and blurring one hemisphere per copy to generate Left and Right hemisphere images for each section (custom MATLAB code).

We used the ilastik software to detect tdTomato14+ cell bodies in each image^94,160^. We first trained the ilastik *Pixel Classification* function on tdTomato fluorescence using a subset of images (3-6 sections out of ∼50 per brain), then batch-processed the remaining images to produce a pixel prediction map for each brain section. Next, we trained the ilastik *Object Classification [raw data + pixel prediction maps]* function to distinguish cell bodies from other fluorescent structures (*e.g.,* dendrites, axons, artifacts), producing binary object prediction images that we converted to the Glasbey look up table (LUT) in FIJI^161^.

We performed a subsequent alignment to the Allen Common Coordinate Framework (CCF) using the Nutil *Quantifier* function^91,93,162^, which provided 3D coordinates for each Ai14+ cell body, .csv reports with cell body counts, and images showing all cell bodies overlaid on their respective CCF sections. We generated the 3D rendering in **Fig. 2b** in MeshViewer^163^, a web-based viewer for registering histologically identified cell bodies or proteins to the CCF. We extracted the cell body counts from the .csv reports using custom Matlab scripts and performed statistics in GraphPad Prism.

#### Immunohistochemistry

To enhance visualization of virally-expressed GFP, mRuby, tdTomato, and mCherry fluorescence (**Fig. 3c,f–r,t,u**), we incubated coronal sections for 1 hr in 5% donkey serum in PBS containing 0.3% Triton X-100 (NDST; donkey serum: Sigma, S30; Triton X-100: Sigma, X100). Sections were then incubated overnight at room temperature in primary antibodies diluted in NDST. After three 10–15 min washes in PBS, sections were incubated for 2 h at room temperature in secondary antibodies diluted in NDST. Sections were then washed three additional times in PBS, mounted onto glass slides (Superfrost Plus Microscope Slides, 12-550-15), and coverslipped with Fluoromount-G (glass coverslips: Globe Scientific, 1419-20; mounting medium: Southern Biotech, 0100-01). We acquired images using an epifluorescence microscope (Zeiss Axio Imager Z1). One panel (**Fig. 3t**) was imaged on a Leica SP8 white light laser confocal microscope with a 20x objective (Leica, 11506343, 0.75 numerical aperture).

### Analyses of clinical data

To study a previously published clinical dataset (**Fig. 6**), we used data provided in table form within the publication^39^ (registered clinical trial #NCT02067273). To calculate changes in pain scores, quantified as ΔVAS values for patients who had 1 day of TMS treatment, we compared each patient’s baseline and post TMS pain scores. For patients who received 5 days of TMS treatment (who had two post TMS pain scores), we calculated ΔVAS between baseline and the minimum of the post TMS scores reported. We excluded two patients who withdrew from the study without follow-up, one patient taking the mixed MOR agonist/KOR antagonist buprenorphine, and two patients taking ketamine (a particularly weak MOR agonist with mixed effects).

### Statistical analyses

We assessed all datasets for normality in Graphpad Prism with a Shapiro-Wilk test. All subsequent analyses respectively involved nonparametric (*e.g.,* Kruskal-Wallis ANOVA) and parametric tests (*e.g.,* t-test, ANOVA) for non-normally and normally distributed datasets. After ANOVAs, we performed multiple comparisons using Fisher’s least significant difference (LSD) test, then adjusted the resulting p-values with the Holm-Bonferroni correction to control for the family-wise error rate (*i.e.,* the probability of making one or more false discoveries among all tested hypotheses when calculating multiple comparisons).

We performed statistical tests (**Figs. 1–3, 5, 6a**) for behavioral and anatomical experiments in GraphPad Prism 10.4.2. When needed, we used the MATLAB function *bonferroni_holm()*^164^ to perform Holm-Bonferroni corrections. We used MATLAB (R2024b) to analyze Neuropixels electrophysiological data (**Fig. 4**) and to perform mixed-effects regression and generalized linear models on the patient dataset (**Fig. 6b–g**).

## References

1. Rikard, S. M., Strahan, A. E., Schmit, K. M. & Guy, G. P., Jr. Chronic Pain Among Adults - United States, 2019-2021. MMWR Morb. Mortal. Wkly. Rep. 72, 379–385 (2023).

2. Dahlhamer, J. et al. Prevalence of chronic pain and high-impact chronic pain among adults - United States, 2016. MMWR Morb. Mortal. Wkly. Rep. 67, 1001–1006 (2018).

3. Zimmer, Z., Fraser, K., Grol-Prokopczyk, H. & Zajacova, A. A global study of pain prevalence across 52 countries: examining the role of country-level contextual factors. Pain 163, 1740–1750 (2022).

4. Mercer Lindsay, N., Chen, C., Gilam, G., Mackey, S. & Scherrer, G. Brain circuits for pain and its treatment. Sci Transl Med 13, eabj7360 (2021).

5. Matthes, H. W. et al. Loss of morphine-induced analgesia, reward effect and withdrawal symptoms in mice lacking the mu-opioid-receptor gene. Nature 383, 819–823 (1996).

6. Nadeau, S. E., Wu, J. K. & Lawhern, R. A. Opioids and chronic pain: An analytic review of the clinical evidence. Front. Pain Res. (Lausanne) 2, 721357 (2021).

7. Noble, M. et al. Long-term opioid management for chronic noncancer pain. Cochrane Database Syst. Rev. 2018, CD006605 (2010).

8. Jandhyala, R., Fullarton, J. R. & Bennett, M. I. Efficacy of rapid-onset oral fentanyl formulations vs. oral morphine for cancer-related breakthrough pain: a meta-analysis of comparative trials. J. Pain Symptom Manage. 46, 573–580 (2013).

9. Sinatra, R. S. & McQuay, H. Oral and Parenteral Opioid Analgesics for Acute Pain Management. in Acute Pain Management (eds. Sinatra, R. S., DeLeon-Cassasola, O. A., Viscusi, E. & Ginsberg, B.) 188–203 (Cambridge University Press, Cambridge, 2009).

10. Fields, H. State-dependent opioid control of pain. Nat Rev Neurosci 5, 565–575 (2004).

11. Basbaum, A. I., Bautista, D. M., Scherrer, G. & Julius, D. Cellular and molecular mechanisms of pain. Cell 139, 267–284 (2009).

12. Soliman, N. et al. Pharmacotherapy and non-invasive neuromodulation for neuropathic pain: a systematic review and meta-analysis. Lancet Neurol. 24, 413–428 (2025).

13. Attal, N. et al. Repetitive transcranial magnetic stimulation for neuropathic pain: a randomized multicentre sham-controlled trial. Brain 144, 3328–3339 (2021).

14. Khedr, E. M. et al. Longlasting antalgic effects of daily sessions of repetitive transcranial magnetic stimulation in central and peripheral neuropathic pain. J. Neurol. Neurosurg. Psychiatry 76, 833–838 (2005).

15. Galhardoni, R. et al. Repetitive transcranial magnetic stimulation in chronic pain: a review of the literature. Arch. Phys. Med. Rehabil. 96, S156–72 (2015).

16. Tsubokawa, T., Katayama, Y., Yamamoto, T., Hirayama, T. & Koyama, S. Treatment of thalamic pain by chronic motor cortex stimulation. Pacing Clin. Electrophysiol. 14, 131–134 (1991).

17. Mo, J.-J. et al. Motor cortex stimulation: a systematic literature-based analysis of effectiveness and case series experience. BMC Neurol. 19, 48 (2019).

18. Antal, A., Terney, D., Kühnl, S. & Paulus, W. Anodal transcranial direct current stimulation of the motor cortex ameliorates chronic pain and reduces short intracortical inhibition. J. Pain Symptom Manage. 39, 890–903 (2010).

19. André-Obadia, N. et al. Is Life better after motor cortex stimulation for pain control? Results at long-term and their prediction by preoperative rTMS. Pain Physician 17, 53–62 (2014).

20. Pirotte, B. et al. Comparison of functional MR imaging guidance to electrical cortical mapping for targeting selective motor cortex areas in neuropathic pain: a study based on intraoperative stereotactic navigation. AJNR Am. J. Neuroradiol. 26, 2256–2266 (2005).

21. Esfahani, D. R., Pisansky, M. T., Dafer, R. M. & Anderson, D. E. Motor cortex stimulation: functional magnetic resonance imaging–localized treatment for three sources of intractable facial pain: Report of 3 cases. J. Neurosurg. 114, 189–195 (2011).

22. André-Obadia, N. et al. Transcranial magnetic stimulation for pain control. Double-blind study of different frequencies against placebo, and correlation with motor cortex stimulation efficacy. Clin. Neurophysiol. 117, 1536–1544 (2006).

23. Hussein, A. E., Esfahani, D. R., Moisak, G. I., Rzaev, J. A. & Slavin, K. V. Motor Cortex Stimulation for Deafferentation Pain. Curr. Pain Headache Rep. 22, 45 (2018).

24. Annak, O. et al. Effects of continuous theta-burst stimulation of the primary motor and secondary somatosensory areas on the central processing and the perception of trigeminal nociceptive input in healthy volunteers. Pain 160, 172–186 (2019).

25. Brasil-Neto, J. P. Motor Cortex Stimulation for Pain Relief: Do Corollary Discharges Play a Role? Front. Hum. Neurosci. 10, 323 (2016).

26. Hazime, F. A. et al. Treating low back pain with combined cerebral and peripheral electrical stimulation: A randomized, double-blind, factorial clinical trial. Eur. J. Pain 21, 1132–1143 (2017).

27. Velasco, M. et al. Motor cortex stimulation in the treatment of deafferentation pain. I. Localization of the motor cortex. Stereotact. Funct. Neurosurg. 79, 146–167 (2002).

28. Meeker, T. J. et al. Non-invasive Motor Cortex Neuromodulation Reduces Secondary Hyperalgesia and Enhances Activation of the Descending Pain Modulatory Network. Front. Neurosci. 13, 467 (2019).

29. Hughes, S., Grimsey, S. & Strutton, P. H. Primary Motor Cortex Transcranial Direct Current Stimulation Modulates Temporal Summation of the Nociceptive Withdrawal Reflex in Healthy Subjects. Pain Med. 20, 1156–1165 (2019).

30. Lefaucheur, J.-P. et al. Neurogenic pain relief by repetitive transcranial magnetic cortical stimulation depends on the origin and the site of pain. J. Neurol. Neurosurg. Psychiatry 75, 612–616 (2004).

31. de Andrade, D. C., Mhalla, A., Adam, F., Texeira, M. J. & Bouhassira, D. Neuropharmacological basis of rTMS-induced analgesia: the role of endogenous opioids. Pain 152, 320–326 (2011).

32. Henssen, D. et al. A systematic review of the proposed mechanisms underpinning pain relief by primary motor cortex stimulation in animals. Neurosci. Lett. 719, 134489 (2020).

33. Cha, M., Um, S. W., Kwon, M., Nam, T. S. & Lee, B. H. Repetitive motor cortex stimulation reinforces the pain modulation circuits of peripheral neuropathic pain. Sci. Rep. 7, 7986 (2017).

34. Jiang, L. et al. Motor cortex stimulation suppresses cortical responses to noxious hindpaw stimulation after spinal cord lesion in rats. Brain Stimul. 7, 182–189 (2014).

35. Lopes, P. S. S., Campos, A. C. P., Fonoff, E. T., Britto, L. R. G. & Pagano, R. L. Motor cortex and pain control: exploring the descending relay analgesic pathways and spinal nociceptive neurons in healthy conscious rats. Behav. Brain Funct. 15, 5 (2019).

36. Kim, J. et al. Motor cortex stimulation and neuropathic pain: how does motor cortex stimulation affect pain-signaling pathways? J. Neurosurg. 124, 866–876 (2016).

37. Jung, H. H. et al. Rostral Agranular Insular Cortex Lesion with Motor Cortex Stimulation Enhances Pain Modulation Effect on Neuropathic Pain Model. Neural Plast. 2016, 3898924 (2016).

38. Pagano, R. L. et al. Transdural motor cortex stimulation reverses neuropathic pain in rats: a profile of neuronal activation. Eur. J. Pain 15, 268.e1–14 (2011).

39. Gaertner, M. et al. Advancing Transcranial Magnetic Stimulation Methods for Complex Regional Pain Syndrome: An Open-Label Study of Paired Theta Burst and High-Frequency Stimulation. Neuromodulation: Technology at the Neural Interface 21, 409–416 (2018).

40. Burchiel, K. J. Trigeminal neuropathic pain. Acta Neurochir. Suppl. 58, 145–149 (1993).

41. Haviv, Y., Zadik, Y., Sharav, Y. & Benoliel, R. Painful traumatic trigeminal neuropathy: an open study on the pharmacotherapeutic response to stepped treatment. J Oral Facial Pain Headache 28, 52–60 (2014).

42. Ding, W. et al. An Improved Rodent Model of Trigeminal Neuropathic Pain by Unilateral Chronic Constriction Injury of Distal Infraorbital Nerve. Journal of Pain (2017) doi:10.1016/j.jpain.2017.02.427.

43. Leung, Y. Y., Lee, T. C. P., Ho, S. M. Y. & Cheung, L. K. Trigeminal neurosensory deficit and patient reported outcome measures: the effect on life satisfaction and depression symptoms. PLoS One 8, e72891 (2013).

44. De Poortere, A., Van der Cruyssen, F. & Politis, C. The benefit of surgical management in post-traumatic trigeminal neuropathy: a retrospective analysis. Int. J. Oral Maxillofac. Surg. 50, 132–138 (2021).

45. Lima, M. C. & Fregni, F. Motor cortex stimulation for chronic pain: systematic review and meta-analysis of the literature. Neurology 70, 2329–2337 (2008).

46. George, M. S. Whither TMS: A one-trick pony or the beginning of a neuroscientific revolution? Am. J. Psychiatry 176, 904–910 (2019).

47. Klein, M. M. et al. Transcranial magnetic stimulation of the brain: guidelines for pain treatment research. Pain 156, 1601–1614 (2015).

48. Gutiérrez-Muto, A. M. et al. The complex landscape of TMS devices: A brief overview. PLoS One 18, e0292733 (2023).

49. Huerta, P. T. & Volpe, B. T. Transcranial magnetic stimulation, synaptic plasticity and network oscillations. J. Neuroeng. Rehabil. 6, 7 (2009).

50. Lioumis, P. et al. Optimization of TMS target engagement: current state and future perspectives. Front. Neurosci. 19, 1517228 (2025).

51. Bonmassar, G. et al. Microscopic magnetic stimulation of neural tissue. Nat. Commun. 3, 921 (2012).

52. Lee, S. W., Fallegger, F., Casse, B. D. F. & Fried, S. I. Implantable microcoils for intracortical magnetic stimulation. Sci. Adv. 2, e1600889 (2016).

53. Liu, L. et al. Design and evaluation of a rodent-specific focal transcranial magnetic stimulation coil with the custom shielding application in rats. Front. Neurosci. 17, 1129590 (2023).

54. Tang, A. D., et al. Construction and evaluation of rodent-specific rTMS coils. Front. Neural Circuits 10, (2016).

55. Boonzaier, J. et al. Design and evaluation of a rodent-specific transcranial magnetic stimulation coil: An in silico and in vivo validation study. Neuromodulation 23, 324–334 (2020).

56. Meng, Q. et al. A novel transcranial magnetic stimulator for focal stimulation of rodent brain. Brain Stimul. 11, 663–665 (2018).

57. Jiang, W., Isenhart, R., Liu, C. Y. & Song, D. A C-shaped miniaturized coil for transcranial magnetic stimulation in rodents. J. Neural Eng. 20, (2023).

58. Wang, F. et al. A sensory-motor-sensory circuit underlies antinociception ignited by primary motor cortex in mice. Neuron (2025) doi:10.1016/j.neuron.2025.03.027.

59. Khokhar, F. A., Voss, L. J., Steyn-Ross, D. A. & Wilson, M. T. Design and demonstration *in vitro* of a mouse-specific transcranial magnetic stimulation coil. IEEE Trans. Magn. 57, 1–11 (2021).

60. Heinricher, M. M. Opiates, rostral ventromedial medulla, and descending control. in Encyclopedia of Pain 2399–2405 (Springer Berlin Heidelberg, Berlin, Heidelberg, 2013).

61. Basbaum, A. I. & Fields, H. L. Endogenous pain control mechanisms: review and hypothesis. Ann Neurol 4, 451–462 (1978).

62. Basbaum, A. I. & Fields, H. L. Endogenous pain control systems: brainstem spinal pathways and endorphin circuitry. Annu. Rev. Neurosci. 7, 309–338 (1984).

63. Fields, H. L., Malick, A. & Burstein, R. Dorsal horn projection targets of ON and OFF cells in the rostral ventromedial medulla. J Neurophysiol 74, 1742–1759 (1995).

64. Fields, H. L., Vanegas, H., Hentall, I. D. & Zorman, G. Evidence that disinhibition of brain stem neurones contributes to morphine analgesia. Nature 306, 684–686 (1983).

65. François, A. et al. A Brainstem-Spinal Cord Inhibitory Circuit for Mechanical Pain Modulation by GABA and Enkephalins. Neuron 93, 822–839.e6 (2017).

66. Fields, H. L. & Basbaum, A. I. Brainstem control of spinal pain-transmission neurons. Annu Rev Physiol 40, 217–248 (1978).

67. Hallett, M. Transcranial magnetic stimulation: a primer. Neuron 55, 187–199 (2007).

68. Rothwell, J. C. et al. Motor cortex stimulation in intact man. 1. General characteristics of EMG responses in different muscles. Brain 110 (Pt 5), 1173–1190 (1987).

69. Day, B. L. et al. Motor cortex stimulation in intact man. 2. Multiple descending volleys. Brain 110 ( **Pt 5**), 1191–1209 (1987).

70. Volz, L. J., Hamada, M., Rothwell, J. C. & Grefkes, C. What makes the muscle twitch: Motor system connectivity and TMS-induced activity. Cereb. Cortex 25, 2346–2353 (2015).

71. Mathis, A. et al. DeepLabCut: markerless pose estimation of user-defined body parts with deep learning. Nat. Neurosci. 21, 1281–1289 (2018).

72. Vos, B. P., Strassman, A. M. & Maciewicz, R. J. Behavioral evidence of trigeminal neuropathic pain following chronic constriction injury to the rat’s infraorbital nerve. J. Neurosci. 14, 2708–2723 (1994).

73. Thibault, K., Rivière, S., Lenkei, Z., Férézou, I. & Pezet, S. Orofacial Neuropathic Pain Leads to a Hyporesponsive Barrel Cortex with Enhanced Structural Synaptic Plasticity. PLoS One 11, e0160786 (2016).

74. Deseure, K. & Hans, G. H. Chronic Constriction Injury of the Rat’s Infraorbital Nerve (IoN-CCI) to Study Trigeminal Neuropathic Pain. J. Vis. Exp. (2015) doi:10.3791/53167.

75. Deseure, K. & Hans, G. Orofacial neuropathic pain reduces spontaneous burrowing behavior in rats. Physiol. Behav. 191, 91–94 (2018).

76. Geha, P. Y. et al. Brain dynamics for perception of tactile allodynia (touch-induced pain) in postherpetic neuralgia. Pain 138, 641–656 (2008).

77. Corder, G. et al. An amygdalar neural ensemble that encodes the unpleasantness of pain. Science 363, 276–281 (2019).

78. Liu, S., Crawford, J. & Tao, F. Assessing orofacial pain behaviors in animal models: A review. Brain Sci. 13, 390 (2023).

79. Rodriguez, E. et al. A craniofacial-specific monosynaptic circuit enables heightened affective pain. Nat. Neurosci. 20, 1734–1743 (2017).

80. Kohútová, B., Fricová, J., Klírová, M., Novák, T. & Rokyta, R. Theta burst stimulation in the treatment of chronic orofacial pain: a randomized controlled trial. Physiol. Res. 66, 1041–1047 (2017).

81. Lefaucheur, J.-P. et al. Evidence-based guidelines on the therapeutic use of repetitive transcranial magnetic stimulation (rTMS): An update (2014–2018). Clin. Neurophysiol. 131, 474–528 (2020).

82. Chung, S. W., Hoy, K. E. & Fitzgerald, P. B. Theta-burst stimulation: a new form of TMS treatment for depression? Depress. Anxiety 32, 182–192 (2015).

83. Chung, S. W., Hill, A. T., Rogasch, N. C., Hoy, K. E. & Fitzgerald, P. B. Use of theta-burst stimulation in changing excitability of motor cortex: A systematic review and meta-analysis. Neurosci. Biobehav. Rev. 63, 43–64 (2016).

84. Borckardt, J. et al. A randomized, controlled investigation of motor cortex transcranial magnetic stimulation (TMS) effects on quantitative sensory measures in healthy adults: Evaluation of TMS device parameters. Clin. J. Pain 27, 486–494 (2011).

85. Paul, A. K., et al. Opioid Analgesia and Opioid-Induced Adverse Effects: A Review. Pharmaceuticals 14, (2021).

86. Bachmutsky, I., Wei, X. P., Kish, E. & Yackle, K. Opioids depress breathing through two small brainstem sites. Elife 9, (2020).

87. Liu, S. et al. Neural basis of opioid-induced respiratory depression and its rescue. Proc. Natl. Acad. Sci. U. S. A. 118, (2021).

88. Quillinan, N., Lau, E. K., Virk, M., von Zastrow, M. & Williams, J. T. Recovery from mu-opioid receptor desensitization after chronic treatment with morphine and methadone. J. Neurosci. 31, 4434–4443 (2011).

89. Baldo, B. A. Opioid-induced respiratory depression: clinical aspects and pathophysiology of the respiratory network effects. Am. J. Physiol. Lung Cell. Mol. Physiol. 328, L267–L289 (2025).

90. Allen, W. E. et al. Thirst-associated preoptic neurons encode an aversive motivational drive. Science 357, 1149–1155 (2017).

91. Yates, S. C. et al. QUINT: Workflow for Quantification and Spatial Analysis of Features in Histological Images From Rodent Brain. Front. Neuroinform. 13, 75 (2019).

92. Wang, Q. et al. The Allen Mouse Brain Common Coordinate Framework: A 3D Reference Atlas. Cell 181, 936–953.e20 (2020).

93. Groeneboom, N. E., Yates, S. C., Puchades, M. A. & Bjaalie, J. G. Nutil: A Pre- and Post-processing Toolbox for Histological Rodent Brain Section Images. Front. Neuroinform. 14, 37 (2020).

94. Berg, S. et al. Ilastik: interactive machine learning for (bio) image analysis. Nat. Methods 16, 1226–1232 (2019).

95. Deng, Z.-D., Lisanby, S. H. & Peterchev, A. V. Electric field depth-focality tradeoff in transcranial magnetic stimulation: simulation comparison of 50 coil designs. Brain Stimul. 6, 1–13 (2013).

96. Fields, H. L., Levine, J. D. & Basbaum, A. I. Descending control of pain transmission. in Sensory Functions 143–149 (Elsevier, 1981).

97. Bagley, E. E. & Ingram, S. L. Endogenous opioid peptides in the descending pain modulatory circuit. Neuropharmacology 173, 108131 (2020).

98. Roth, B. L. DREADDs for Neuroscientists. Neuron Preprint at 10.1016/j.neuron.2016.01.040 (2016).

99. McElvain, L. E. et al. Specific populations of basal ganglia output neurons target distinct brain stem areas while collateralizing throughout the diencephalon. Neuron 109, 1721–1738.e4 (2021).

100. Mercer Lindsay, N., et al. Orofacial Movements Involve Parallel Corticobulbar Projections from Motor Cortex to Trigeminal Premotor Nuclei. Neuron 104, 765–780.e3 (2019).

101. Kuramoto, E. et al. Individual mediodorsal thalamic neurons project to multiple areas of the rat prefrontal cortex: A single neuron-tracing study using virus vectors. J. Comp. Neurol. 525, 166–185 (2017).

102. Jeong, M. et al. Comparative three-dimensional connectome map of motor cortical projections in the mouse brain. Sci. Rep. 6, 1–14 (2016).

103. De Preter, C. C. & Heinricher, M. M. The ‘in’s and out’s’ of descending pain modulation from the rostral ventromedial medulla. Trends Neurosci. 47, 447–460 (2024).

104. Jun, J. J. et al. Fully integrated silicon probes for high-density recording of neural activity. Nature 551, 232–236 (2017).

105. Lüscher, C. & Malenka, R. C. NMDA receptor-dependent long-term potentiation and long-term depression (LTP/LTD). Cold Spring Harb. Perspect. Biol. 4, a005710–a005710 (2012).

106. Barbaro, N. M., Heinricher, M. M. & Fields, H. L. Putative pain modulating neurons in the rostral ventral medulla: reflex-related activity predicts effects of morphine. Brain Res. 366, 203–210 (1986).

107. Fatt, M. P. et al. Morphine-responsive neurons that regulate mechanical antinociception. Science 385, eado6593 (2024).

108. Bodnar, R. & Heinricher, M. M. Central mechanisms of pain suppression: Central mechanisms of pain modulation. in Neuroscience in the 21st Century 3861–3886 (Springer International Publishing, Cham, 2022).

109. de Andrade, E. M. et al. Neurochemical effects of motor cortex stimulation in the periaqueductal gray during neuropathic pain. J. Neurosurg. 132, 239–251 (2019).

110. BRAIN Initiative Cell Census Network (BICCN). A multimodal cell census and atlas of the mammalian primary motor cortex. Nature 598, 86–102 (2021).

111. DosSantos, M. F. et al. Building up analgesia in humans via the endogenous μ-opioid system by combining placebo and active tDCS: a preliminary report. PLoS One 9, e102350 (2014).

112. Fields, H. L., Basbaum, A. I. & Heinricher, M. M. Central nervous system mechanisms of pain modulation. in Wall and Melzack’s Textbook of Pain 125–142 (Elsevier, 2006).

113. Mayer, D. J. & Price, D. D. Central nervous system mechanisms of analgesia. Pain 2, 379–404 (1976).

114. Hosobuchi, Y. The current status of analgesic brain stimulation. Acta Neurochir. Suppl. (Wien) 30, 219–227 (1980).

115. Watkins, L. R. et al. Neurocircuitry of conditioned inhibition of analgesia: effects of amygdala, dorsal raphe, ventral medullary, and spinal cord lesions on antianalgesia in the rat. Behav. Neurosci. 112, 360–378 (1998).

116. Yamada, K. & Nabeshima, T. Stress-induced behavioral responses and multiple opioid systems in the brain. Behav. Brain Res. 67, 133–145 (1995).

117. Bannister, K. & Dickenson, A. H. The plasticity of descending controls in pain: translational probing. J. Physiol. 595, 4159–4166 (2017).

118. Guan, Y., Terayama, R., Dubner, R. & Ren, K. Plasticity in excitatory amino acid receptor-mediated descending pain modulation after inflammation. J. Pharmacol. Exp. Ther. 300, 513–520 (2002).

119. Chen, Q. & Heinricher, M. M. Descending Control Mechanisms and Chronic Pain. Curr. Rheumatol. Rep. 21, 13 (2019).

120. Zhou, H.-Y., Chen, S.-R., Chen, H. & Pan, H.-L. Opioid-induced long-term potentiation in the spinal cord is a presynaptic event. J. Neurosci. 30, 4460–4466 (2010).

121. Dacher, M. & Nugent, F. S. Morphine-induced modulation of LTD at GABAergic synapses in the ventral tegmental area. Neuropharmacology 61, 1166–1171 (2011).

122. Mussetto, V. et al. Opioids induce bidirectional synaptic plasticity in a brainstem pain center in the rat. J. Pain 24, 1664–1680 (2023).

123. Dacher, M. & Nugent, F. S. Opiates and plasticity. Neuropharmacology 61, 1088–1096 (2011).

124. St. Laurent, R., Martinez Damonte, V., Tsuda, A. C. & Kauer, J. A. Periaqueductal Gray and Rostromedial Tegmental Inhibitory Afferents to VTA Have Distinct Synaptic Plasticity and Opiate Sensitivity. Neuron 106, 624–636.e4 (2020).

125. Li, X.-H., Miao, H.-H. & Zhuo, M. NMDA receptor dependent long-term potentiation in chronic pain. Neurochem. Res. 44, 531–538 (2019).

126. Laumet, G., Chen, S.-R. & Pan, H.-L. NMDA receptors and signaling in chronic neuropathic pain. in The NMDA Receptors 103–119 (Springer International Publishing, Cham, 2017).

127. Petrenko, A. B., Yamakura, T., Baba, H. & Shimoji, K. The role of N-methyl-D-aspartate (NMDA) receptors in pain: a review. Anesth. Analg. 97, 1108–1116 (2003).

128. Duong, S., Bravo, D., Todd, K. J., Finlayson, R. J. & Tran, D. Q. Treatment of complex regional pain syndrome: an updated systematic review and narrative synthesis. Can. J. Anaesth. 65, 658–684 (2018).

129. Moore, T. J., Alami, A., Alexander, G. C. & Mattison, D. R. Safety and effectiveness of NMDA receptor antagonists for depression: A multidisciplinary review. Pharmacotherapy 42, 567–579 (2022).

130. Ramsay, D. S. et al. Nitrous oxide analgesia in humans: acute and chronic tolerance. Pain 114, 19–28 (2005).

131. Beaupain, M. C. et al. NMDA receptor involvement in dopaminergic modulation of neuroplasticity induced by paired associative stimulation. Int. J. Neuropsychopharmacol. 28, (2025).

132. Grewe, B. F. et al. Neural ensemble dynamics underlying a long-term associative memory. Nature 543, 670–675 (2017).

133. Fadok, J. P., Dickerson, T. M. K. & Palmiter, R. D. Dopamine is necessary for cue-dependent fear conditioning. J Neurosci 29, 11089–11097 (2009).

134. Johansen, J. P. et al. Hebbian and neuromodulatory mechanisms interact to trigger associative memory formation. Proc Natl Acad Sci U S A 111, E5584–92 (2014).

135. Downar, J., Siddiqi, S. H., Mitra, A., Williams, N. & Liston, C. Mechanisms of action of TMS in the treatment of depression. Curr. Top. Behav. Neurosci. 66, 233–277 (2024).

136. Antonelli, M. et al. Transcranial Magnetic Stimulation: A review about its efficacy in the treatment of alcohol, tobacco and cocaine addiction. Addict. Behav. 114, 106760 (2021).

137. Neggers, S. F. W., Petrov, P. I., Mandija, S., Sommer, I. E. C. & van den Berg, N. A. T. Understanding the biophysical effects of transcranial magnetic stimulation on brain tissue: the bridge between brain stimulation and cognition. Prog. Brain Res. 222, 229–259 (2015).

138. Epstein, C. M., Wassermann, E. M. & Ziemann, U. Physics and Biophysics of TMS. Oxford Handbook of Transcranial Stimulation (2008).

139. Klomjai, W., Katz, R. & Lackmy-Vallée, A. Basic principles of transcranial magnetic stimulation (TMS) and repetitive TMS (rTMS). Ann. Phys. Rehabil. Med. 58, 208–213 (2015).

140. Aberra, A. S., Wang, B., Grill, W. M. & Peterchev, A. V. Simulation of transcranial magnetic stimulation in head model with morphologically-realistic cortical neurons. Brain Stimul. 13, 175–189 (2020).

141. Seo, H., Schaworonkow, N., Jun, S. C. & Triesch, J. A multi-scale computational model of the effects of TMS on motor cortex. F1000Res. 5, 1945 (2016).

142. Pashut, T. et al. Mechanisms of magnetic stimulation of central nervous system neurons. PLoS Comput. Biol. 7, e1002022 (2011).

143. Siebner, H. R. et al. Transcranial magnetic stimulation of the brain: What is stimulated? - A consensus and critical position paper. Clin. Neurophysiol. 140, 59–97 (2022).

144. Terao, Y. & Ugawa, Y. Basic mechanisms of TMS. J. Clin. Neurophysiol. 19, 322–343 (2002).

145. Ribeiro, F. C. P., Brasil, J. S., Vianna, D. C., Pereira, K. F. & Fregni, F. The synergistic effects of cycloserine and anodal tDCS: a systematic review and meta-analysis. Exp. Brain Res. 243, 94 (2025).

146. Antal, A. & Paulus, W. A case of refractory orofacial pain treated by transcranial direct current stimulation applied over hand motor area in combination with NMDA agonist drug intake. Brain Stimul. 4, 117–121 (2011).

147. Wisner, A. et al. Human Opiorphin, a natural antinociceptive modulator of opioid-dependent pathways. Proc Natl Acad Sci U S A 103, 17979–17984 (2006).

148. Bellintani-Guardia, B., Schweizer, M. & Herbert, H. Analysis of projections from the cochlear nucleus to the lateral paragigantocellular reticular nucleus in the rat. Cell Tissue Res. 283, 493–505 (1996).

149. Liew, Y. J. et al. Inferring thalamocortical monosynaptic connectivity in vivo. J. Neurophysiol. 125, 2408–2431 (2021).

150. Oberle, H. M., Ford, A. N., Czarny, J. E., Rogalla, M. M. & Apostolides, P. F. Recurrent circuits amplify corticofugal signals and drive feedforward inhibition in the inferior colliculus. J. Neurosci. 43, 5642–5655 (2023).

151. Paxinos, G. & Franklin, K. B. J. Paxinos and Franklin’s the Mouse Brain in Stereotaxic Coordinates. (Academic Press, 2019).

152. Peters, A. Neuropixels Trajectory Explorer. https://github.com/petersaj/neuropixels_trajectory_explorer.

153. Peters, A. Neuropixels_trajectory_explorer: Neuropixels Trajectory Explorer with the Allen CCF Mouse Atlas. (Github).

154. SpikeGLX: SpikeGLX Recording System GUI [Neuropixels NI]. (Github).

155. Pachitariu, M., Sridhar, S., Pennington, J. & Stringer, C. Spike sorting with Kilosort4. Nat. Methods (2024) doi:10.1038/s41592-024-02232-7.

156. Cyrille Rossant, International Brain Laboratory, Cortex Lab (UCL), Alessio Buccino, Michael Economo, Cedric Gestes, Dan Goodman, Max Hunter, Shabnam Kadir, Christopher Nolan, Martin Spacek, Nick Steinmetz. Phy github. https://github.com/cortex-lab/phy.

157. Puchades, M. A., Csucs, G., Ledergerber, D., Leergaard, T. B. & Bjaalie, J. G. Spatial registration of serial microscopic brain images to three-dimensional reference atlases with the QuickNII tool. PLoS One 14, e0216796 (2019).

158. NITRC: QuickNII - Serial section aligner to volumetric atlases: Tool/Resource Info. https://www.nitrc.org/projects/quicknii.

159. NITRC: VisuAlign - Nonlinear adjustments after QuickNII: Tool/Resource Info. https://www.nitrc.org/projects/visualign/.

160. ilastik - ilastik. https://www.ilastik.org/.

161. Goldstein, J. I. et al. ImageJ and Fiji. in Scanning Electron Microscopy and X-Ray Microanalysis 187–193 (Springer New York, New York, NY, 2018).

162. NITRC: Nutil - Neuroimaging utilities: Tool/Resource Info. https://www.nitrc.org/projects/nutil/.

163. NXTView v0.8. https://www.nesys.uio.no/MeshView/meshview.html?atlas=ABA_mouse_v3.

164. bonferroni_holm - File Exchange - MATLAB CentralFile Exchange - MATLAB Central. https://www.mathworks.com/matlabcentral/fileexchange/69817-bonferroni_holm), (2019).

